# Paleolithic Gene Duplications Primed Adaptive Evolution of Human Amylase Locus Upon Agriculture

**DOI:** 10.1101/2023.11.27.568916

**Authors:** Feyza Yılmaz, Charikleia Karageorgiou, Kwondo Kim, Petar Pajic, Kendra Scheer, Human Genome Structural Variation Consortium, Christine R. Beck, Ann-Marie Torregrossa, Charles Lee, Omer Gokcumen

**Affiliations:** The Jackson Laboratory for Genomic Medicine, Farmington, CT, 06032, USA; Department of Biological Sciences, University at Buffalo, Buffalo, NY, 14260, USA; University of Connecticut, Institute for Systems Genomics, Storrs, CT, 06269, USA; The University of Connecticut Health Center, Farmington, CT, 06032, USA; Department of Psychology, University at Buffalo, Buffalo, NY 14260, USA; University at Buffalo Center for Ingestive Behavior Research, University at Buffalo, University at Buffalo, NY 14260, USA

**Author notes:** Contributed equally (Feyza Yılmaz, Charikleia Karageorgiou, Kwondo Kim).

## Abstract

Starch digestion is a cornerstone of human nutrition. The amylase genes code for the starch-digesting amylase enzyme. Previous studies suggested that the salivary amylase (*AMY1*) gene copy number increased in response to agricultural diets. However, the lack of nucleotide resolution of the amylase locus hindered detailed evolutionary analyses. Here, we have resolved this locus at nucleotide resolution in 98 present-day humans and identified 30 distinct haplotypes, revealing that the coding sequences of all amylase gene copies are evolving under negative selection. The phylogenetic reconstruction suggested that haplotypes with three *AMY1* gene copies, prevalent across all continents and constituting about 70% of observed haplotypes, originated before the out-of-Africa migrations of ancestral modern humans. Using thousands of unique 25 base pair sequences across the amylase locus, we showed that additional *AMY1* gene copies existed in the genomes of four archaic hominin genomes, indicating that the initial duplication of this locus may have occurred as far back 800,000 years ago. We similarly analyzed 73 ancient human genomes dating from 300 - 45,000 years ago and found that the *AMY1* copy number variation observed today existed long before the advent of agriculture (∼10,000 years ago), predisposing this locus to adaptive increase in the frequency of higher amylase copy number with the spread of agriculture. Mechanistically, the common three-copy haplotypes seeded non-allelic homologous recombination events that appear to be occurring at one of the fastest rates seen for tandem repeats in the human genome. Our study provides a comprehensive population-level understanding of the genomic structure of the amylase locus, identifying the mechanisms and evolutionary history underlying its duplication and copy number variability in relation to the onset of agriculture.

## Results

Copy number variation at the amylase locus is frequently attributed to human health and adaptation (*1*). As such, this structurally variable locus is a prime target for research on the fundamental biology of gene duplications. There are two types of amylase genes, *AMY1* and *AMY2,* which are reported to be expressed in the salivary glands and pancreas, respectively (*2*). Both genes encode for the amylase enzyme, which breaks down polymeric starch into simple sugar molecules, a crucial digestive process for starch-eating species (*3*). It has been shown that mammals that consume starch-rich diets underwent independent bursts of amylase gene duplications from the ancestral pancreatic *AMY2-like* gene (*4*). A great ape-specific duplication resulted in the formation of the salivary *AMY1* gene (*5*), which has since evolved to produce unusual copy number variations, ranging from 2 to 17 copies per diploid cell (*1*, *6*). This variation is especially prominent in human populations with high starch consumption, particularly those with a history of agriculture (*1*, *6*, *7*). These evolutionary insights indicate that copy number variation at the amylase locus may play an adaptive role in shaping the metabolic response to starchy diets, including the presence of microbes that break down amylase-resistant starch (*8*).

Given its adaptive and putative functional roles, the duplications of the *AMY1* gene was linked to the advent of agriculture approximately 10,000 years ago (*1*). However, the evolutionary analyses of the *AMY1* gene is hindered by the presence of highly similar DNA sequences (> 99%) within the amylase locus and makes it challenging to accurately resolve the haplotypes at this locus. The lack of precise nucleotide-level evolutionary analysis leads to disagreements about the timing and functional significance of *AMY1* gene duplications in relation to starch-rich diets and human evolution (*7*, *9–12*). To address this issue, we have resolved this locus at nucleotide resolution at a population level across 98 individuals from different populations, using optical genome mapping and long-read sequencing techniques. The nucleotide-resolved haplotypes of this locus subsequently allowed us to conduct evolutionary genetic analyses on ancient human and archaic hominin genomes to investigate the timing of *AMY1* gene duplications within the context of agriculture.

### Structural haplotypes at the human amylase locus

The amylase locus in the human genome is a ∼212.5-kbp region on chromosome 1 (GRCh38; chr1:103,554,220–103,766,732) which contains *AMY2B*, *AMY2A*, *AMY1A*, *AMY1B*, and *AMY1C* genes (Fig. 1A). This locus is largely composed of segmental duplications with > 99% sequence similarity, which complicates its accurate assembly using short-read sequencing (fig. S1). Using the sequence similarity of segmental duplications and the labeling patterns from the optical genome mapping data from GRCh38 *in silico* map, we defined six distinct amylase segments overlapping the amylase genes, depicted by colored arrows (Fig. 1A and table S1). Using optical genome mapping, which has been previously demonstrated to resolve similar complex regions (*13–15*), we constructed haplotype-resolved diploid assemblies for 98 individuals (n = 196 sampled alleles (i.e., haploid sample size); table S2) and characterized this locus using the copy number and orientation of the amylase segments (Fig. 1A). This approach allowed us to identify 51 distinct amylase haplotypes (fig. S2, A and B, and table S3), of which 7 were previously reported (*7*) (fig. S3). These haplotypes were then classified based on the number of amylase gene copies, adhering to the established nomenclature; HXAYBZ: HX: represents *AMY1*, AY: represents *AMY2A*, and BZ: represents *AMY2B* copy numbers, with superscript “a” indicating ancestral and superscript “r” indicating reference haplotypes (fig. S4) (*7*, *11*). We subsequently defined 30 high-confidence haplotypes (from 117 observed alleles in 81 individuals) that were orthogonally supported by *de novo* assemblies based on long-read sequencing (Fig. 1B and Data Files-S1). This represents the first nucleotide-level reconstruction of the amylase locus at a population scale.

**Fig. 1.**
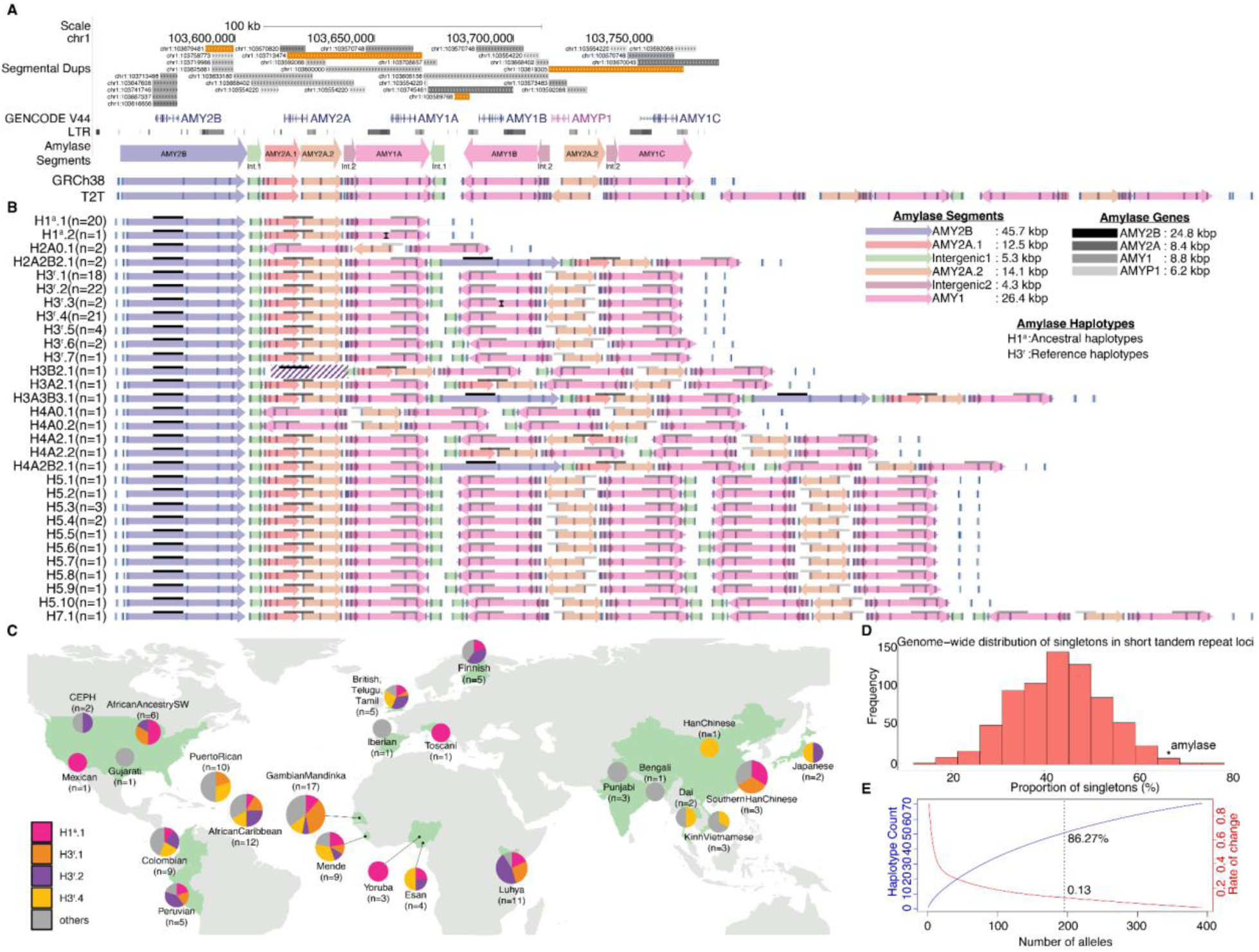
Amylase structural haplotypes identified from present-day humans in this study. (**A**) Segmental duplications (light to dark gray: 90-98% similarity, light to dark orange: > 99% similarity), GENCODE V44 gene annotations, and long terminal repeats (LTRs) are represented as tracks. The lower panel shows amylase segments (colored arrows), and haplotype structures of GRCh38 and T2T-chm13 reference assemblies, represented as *in silico* maps with white backgrounds and vertical blue lines displaying optical mapping labels. The AMY2B segment overlaps the *AMY2B* gene, AMY2A.1 and AMY2A.2 segments overlap the *AMY2A* gene, and the AMY1 segment overlaps the *AMY1* gene. (**B**) The high-confidence amylase structural haplotypes resolved in our dataset (n = 30). The vertical black line in the second AMY1 segment of H3^r^.3 represents the polymorphic label present in three alleles. The diagonal stripes in the second AMY2B segment of H3B2.1 indicate that it is a partial copy of the first AMY2B segment. Haplotype IDs describe: HX: X denotes the number of *AMY1* copies; AX: X denotes the number of *AMY2A* copies; BX: X denotes the number of *AMY2B* copies. The superscript “a’’ denotes the ancestral amylase haplotype structure, and the superscript “r” denotes the reference amylase haplotype structure. The number in parentheses indicates the number of alleles. (**C**) The distribution of common amylase haplotypes across 26 population samples. (**D**) The proportion of singletons for tandem repeat loci (EnsembleTR) across the genome. For adequate comparison, we used the same individuals (n = 33) for whom we were able to reconstruct amylase haplotypes in our dataset. Additionally, we filtered the tandem repeat loci (719 loci, unit length 1-6 bp) that we analyzed to match the number of distinct alleles (n = 21) observed in the amylase locus. The asterisk (*) represents the proportion of singletons among all distinct haplotypes (∼67%, 14 out of 21) detected at the amylase locus. (**E**) Rarefaction and extrapolation sampling curve based on 51 amylase haplotypes, displaying how the number of distinct haplotypes (blue line) is projected to saturate with the increase in the number of alleles. The rate of change (red line, 0.13) indicates the number of novel haplotypes discovered per unit increase in the number of analyzed alleles. The dashed line shows the proportion of estimated number of samples (86.72%) captured in our study.

The length of amylase haplotypes ranged from 111-kbp (H1^a^.1 and H1^a^.2) to 402-kbp (H7.1) (Fig. 1B), capturing those that are structurally identical to the GRCh38 (H3^r^.1) and the T2T-chm13 (H7.2) (*16*) assemblies (Fig. 1A). Four haplotypes, H1^a^.1 (n = 20/117), H3^r^.1 (n = 18/117), H3^r^.2 (n = 22/117), and H3^r^.4 (n = 21/117), were categorized as common, each with an allele frequency exceeding 5% across all populations. These four common haplotypes collectively constitute approximately 70% of all amylase haplotypes (n = 81/117 identified in this study). We also detected singletons, each detected only once (n = 19/117). Despite our limited sample size, we found that all common haplotypes exist in all continental regions (Fig. 1C and fig. S5, A and B). In addition, *AMY1* copy number variations do not exhibit a discernible geographic specificity (p-value = 0.406, Kruskal-Wallis rank sum test).

Out of 30 distinct amylase haplotypes, we identified 19 (63%) as singletons. To infer the relative mutation rate of the amylase locus, we compared this number to that of tandem repeats. To avoid potential biases, we used the same 33 individuals that are present in both the tandem repeat database (EnsembleTR) and our dataset. Only 21 of the 30 distinct amylase haplotypes were present in these 33 individuals. To make sure the analysis was consistent across different genomic regions, we only considered tandem repeat loci that had 21 detected alleles in the human population. This yielded 719 tandem repeat loci (fig. S6). We found that the proportion of singleton haplotypes was significantly higher for the amylase locus than the genome-wide average for the 719 tandem repeat loci (the observed empirical percentile = 0.017, Fig. 1D). This observation is informative for understanding the mutation rate at this locus because the allele frequency spectrum and proportion of singletons are determined by mutation rate and genetic drift (*17*). By using the same individuals and matching the number of distinct haplotypes in our comparison, we control for demographic biases and provide a relative estimation of the mutation rate for the amylase locus. Considering that short tandem repeats have mutation rates as low as 10^−8^ (similar to single nucleotide variant mutation rate) and, in some cases, can be as high as 10^−2^ mutations per locus per generation (*18*), our analysis encompasses the entire range of mutation rates found in the genome. We acknowledge that an ideal future comparison would involve other amylase-like loci, which exhibit similar mutational mechanisms and levels of structural variation, and that are resolved using comparable approaches once such databases become available. In the meantime, when we repeated this analysis for the complex 3q29 locus, known for its segmental duplication-rich nature and high levels of structural variation and resolved with similar approaches (*15*, *19*), we found 11 singletons (fig. S7). Thus, the amylase locus is mutating faster than a typical structural variation hotspot and 98.3% of analyzed tandem repeats.

To understand how much of all amylase haplotypes that exist in the human population has been captured by our study, we conducted rarefaction analysis on the 98 samples and found that we have identified all common haplotypes with a frequency of >= 5% (Fig. 1E and fig. S8, A and B) and 86% of all haplotypes overall (fig. S9).

### Strong negative selection limits functional variation among amylase gene copies

To systematically assess the selection pressure on the amylase coding sequences, we examined the degree of protein-coding sequence variation associated with our high-confidence amylase haplotypes (30 haplotypes from 117 alleles). Gene annotation predicted 582 distinct intact protein-coding amylase gene copies in 117 alleles and we experimentally validated these copy number predictions using digital droplet PCR on 18 randomly selected individuals (R^2^ = 0.94; fig. S10A and table S4, S5). The reconstruction of the coding sequence phylogeny revealed that all human amylase gene copies can be robustly clustered into three distinct types: *AMY2B*, *AMY2A*, and *AMY1* (bootstrap value = 96%; Fig. 2A and fig. S11). We found that the *AMY2B*, *AMY2A*, and *AMY1* genes had 23, 23, and 36 fixed coding sequence variations unique to each type, resulting in 6, 11, and 19 gene-specific amino acid differences, respectively (table S6). Based on the coding sequence alignment (Data Files-S2), we estimated the synonymous and nonsynonymous substitution (dN/dS) ratios using codeml (*20*), where we found no evidence for lineage-specific selection pressure acting on any of the amylase gene types (FDR adjusted p-value from χ^2^ test > 0.05; table S7). In contrast, all amylase gene types show significant signatures of negative selection (FDR adjusted p-value from χ^2^ test < 0.05; Fig. 2A and table S7). These observations suggest that negative selection (dN/dS ratio < 1) has acted to retain the amino acid sequence of amylase gene copies, both within and between the three amylase gene types. It is of note that we identified two amino acid variants at positions 211 and 366 (Accession: P0DUB6) that could contribute to functional differences and may have biomedical significance. (Fig. 2B, fig. S12, and table S8).

**Fig. 2.**
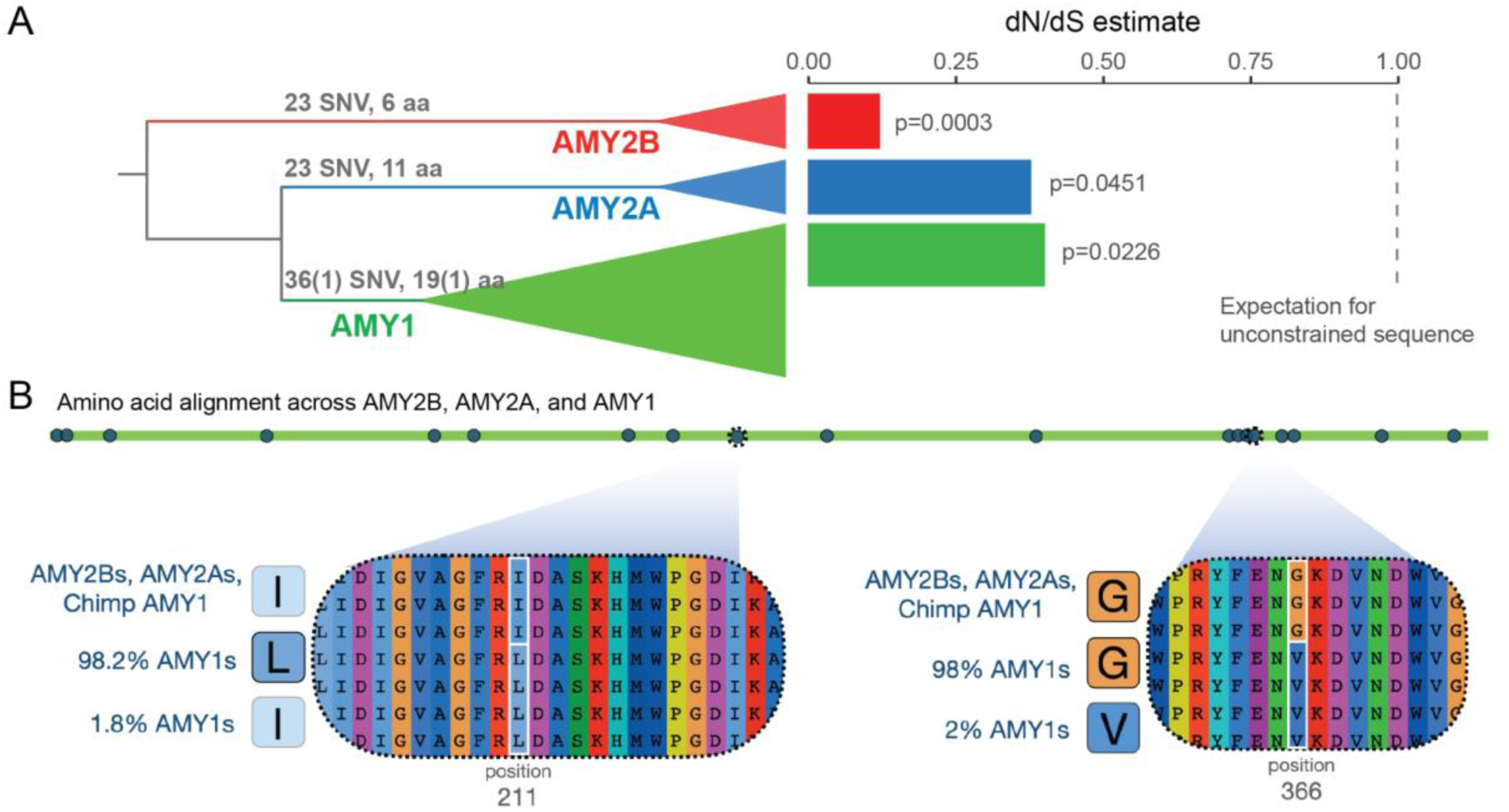
The variants in amylase coding sequences and negative selection on three amylase gene types. (**A**) The maximum likelihood phylogenetic tree of amylase coding sequences (left) and dN/dS estimate for each amylase gene type (right). The phylogenetic tree is rooted with a coding sequence from the sheep genome (Oar_rambouillet_v1.0). The number of nucleotide and resultant amino acid changes that are paralog-specific are indicated. The numbers in parenthesis indicate nucleotide and amino acid changes that are variable within the AMY1 branch. The numbers to the right of each bar represent the FDR-adjusted p-value for the likelihood ratio between H0_2_ (dN/dS ratio is fixed to one on the foreground branches) and H1 (two dN/dS ratios are allowed on the foreground and background branches, respectively) for each gene type. (**B**) The positions of amino acid variants within and between *AMY2B*, *AMY2A* and *AMY1* protein sequences. The 211 and 366 positions are highlighted because they overlap with a conserved section of the amylase protein sequence and have a predicted functional impact (AlphaMissense (*21*). The coordinates are based on the residues of the amylase enzyme from UniProtKB (Accession: P0DUB6).

Our coding sequence data from gene annotations covering 582 intact amylase gene copies confirmed the explicit difference between *AMY1, AMY2A,* and *AMY2B* genes, which is crucial to distinguish the expression of these genes across tissues. Indeed, according to existing data portals, Genotype-Tissue Expression and The Human Protein Atlas, *AMY2A* and *AMY2B* are expressed in the pancreas and, to some level, in adipose and brain tissues, while the *AMY1* gene is expressed primarily in the parotid salivary gland (fig. S13). These observations are consistent with the idea that *AMY1* first emerged in the ancestor of great apes, resulting in a gain of expression in the parotid gland tissues (*5*). Furthermore, subsequent *AMY1* gene duplications in the human lineage appear to affect the dosage of amylase in the parotid salivary glands (*1*). Among the haplotypes we identified in this study, we found 110 amylase pseudogenes and showed they share a single phylogenetic origin from an ancestral incomplete gene duplication of the *AMY2A* gene (*22*) (fig. S14). Thus, the pseudogenization of *AMY2A* is more likely due to a single mutational event rather than a loss of constraint and the repeated occurrence of novel loss-of-function variants.

### Evolution of the copy number of AMY1 gene

We showed above that all human amylase gene copies can be robustly clustered into three distinct types: *AMY2B*, *AMY2A*, and *AMY1.* However, to specifically study the evolution of the copy number of the salivary *AMY1* gene, we needed to identify the sequences within the amylase locus that were the most phylogenetically informative. To achieve this, we systematically evaluated the haplotype variation in 117 alleles by aligning sequences and identified a consensus sequence for each amylase segment (fig. S15, A and B). We identified an interval (from 22,850 to 26,730 bp; Data Files-S2) within the AMY1 segment (fig. S15C), where all observed AMY1 segments (n = 337) can be phylogenetically structured into three distinct clusters; AMY1A (n = 124), AMY1B (n = 99), and AMY1C (n = 114) (fig. S15D). These clusters correspond to the segments represented in the GRCh38 assembly. Further, the chimpanzee (panTro6) and gorilla (gorGor6) reference genomes each contain only one AMY1C-like segment diverged from human H1^a^.1 haplotypes (figs. S16 and S17). These findings suggest that the common ancestor of humans and chimpanzees possessed a single AMY1C-like segment, and the AMY1A and AMY1B segments have evolved only in the human lineage (fig. S17). Note that the bonobo reference genome (panPan3) contains two AMY1 segments: one ancestral and one resulting from an independent, bonobo-specific duplication, as determined through synteny and phylogenetic analysis (figs. S16 and S17). One of these duplications is likely non-functional due to a previously reported a disrupted coding sequence (*1*), a finding corroborated by the most recent annotation (RS_2024_02/NHGRI_mPanPan1-v2.0_pri).

To further understand the evolution of the *AMY1* gene copy number, we aligned all AMY1 segments from the most common haplotypes, H3^r^.1 and H3^r^.2, that harbor all three AMY1 segment types. Two independent Bayesian phylogenies based on these alignments indicate that the AMY1B segment arose from the AMY1C approximately 140 to 270 kilo (thousand) years ago (KYA), followed by the duplication of AMY1A from the AMY1B segment approximately 120 to 240 KYA (fig. S18 and table S9). Gene conversion between the GC-rich segmental duplications complicates time estimation based on a molecular clock and is a known phenomenon at the amylase locus (*6*, *23*). Considering gene conversion between AMY1 segments, the actual duplication dates are expected to be older than the estimates above. Some studies have used single nucleotide variants in the flanking regions to infer the phylogenetic history of this locus, positing coalescence dates for human amylase locus at ∼279 and ∼450 KYA (*11*, *12*). However, as described previously (*7*), we found that the linkage disequilibrium between the flanking single nucleotide variants and structural haplotypes is low (e.g., H1^a^.1 average: R^2^ = ∼0.26 and median R^2^ = ∼0.03) (fig. S19), complicating the time estimation of *AMY1* duplications using flanking regions. Therefore, our estimates avoid these complications and support the conclusion that the initial *AMY1* gene duplications substantially predated out-of-Africa migrations, by at least 30 KYA (table S9).

### AMY1 copy number variation in archaic hominin genomes

A complementary approach for estimating the relative timing of gene duplications involves analyzing the read-depth of unique k-mers in ancient human and archaic hominin genomes. We first tested a k-mer approach using the GeneToCN algorithm on short-read sequencing data (*26*) to estimate the copy numbers of the *AMY1*, *AMY2A*, and *AMY2B* genes in 116 present-day human genomes (table S5). Notably, the k-mer approach achieved an R² > 0.99 correlation for 32 individuals, each with both haplotypes of the amylase locus reconstructed (fig. S10B). We tested our k-mer approach in a separate cohort of 101 samples with digital PCR validation data (R² = 0.95; fig. S10A). These results suggest that the GeneToCN algorithm may be more accurate than digital PCR estimations and a viable option for estimating *AMY1* gene copy numbers in short-read whole genome datasets.

We then aimed to estimate *AMY1* gene copy number in eight archaic hominin genomes using two approaches: (1) the validated k-mer method described above, as well as (2) an independent read-depth analysis (table S10). Given the varying genome-wide coverage in most of these genomes, we needed to conduct a downsampling analysis to empirically determine that 1X and 5X genome-wide coverage provides >85% and >95% accuracy in estimating *AMY1* copy number, respectively (fig. S20 and table S11). Taking into account GC bias and coverage within the amylase locus for each sample (see Methods), we were able to reliably estimate the *AMY1* copy number for eight archaic hominin genomes (table S10). We found increased *AMY1* copy number in two Eastern and one Western Neanderthals, as well as one Denisovan genome (Fig. 3A, fig. S21 and table S10). These include the Altai Neanderthal (2.6 copies), Denisova 2 (8 copies), GoyetQ56-1 (5.0 copies), and Mezmaiskaya 2 (4.7 copies). Previous read-depth analysis of the Altai Neanderthal genome found no evidence for an increase of *AMY1* gene copy number (*10*, *27*). By incorporating eight archaic hominin genomes, we have now detected signatures of *AMY1* gene duplication in some of the Neanderthals and Denisovan genomes, indicating that *AMY1* duplication was actually polymorphic among archaic hominins.

**Fig. 3.**
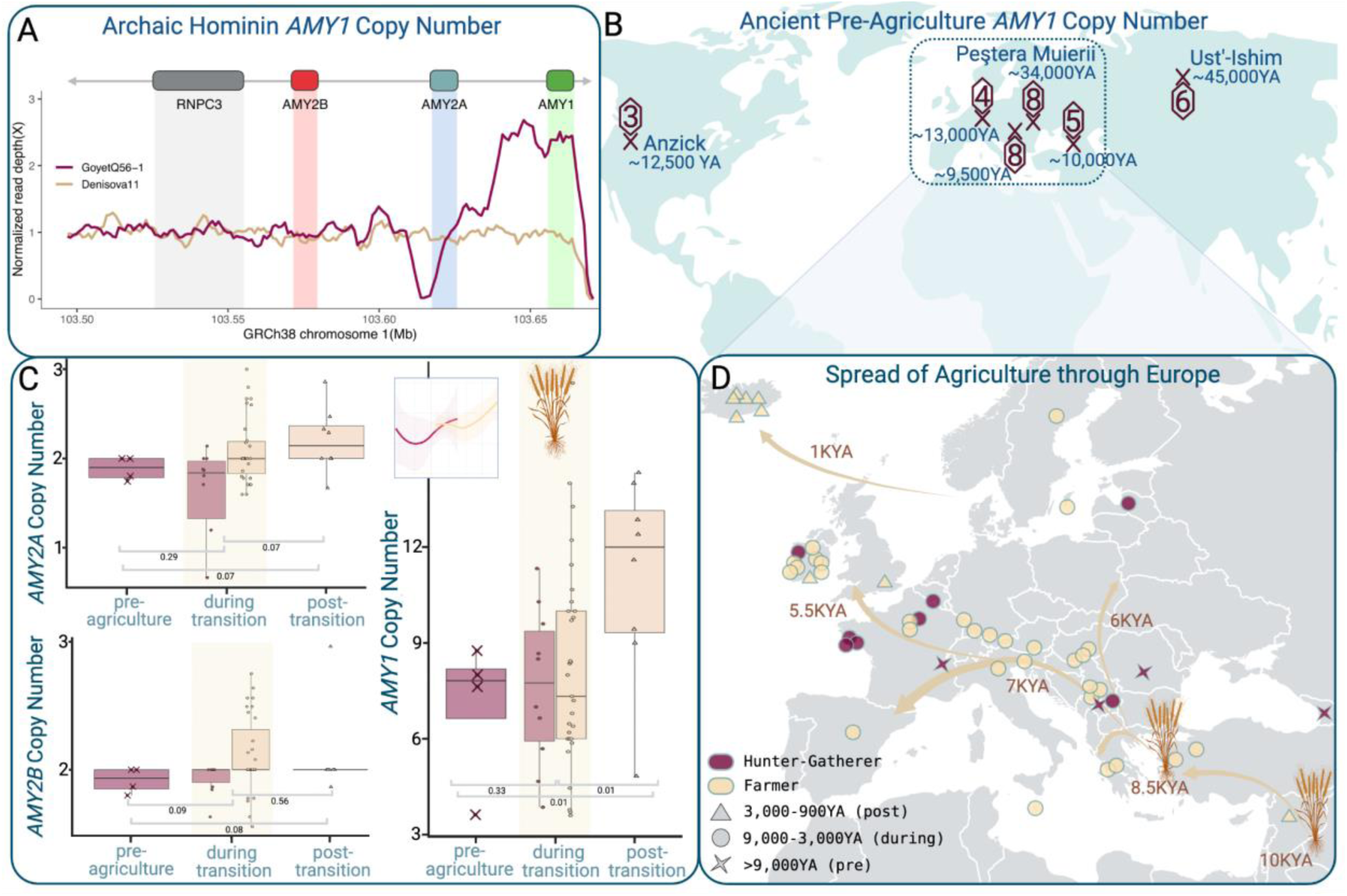
Amylase gene duplication and the history of agriculture. (**A**) The read-depth of amylase locus spanning *RNPC3*, *AMY2B*, *AMY2A*, and *AMY1* genes (chr1:103494306-103668306) for GoyetQ56-1 (maroon line), a Neanderthal excavated in present-day Belgium, showing signatures of *AMY1* duplication, and for Denisova 11 (beige line), a hybrid hominin (Neanderthal and Denisovan) excavated from present-day Russia, showing signatures consistent with an ancestral single-copy AMY1 haplotype. The maroon and beige lines indicate average read-depths with 5-kbp window and 1-kbp step for each sample. The average read-depth of each 5-kbp window was normalized by the average read-depth of the *RNPC3* gene. Only uniquely mapped reads were used for this visualization. (**B**) A world map displaying the locations of ancient human samples. Sample locations are indicated with an ‘X’, with corresponding hexagons showing the estimated *AMY1* copy number. Carbon dating (Kilo Years Ago; KYA) estimated for each sample is indicated in blue. (**C**) Amylase copy number estimations from Europeans who were farmers (beige) and hunter-gatherers (maroon). Samples are binned on the x-axis according to three time periods; pre-agriculture (before the transition to agriculture, >9 KYA), during the transition (∼ 9 - 3 KYA), and post-transition (complete transition to agriculture, ∼ 3 KYA - 900 years ago). The shape of data points corresponds to sample dating estimates (Xs for more than 9 KYA, circles for 9 - 3 KYA, and triangles for 3 KYA - 900 years ago). The inset of the right panel shows nonparametric regression lines for *AMY1* copy number across time, for hunter-gatherers (maroon) and farmers (beige), with confidence intervals in lighter corresponding color, respectively. (**D**) Zoomed-in map showing the spread of agriculture into Europe from Asia. Major agricultural footholds are indicated by wheat pictograms. Tan arrows show general trends of human agricultural migration throughout Europe, with predicted time periods (*24*, *25*). Ancient human samples that were analyzed for amylase copy number are annotated with shapes and colors for time period and lifestyle, respectively.

The *AMY1* duplication in these archaic hominins could be postulated by four scenarios. First, it is possible that because of the complexity of the locus and the complications inherent in archaic hominin genomic sequencing, there may be a technical bias in our detection. However, the observation of duplications using two different approaches and in multiple genomes provides confidence in our results. Second, it is plausible that introgression into archaic hominins from humans may explain the presence of duplications in the former. While still a possibility to be tested when more archaic hominin genomes are available, we observed the duplication in western Neanderthals (e.g., Goyet), which shows no signature for introgression from humans (*28*, *29*). Third, it is plausible that the duplication evolved independently in the archaic hominin lineage. However, we argue that two independent duplications (one in humans and another in archaic hominins) in less than a million years is unlikely, given that initial duplications from single *AMY1* copy haplotypes are rare in great apes (*4*) (fig. S16). The fourth and, in our opinion, the most plausible scenario is that the *AMY1* gene might already have been copy number variable before the human-Neanderthal/Denisovan divergence (∼800 KYA (*30*)), albeit to a limited extent as compared to what is observed in present-day humans. Overall, our results suggest a complex history of amylase duplications, which will be further scrutinized as more high-coverage archaic hominin genomes become available.

### The frequency of higher copy number of AMY1 has increased in the last 10,000 years, coinciding with the agricultural transition

To explore the changes in allele frequency of *AMY1* copy number since the migration out of Africa 60 - 90 KYA, we analyzed the genomes of 73 ancient human genomes (table S12). The oldest genome analyzed was the Ust’Ishim sample from Siberia (∼45 KYA), which has six *AMY1* gene copies per diploid cell. The Peştera Muierii sample from Romania (∼34 KYA) has eight *AMY1* gene copies per diploid cell. This indicates that high *AMY1* gene copies (*defined here as* >= 6 copies per diploid cell) had already spread across Eurasia as far back as ∼45 KYA (*31*)) (Fig. 3B).

We next analyzed the ancient human genomes in the context of agricultural development and found a general trend where the *AMY1* gene copy number is significantly higher among samples excavated from archaeological agricultural contexts compared to those from hunter-gatherer contexts (p-value = 0.037; fig. S22). To further investigate the intricacies of this trend, we examined ancient human genomes from Europe, where we have a clear timeline of the Neolithic transition. Specifically, based on archaeological evidence, we categorized genomes based on three distinct European time periods: the pre-agricultural period (> 9 KYA), where all our samples are from hunter-gatherers; the agricultural transition period (9 to 4 KYA), representing the long period of transition to agriculture in Europe; and the agricultural period, during which no hunter-gatherers are recorded in our sample datasets (< 3 KYA). These period choices were made with the assumption that the putative adaptive effects of agricultural diets would take some time to become visible at the population level. We found that pre-agricultural genomes already harbored four to eight *AMY1* copies per diploid cell (table S12). We also observed a consistent and significant increase in *AMY1* copy numbers across these periods (*e.g.*, during vs. post p-value < 0.01; Fig. 3, C and D), and found similar trends for the *AMY2A* genes (Fig. 3C) and non-European samples (figs. S23 and S24). These findings support the notion that haplotypes with higher numbers of amylase gene copies have increased in frequency over the last 10 KYA. We found no significant differences in *AMY1* copy number between samples excavated from agricultural versus hunter-gatherer archaeological contexts during the “agricultural transition” period when farmers and hunter-gatherers shared the same habitat.

Taken together, these findings are consistent with the idea that either neutral evolution or weak-adaptive forces, perhaps due to preagricultural experimentation with food processing techniques (*32*), such as flour production from wild cereals (*33*), have retained the wide range of standing *AMY1* copy number variation in preagricultural Europe. The gradual increase in starch availability as Europe transitioned to an agricultural lifestyle (*34*) likely underlies the observable adaptive force acting on high copy number haplotypes, explaining the increase of *AMY1* copy number in late post-agriculture European populations.

### Multiple mutational origins that resulted in a burst of copy number variation since the initial AMY1 duplication

We next investigated the likely mutational origins of present-day amylase haplotypes. To do this, we first examined the relationship among the four most common haplotypes, H3^r^.1, H3^r^.2, H3^r^.4, and H1^a^.1. Building upon the previously reported link between one-copy and three-copy haplotypes (*35*) and the fact that we observe only three types of AMY1 segments, we propose an evolutionary model linking the ancestral chimpanzee-like haplotype (H1^a^-like) to the common three-copy haplotypes (H3^r^.1 and H3^r^.2) (fig. S25). According to this model, the initial duplications of *AMY1* starting from H1^a^-like ancestral haplotype led to the emergence of H3^r^ haplotypes. Given that multiple mutational steps are required to move from one-copy common haplotypes to other common haplotypes and that we do not observe any intermediate haplotypes in the present-day human genomes, the duplication from one-copy to three-copy haplotypes is likely to have occurred only once in the human lineage. This is also supported by the absence of duplications in the amylase segments within the H1^a^.1 haplotype, which would impede non-allelic homologous recombination (NAHR) events. In contrast, H3^r^.1 and H3^r^.4 haplotypes harbor copies of identical and unidirectional sequences (AMY2A.2 segments), providing an ideal template for recurrent NAHR events leading to the diverse haplotypes we see today.

To further investigate the mutational relationships between the H3^r^ haplotypes and other extant haplotypes, we utilized dotplots and sequence alignments to delineate the breakpoints of structural differences (Supplementary Results). In parallel, we conducted a scan for PRDM9-binding motifs within the amylase locus to pinpoint possible recombination sites (Fig. 4A, fig. S18 and table S13). By integrating all these observations, we were able to construct a putative evolutionary path with the fewest plausible mutational steps that can explain the origin of present-day amylase haplotypes starting from NAHR-prone H3^r^ haplotypes (fig. S26, S27; Supplementary Results). Our proposed evolutionary model of mutational events is consistent with our hypothesis regarding the central role of the H3^r^ haplotypes in seeding extant haplotypic variation and offers three major insights into the evolution of the amylase locus in humans.

**Fig. 4.**
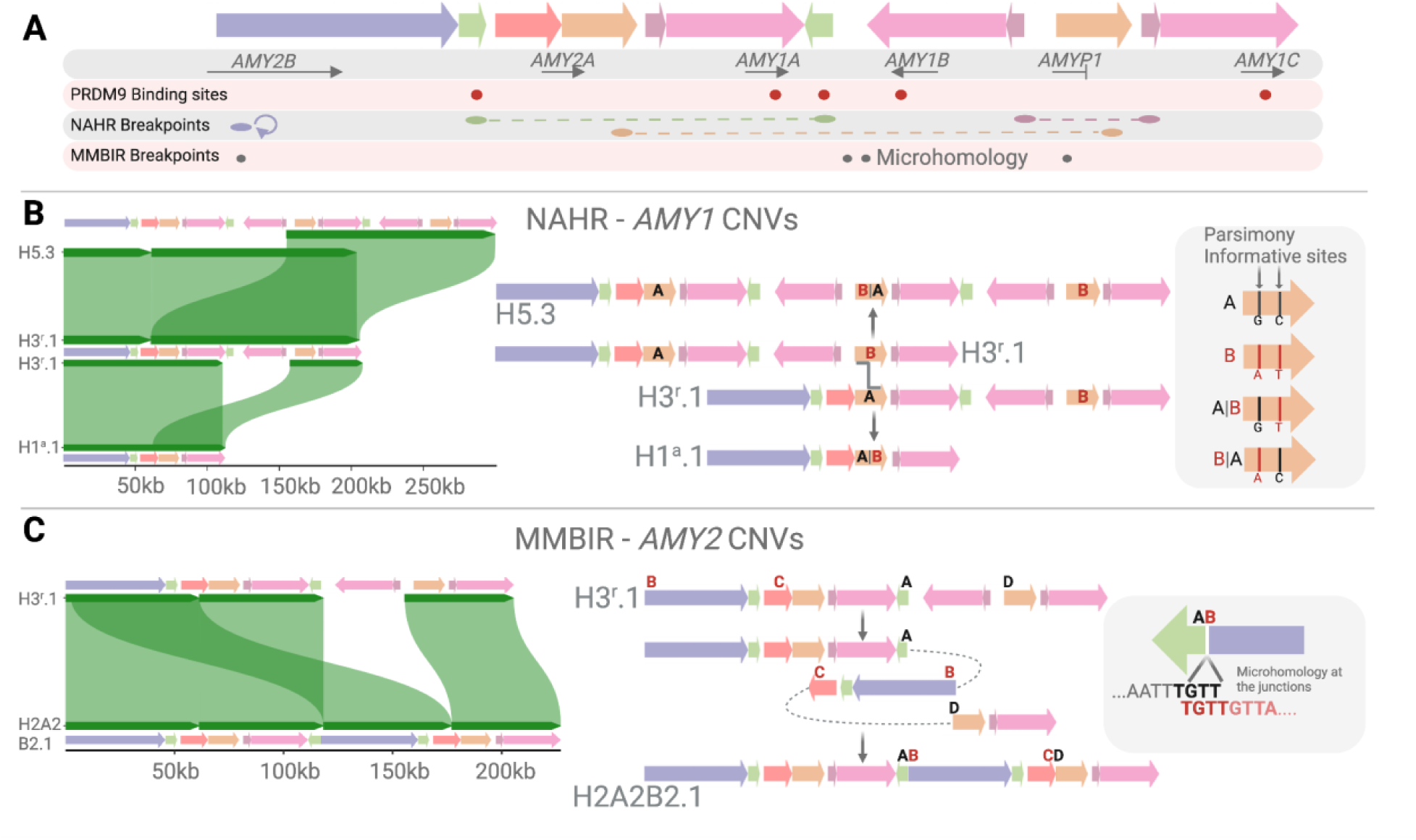
The evolutionary and mutational connections among common haplotypes. (**A**) The structural variation breakpoints and recombination hotspots in the amylase locus. The colored arrows represent amylase segments. The PRDM9 binding sites are represented with red dots. The non-allelic homologous recombination (NAHR) breakpoints are represented with purple, green and orange dots, and dashed lines. The microhomology-mediated break-induced replication breakpoints are represented with gray dots. (**B**) NAHR-mediated duplication and deletion of the *AMY1A*-*AMY1B* cluster. The AMY2.2 (orange) segment serves as the recombination substrate for the crossover, resulting in the duplication or deletion of the *AMY1A*-*AMY1B* cluster as illustrated in the middle panel. Chimeric AMY2.2 segments have been identified, utilizing parsimony informative sites within the AMY2.2 segment (right panel). (**C**) Microhomology-mediated break-induced replication based copy-number gain resulting in the formation of H2A2B2.1. The middle panel shows the mutational mechanism. Four nucleotides of microhomology internal to the breakends were identified at the breakpoint junction (right panel).

First, we found evidence for recurrent NAHR events among common haplotypes (H3^r^.1, H3^r^.2, and H3^r^.4) harboring the AMY1A and AMY1B segments with breakpoints in the AMY2A.2 (orange) segment. These NAHR events, which may occur among different haplotype combinations, could concurrently result in the duplication and deletion of two *AMY1* gene copies (*e.g.*, Fig. 4B and fig. S28; Supplementary Results). Therefore, while other less likely scenarios are possible, NAHR-based deletion and duplications underlie copy number variation of the AMY1 segments and thus *AMY1* genes. Specifically, as this proposed mechanism always adds or deletes two copies of the *AMY1* genes, our finding explains how most human haploid genomes harbor odd-numbered *AMY1* gene copies (*7*) (fig. S29). It would seem that the majority of H1^a^.1 haplotypes among present-day humans have predominantly originated from H3^r^ haplotypes. If true, this hypothesis explains the homoplastic occurrence of H1^a^ haplotypes across the amylase phylogenetic tree (fig. S19C) and the lack of an out-of-Africa signal in H1^a^ nucleotide diversity, which would be expected if the H1^a^ haplotypes arose before out-of-Africa migrations (table S14). Therefore, we argue that although the H1^a^.1 haplotype is structurally identical to the ancestral human amylase haplotype, the current versions in our datasets have arisen recurrently from H3^r^ haplotypes.

Second, we characterized three microhomology-mediated break-induced replication events and identified the accompanying microhomologies at the breakpoint junctions (Fig. 4C; Supplementary Results). Even though these three rearrangements constitute only five alleles (H2A2B2.1, H3^r^.6, and H3B2.1) (∼ 4%), they hold substantial biological relevance since H2A2B2.1 and H3B2.1 harbor duplications of the *AMY2* genes. Thus, different mechanisms drive the copy number variation of the salivary *AMY1* genes and pancreatic *AMY2* genes, with the lower copy number variation of *AMY2* genes explained by the slower rate of non-recurrent microhomology-mediated break-induced replication events (*36*).

Third, recurrent NAHR-mediated inversion events at the amylase locus, similar to those described previously (*37*), underpin the mutational connections between the common H3^r^.1, H3^r^.2, and H3^r^.4 haplotypes (fig. S30), as well as several other inversions among extant haplotypes (Supplementary Results). Given that inversions underlie the structural differences between the majority of the distinct haplotypes, their functional and adaptive relevance presents an intriguing avenue for future research.

## Discussion

In this study, we have dissected the evolution of the amylase locus. First, we hypothesize that the initial duplications of the *AMY1* genes occurred once through multiple duplications, evolving from a one-copy ancestral haplotype to the three-copy present-day haplotypes frequently observed in our dataset. Analysis of archaic hominin genomes suggests that these initial duplications occurred well before the split of the human-Neanderthal/Denisovan. This observation is concordant with the recent evidence of Neanderthal starch consumption (*38*), and perhaps the availability of cooked starch in archaic hominins made possible through the domestication of fire (*39*) (Fig. 5).

**Fig. 5.**
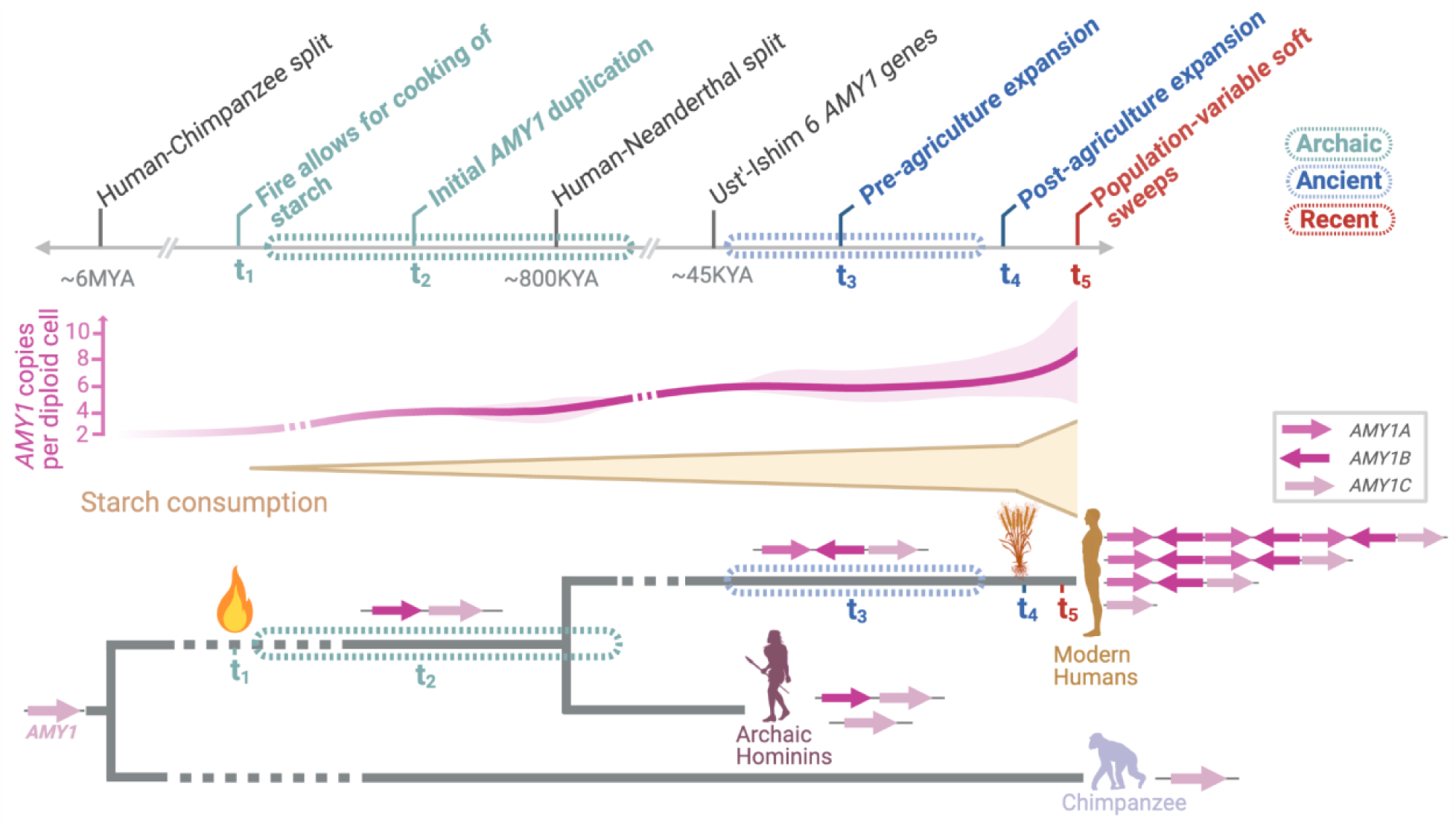
An evolutionary model of the human amylase genes and resulting hypotheses. Top: A timeline of human amylase locus evolution based on the results of this study, with relevant events indicated on top. Middle: A schematic view showing the increase in *AMY1* copy number variation and the mean number of *AMY1* copies in present-day human populations throughout history as starch consumption increases. Bottom: A phylogenetic representation of the hypothesized amylase duplication timeline.

Second, we hypothesize that selection acted on abundant standing *AMY1* copy number variation at this locus rather than on *de novo* variants. We observed a wide range of *AMY1* gene copy number variation (3 - 9 copies) in samples that predate agriculture. We further found that late agricultural populations consistently harbored amylase haplotypes with higher *AMY1* copy numbers (Fig. 5). Yet, our data do not indicate strong, pervasive selection. Instead, we propose that partial soft sweeps involving pre-existing amylase structural haplotypes may have influenced *AMY1* copy number variation in relation to historical starch consumption patterns in different populations. The effects of amylase gene duplications on taste preferences and starch metabolism may have also predisposed humans to prefer and tolerate consuming wild grains, as reported for the Mesolithic groups in the Balkans (*40*), facilitating the adoption of starch-rich diets and eventual domestication of these plants. Overall, our results support the complex narrative of the transition to agriculture, which includes the replacement of western hunter-gatherers by Anatolian farmers in Europe (*41*), potentially bringing with them amylase haplotypes that harbor higher *AMY1* gene copies. Similarly, transient interactions between hunter-gatherer and agricultural groups (*42*) could explain the similar copy numbers observed between these groups during the transition period. Future comparative analyses focusing on partial soft sweeps in indigenous populations with recently diverged dietary habits involving heavy starch consumption—such as the Andeans (*43*, *44*) could provide further insights into these signatures of selection at the human amylase locus.

Third, we found that NAHR and microhomology-mediated break-induced replication underlie the copy number variation of *AMY1* and *AMY2* genes, respectively, explaining the different rates of their structural evolution. The extremely high rate of structural variation due to NAHR led to remarkable *AMY1* copy number variation with distinct mutational propensities. For example, our evolutionary model involving common H3^r^ haplotypes suggests recurrent NAHR events mediated by highly similar sequences (> 99%), resulting in duplications or deletions of both *AMY1A* and *AMY1B* genes. In contrast, the H1^a^.1 haplotype harboring a single *AMY1* copy and divergent *AMY2* genes rendering it less susceptible to NAHR events. Therefore, the mutation types and rates of the amylase locus may differ depending on extant haplotype variation in a population, especially in bottlenecked populations such as indigenous Americans (*45*). It is a distinct possibility that a bottlenecked population ends up with a very high frequency of H1^a^.1 due to drift. In this case, the absence of segments with highly similar sequences within H1^a^.1 haplotype would mitigate recurrent NAHR events, resulting in a slower accumulation of variation in this population. In contrast, if one of the larger amylase haplotypes were to become prevalent due to drift, the rate of variation would increase exponentially. Within this general context, one interesting question for future work is whether larger amylase haplotypes experience negative selection due to increased genomic instability. Taken together, our study underscores how gene duplications in early human history provided the genetic foundation for dietary flexibility during agricultural innovations, contributing to modern human evolution.

## Materials and Methods

### Sample Collection and Datasets

Samples (n = 98) from diverse populations studied by the Human Genome Structural Variation Consortium (HGSVC) (*19*), Human Pangenome Reference Consortium (HPRC) (*46*) and Genome In a Bottle (GIAB) (HG002) datasets were included in this study. These datasets consist of African (n = 43), American (n = 24), East Asian (n = 13), European (n = 9), and South Asian (n = 9) (table S2) individuals. Bionano Genomics optical genome mapping and PacBio HiFi datasets of HGSVC samples were downloaded from http://ftp.1000genomes.ebi.ac.uk/vol1/ftp/data_collections/HGSVC2/working/ and http://ftp.1000genomes.ebi.ac.uk/vol1/ftp/data_collections/HGSVC3/working/, and HPRC and GIAB samples containing phased assemblies and optical genome mapping datasets were downloaded from https://s3-us-west-2.amazonaws.com/human-pangenomics/index.html?prefix=working/).

### Amylase Segments

We characterized a ∼212.5-kbp region on chromosome 1 (GRCh38; chr1:103,554,220– 103,766,732) overlapping paralogous copies of segmental duplications that have > 99% sequence similarity. The amylase segments, depicted by colored arrows, were modified from a previous study by Usher and colleagues (*7*), and determined based on the labeling pattern from the Bionano Genomics optical genome mapping data and the sequence similarity between segments using blastn (BLASTN 2.9.0+) (*47*). First, using optical genome mapping *in silico* map labeling patterns of GRCh38 reference assembly, amylase segments were identified using pairwise alignments between smaller duplicons within segmental duplications. Then, to obtain accurate coordinates of segments, the GRCh38 fasta sequence of the locus was aligned against itself using blastn (BLASTN 2.9.0+). The alignments with less than 99% sequence similarity and 60% coverage were filtered out and coordinates of all reference segments were finalized. We identified one copy of AMY2B (purple; 45.7-kbp), two copies of Intergenic1 (green; 5.3-kbp), one copy of AMY2A.1 (red; 12.4-kbp), two copies of AMY2A.2 (orange; 14.1-kbp), three copies of Intergenic2 (maroon; 4.3-kbp), and three copies of AMY1 (pink; 26.3-kbp) segments within the GRCh38 amylase locus (table S1). Next, the GRCh38 segment sequences were used as a guide to detect amylase segments from each sample included in this study using blastn. Same sequence similarity (> 99%) and coverage thresholds (> 60%) were used to obtain the amylase segment coordinates from each sample. Amylase segments in nonhuman primates, chimpanzee (Clint_PTRv2_panTro6), bonobo (Mhudiblu_PPA_v0_panPan3) and gorilla (Kamilah_GGO_v0_gorGor6), were detected by using the same approach described above with sequence similarity threshold of >= 90% using genome reference assembly fasta files that were obtained from University of California Santa Cruz Genome Browser.

### De novo assembly of optical genome map dataset

Optical genome maps of HGSVC and HPRC samples were *de novo* assembled and using the Bionano Solve v3.5 (https://bionanogenomics.com/support/software-downloads/) assembly pipeline, with default settings as described previously (*48*) (fig. S31A).

~~~
python2.7 Solve3.5.1_01142020/Pipeline/1.0/pipelineCL.py -T 64 -U
-j 64 -jp 64 -N 6 -f 0.25 -i 5 -w -c 3 \
-y \
-b ${bionano_bnx} \
-l ${output_dir} \
-t Solve3.5.1_01142020/RefAligner/1.0/ \
-a
Solve3.5.1_01142020/RefAligner/1.0/optArguments_haplotype_DLE1_s
aphyr_human.xml \
-r ${reference_genome}
~~~

A pairwise comparison of DNA molecules (min. 250-kbp) was generated to produce the initial consensus genome maps. During an extension step, molecules were aligned to genome maps, and maps were extended based on the molecules aligning past the map ends. Overlapping genome maps were then merged. Extension and merge steps were repeated five times before a final refinement of the genome maps. Clusters of molecules aligned to genome maps with unaligned ends > 30-kbp in the extension step were re-assembled to identify all alleles. To identify alternate alleles with smaller size differences from the assembled allele, clusters of molecules aligned to genome maps with internal alignment gaps of size < 50-kbp were detected, and the genome maps were converted into two haplotype maps. The final genome maps were aligned to the reference genome, GRCh38.p12.

### Haplotype detection and single molecule support using optical genome maps

To resolve haplotype structures at the amylase locus, *de novo* assemblies of optical genome maps of samples were first aligned to the GRCh38.p12 human genome reference assembly. Next, alignment results were visualized using the Bionano Access™ software to resolve amylase haplotypes. Haplotypes were identified from these visualizations using amylase segments and *in silico* map labeling patterns as a guide. Next, molecule support for each haplotype was evaluated using the following steps (fig. S31B): contigs aligning to the region of interest (ROI) (GRCh38; chr1:103,554,220–103,766,732) were detected. Any samples with greater than or less than two contigs aligning to the ROI were excluded. Next, ROIs in each contig were identified and extended 10-kbp on both proximal and distal regions. The number of molecules aligning to the ROI of the amylase locus was collected, and the average molecule length was calculated for each contig. If the average molecule length was less than the ROI, the number of molecules that were anchored in the proximal and distal unique regions was counted. Next, the length of the ROI was divided by the average molecule length, and new smaller ROIs were identified based on quotient, and the number of molecules was counted for each new ROI. Only haplotypes supported by three or more molecules were considered accurate haplotypes (Data File-S1).

### De novo assembly and orthogonal support

PacBio HiFi long-read sequencing dataset from HGSVC (n = 60) was *de novo* assembled using hifiasm (*49*) with the following command:

~~~
hifiasm -o ${outputdir}${sampleID}.asm -t 32 ${reads}
~~~

Publicly available phased assemblies from the HPRC were analyzed to validate the optical genome mapping based haplotype structures, as described previously (*15*). Validation process for both HPRC and HGSVC assemblies includes the following steps: First, fasta files of HPRC phased assemblies and HGSVC hifiasm assemblies were converted to *in silico* maps using fa2cmap_multi_color.pl (Bionano Solve^TM^ v3.5.1). The resulting *in silico* maps were aligned to the GRCh38p.12 reference assembly *in silico* map using the refAligner tool (Bionano Solve^TM^ v3.5.1).

~~~
Solve3.5.1_01142020/RefAligner/1.0/RefAligner   -ref
${referenceFile} \ -maxthreads ${threads} \
-i ${inputfile} \
-o ${outputdir}EXP_REFINEFINAL1
~~~

Next, a visual pairwise comparison of HPRC optical genome mapping and phased assemblies for each sample was performed using Bionano Access^TM^. Then, amylase segments in each sample with a haplotype consistent with the optical genome mapping haplotype were detected using blastn (BLASTN 2.9.0+), as described in the ‘Amylase Segments’ part. Next, all orthogonally supported haplotype alignments were visualized using pygenomeviz (*50*) to confirm haplotype structures. Finally, haplotypes were clustered using pgr-tk v0.5.1 (*51*) (fig. S4).

### Significance of AMY1 copy number distribution across populations

To evaluate whether the *AMY1* gene copy numbers exhibit population-specific patterns, we performed Kruskal-Wallist rank sum test. This non-parametric test was used to determine if there are statistically significant differences between the medians of three or more independent groups. We chose this test due to its robustness in handling non-normally distributed data. We conducted our analysis on R Studio (2022.07.2 Build 576). Below are the commands we used for the analysis:

~~~
counts <-c(174, 79, 36, 28, 23) #AMY1 copy number counts
populations <- factor(rep(c(“AFR”, “AMR”, “EAS”, “EUR”, “SAS”),
times = c(1, 1, 1, 1, 1)))
kruskal_test_result <- kruskal.test(counts, populations).
~~~

### Distribution of singleton proportions for genome-wide tandem repeat loci

To quantitatively compare the fraction of singletons in the amylase locus with other fast-evolving loci, we downloaded genotypes of tandem repeats (TR) loci from (https://github.com/gymrek-lab/EnsembleTR), not taking into account the length of repetitive elements, and selected diploid genotypes of individuals present in both the TR dataset and our dataset (33 individuals, 66 alleles). Our study comprises 81 samples with 30 high-confidence haplotypes. Among these, 33 samples are also present in the EnsembleTR dataset. By comparing data from the same individuals, we eliminated the effect of genetic drift in our comparison. From the 33 samples, 21 out of the 30 amylase haplotypes were identified. We then selected 719 TR loci that have the same number of distinct alleles (21 alleles from 33 individuals) as our comparative data. The remaining nine alleles from the high-confidence haplotypes (n = 30) correspond to the alleles detected from samples that are not present in the TR dataset. For the selected loci, the distribution of singleton proportions was generated by dividing the number of singletons by the total number of alleles (n = 21) for each locus. The empirical p-value is represented as the proportion of TR loci with a singleton proportion greater than or equal to that observed at the amylase locus (n = 14). Note that the number of singleton haplotypes (n = 19) that we detected as part of the high-confidence haplotypes were reduced to 14 since the EnsembleTR didn’t have samples that carry the remaining five singleton haplotypes. For a comparison, we repeated this analysis for 3q29 locus, known for its segmental duplication-rich nature and high levels of structural variation similar to the amylase locus. For adequate comparison, we used the same individuals (n = 39) for whom we were able to reconstruct 3q29 haplotypes for both of the chromosomes. Haplotype information for the 3q29 locus was obtained from a previous study (*15*). In addition, we filtered the tandem repeat loci (601 loci (unit length 1-6 bp)) from EnsembleTR that we analyzed to match the number of distinct alleles (n = 22) that we observed in the 3q29 locus.

### Rarefaction analysis

Rarefaction analysis was performed using the R package called “iNEXT” (*52*) to evaluate haplotype diversity and determine whether the number of samples included sufficiently captures majority of diversity at the amylase locus. The following command was used for both all amylase haplotypes (n = 51) and four common amylase haplotypes:

~~~
Data <- read.table(“rarefaction_allPopsCombined.txt”, header =
TRUE, sep = “\t”)[6,]
Obj <- iNEXT(as.numeric(Data), datatype=“abundance”, q=0,
knots=sum(Data)*2)
Temp_data <- Obj$iNextEst$size_based
Rate <- c()
for (i in 1:nrow(Temp_data)){
     Rate[i+1] <- Temp_data[i+1,5]-Temp_data[i,5]
}
Plot_data <- data.frame(NumChr=Temp_data$m, NumHap=Temp_data$qD,
Rate=Rate[-length(Rate)])
pdf(“Sample-size-based_REcurve_with_rate_of_change.pdf”, width=6,
height=4)
     twoord.plot(Plot_data$NumChr,Plot_data$NumHap,Plot_data$Num
Chr,Plot_data$Rate, type=“l”, xlab=“Number of alleles”,
ylab=“Haplotype count”, rylab=“Rate of change”,
lytickpos=seq(0,70,10), lcol=“blue3”,rcol=“red2”)
     abline(v=sum(Data), lty=“dashed”)
dev.off()
pdf(“Coverage-based_REcurve.pdf”, width=6, height=4)
     ggiNEXT(Obj, type=2)+
          theme_bw()+
          theme(legend.position=“none”)+
          xlab(“Number of alleles”)
dev.off()
~~~

### Amylase gene annotation using protein sequence homology

To predict amylase gene coordinates and copy number on each haplotype, we used homology-based gene prediction tool (“protein2genome”) implemented in Exonerate v2.4.0 (*53*) with default settings except for maximum intron length of 20-kbp (`--maxintron 20000`). For the homology search, we used the amylase protein sequences (n = 3) with experimental evidence from UNIPROT (“Evidence at protein level”) (*54*) under the following accession numbers; *AMY2B*: P19961, *AMY2A*: P04746, *AMY1A*: P0DUB6, *AMY1B*: P0DTE7, and *AMY1C*: P0DTE8 (The protein sequences of *AMY1A*, *AMY1B*, and *AMY1C* are identical). We clustered any overlapping hits from the prediction results and selected the best hit from each cluster. The hits that can be translated into 511 full-length polypeptide sequences were then kept as a final annotation. In cases where we found conflict between the homology-based annotation and the predicted copy number of *AMY* genes from amylase haplotypes, the homology-based prediction was manually curated to ensure it to be consistent with the optical mapping label. Additionally, genome annotations of *de novo* assemblies were obtained by using the T2T-chm13 gff annotation file with liftoff (*55*).

~~~
liftoff -db $dbfile -o $outputfile.gff -u $outputfile.unmapped –
dir $outputdir -p 8 -m $minimap2dir -sc 0.85 -copies $fastafile -
cds $refassembly
~~~

The liftoff results were compared with homology-based annotation above and were used when the manual curation was not able to produce annotation consistent with the optical mapping label, where we found only one case (HG01175_H7.1; table S4). Lastly, pseudogene annotations were obtained from the liftoff annotation files (table S4).

### Validation of AMY1 copy number using digital droplet PCR

Custom primer and probe oligos were designed to target a conserved sequence within *AMY1* gene in exon 5 across all annotated amylase gene copies (avoiding pseudogenes) in the T2T-chm13 human reference genome, with the following sequences: forward primer (CCTCAACAGGTCAGAGATTGTC), reverse primer (GGCAATCTTAGAACGCACATAATC), and probe (TGGTCTTCTCGATCTTGCACTGGG). The ddPCR protocol, reference primer, and reagents used are as previously described (*4*). Human DNA samples from LCLs (Coriell Institute) of 101 individuals (table S5) were used to estimate total amylase gene copies in duplicates. Restriction enzyme *HindIII* (New England Biolabs), was used in the suggested concentration to digest DNA prior to ddPCR.

### Alignments of amylase segments and coding sequences

The alignments of each amylase segment and coding sequences (CDS) were constructed using MAFFT v7.310 (*56*) and PAGAN v1.53 (*57*) with --codon option, respectively. As an outgroup, chimpanzee sequences for each segment were prepared by searching a sequence that covers >= 30% of corresponding segment of the GRCh38 reference genome with >= 97% sequence similarity in mPanTro3 genome from the Vertebrate Genome Project (https://www.genomeark.org/t2t-all/Pan_troglodytes.html). Only one of the haplotypes (“haplotype 1”) from mPanTro3 was used to align segments. The raw alignments of each segment were summarized by averaging frequencies of major alleles (including a gap) across a 200-bp window with a 10-bp step (fig. S15A). We determined the phylogenetically informative interval (from 22,850 to 26,730 bp in the AMY1 segment alignment, Data File-S2) in the AMY1 alignment based on the change point of consensus determined by a statistical method, “AMOC” (*58*) implemented in the R package, “changepoint”, and transposable element information in the alignment. The CDSs of *AMY2B*, *AMY2A*, and *AMY1* genes for chimpanzees were retrieved from the Ensembl annotation of panTro5 (release 109) (*59*). Additionally, as an outgroup to all amylase CDSs, we incorporated amylase CDS from the sheep genome, Oar_rambouillet_v1.0 (release 109) (*59*) which is known to have only one copy of the amylase gene (*4*). We also generated an alignment of the phylogenetically informative interval of the AMY1 segment from humans and the non-human primates chimpanzee (Clint_PTRv2_panTro6), bonobo (Mhudiblu_PPA_v0_panPan3), and gorilla (Kamilah_GGO_v0_gorGor6) using MAFFT v7.310.

### Phylogenetic reconstruction of AMY1 segments and coding sequences

We used the AMY1 segment, the phylogenetically informative interval of the AMY1 segment, and CDS alignments (see “*Alignments of amylase segments and coding sequences*”) to phylogenetically cluster sequences. A maximum-likelihood tree for each alignment was reconstructed using IQ-TREE v2.2.0 (*60*) with the following options: `-m MFP` (*61*), `–alrt 1000`, `-B 1000` (*62*). The chimpanzee (mPanTro3) AMY1 segment and CDSs from chimpanzee (panTro5) and sheep (Oar_rambouillet_v1.0) amylase genes were used as outgroups for the AMY1 segment and CDS, respectively. The tree for the phylogenetically informative interval of the AMY1 segment was rooted by midpoint. All the reconstructed trees were visualized using FigTree v.1.4.4 (http://tree.bio.ed.ac.uk/software/figtree/).

### Estimation of dN/dS for amylase genes

To estimate dN/dS ratios, we used codeml implemented in PAML v4.10.6 package (*20*). First, we pruned the tree topology (fig. S11) and CDS alignment (Data File-S2) to keep only unique CDSs that are not identical to each other. Based on the pruned tree and alignment, we employed a branch model (H1), which allows the dN/dS ratio to vary among branches (model = 2, NSsites = 0, fix_omega = 0 and omega = 0.4). We estimated dN/dS ratio for each gene type by labeling the clade including all CDSs from the gene type of interest as foreground branches. To obtain statistical significance for gene type-specific selection pressure, we calculated a likelihood for a null branch model (H0_1_) where dN/dS ratio did not vary among branches (model = 0, NSsites = 0, fix_omega = 0 and omega = 0.4) and performed a χ^2^ test with the likelihood ratio between H0_1_ and H1 models. To obtain statistical significance for negative selection pressure, we calculated a likelihood for a null branch model (H0_2_) where dN/dS ratio for the foreground branches (e.g. human *AMY1* clade) is fixed to one (model = 2, NSsites = 0, fix_omega = 1 and omega = 1) and performed a χ^2^ test with the likelihood ratio between H0_2_ and H1 models. The raw p-values were adjusted by controlling false discovery rate (FDR) (*63*).

### Functional analysis for amylase protein sequences

To examine the potential functional effects associated with differences between different amylase gene types (*AMY1*, *AMY2A*, *AMY2B*), all predicted peptide amylase sequences, derived from amylase haplotypes in this study, were aligned using CLUSTALO (*64*). Amino acid differences that were present in six or more haplotypes within and between amylase sequences were then noted (Fig. S12). Locations of site differences were subsequently intersected with previously identified functional domains available through UniProtKB (Accession: P0DUB6). Further functional analysis followed using data produced from recently published AlphaMissense (J. Cheng et al. 2023), examining all possible amino acid substitutions per site across the entire amylase peptide sequence. Using this data, we examined the above noted amylase amino acid site differences and mined previously assigned functional impact scores from AlphaMissense. Mutations from AlphaMissense were described as having no effect (“likely benign”), undetermined effect (“ambiguous”), or having a functional consequence (“likely pathogenic”) (table S8).

### Detecting sequence origin and breakpoints in AMY1 segments

To explore the origin of sequences within the AMY1 segment, we first generated consensus sequences for each AMY1 segment cluster by selecting the major allele at each site in the alignment. We then prepared a pseudo reference sequence including only a single copy of each amylase segment (AMY1C segment was included as AMY1 segment) from the GRCh38 reference genome. For each 200-bp sliding window with 10 step intervals in each cluster, we searched for the best hit against the pseudo reference sequence using BLAT v. 37×1 (*65*), and determined it as the origin of the 200-bp sequence. The windows where more than or equal to 50% of the sequence is gap were excluded in this analysis. The breakpoints of possible NAHR events were manually determined around the transition of origins by using segment-specific variants (table S15). To visualize homology between the consensus sequences from each AMY1 segment cluster and the other segments in the context of haplotype, we aligned the sequences using nucmer v3.1 and generated dotplots using mummerplot v3.5 (*66*) (fig. S32).

### Dating AMY1 duplication events

To estimate the timing of *AMY1* duplication events, we first constructed a starting tree using the whole AMY1 segment alignment above (see the “Alignments of segments and coding sequences’’ section) with IQTREE v2.2.0 (*60*) and --alrt 1000 -B 1000 options. We then trimmed out the phylogenetically informative interval (from 22,850 to 26,730 bp in the AMY1 segment alignment, Data File-S2) that is potentially poorly aligned due to the presence of rearrangement breakpoints. Exonerate v2.4.0 (*53*) with maximum intron length of 20-kbp (--maxintron 20000) was used to determine coordinates of the coding region in the alignment and to ensure that all sequences in the alignment have the same coordinates for the coding region. Based on the annotation, we partitioned the alignment into four categories: non-coding region, 1st position of codon, 2nd position of codon, and 3rd position of codon. This partitioned alignment was used as an input for the downstream analyses. We extracted sequences and topology of H3^r^.1 or H3^r^.2 haplotypes and chimpanzee genome (mPanTro3) from the partitioned alignment and the starting tree, respectively. We restricted our analyses to H3^r^.1 or H3^r^.2 haplotype that is likely to be ancestral (due to their high allele frequency) to other more complex haplotypes to reduce the genetic exchange effect of inversion between different AMY1 segments.

For estimating the timing of *AMY1* gene duplication events (that is, AMY1 segment duplication events), BEAST v2.7.5 (*67*), a Bayesian framework, was used with the alignments and starting trees of H3^r^.1 and H3^r^.2 haplotypes, separately. First, we performed bModel Test implemented in the BEAST to assess whether the four partitions are evolving differently or not using unliked site models and calibrated Yule model. Based on the results of bModel Test, we decided to use average substitution models using bModel Test in the real run to incorporate heterogeneity across partitions. Then, a path sampling approach from BEAST’s MODEL_SELECTION package was used for all model combinations to determine best-fit clock and tree models. We tested both strict and optimized relaxed clock models with constant and Bayesian skyline tree models. For the path sampling, 40 path steps with 25,000,000 iterations were used with an alpha parameter of 0.3, pre-burn-in of 75,000 iterations, and an 80% burn-in of the complete chain. While the models with a strict clock model could not produce marginal likelihood estimates, there was no significant difference among the models with an optimized relaxed clock model (bayes factor = 1.32 and 0.14 for H3^r^.1 and H3^r^.2; table S9). Therefore, we estimated time for both models with Markov chain Monte Carlos (MCMC) with 75,000,000 iterations, sampling every 2,000 trees and using a burn-in of 10% iterations for each model. In all the time estimations above, MRCA prior to assuming the human-chimpanzee divergence time to 6 million years ago under normal distribution (a mean of 6 million years ago and variance of 0.5) was used to calibrate time estimates. Also, we did not allow optimization of the topology of the starting tree during the estimation to avoid shuffling between AMY1 segment clusters likely due to the observed gene conversion effect. Tracer v1.7.2 was used to examine effective sample size and convergence of estimates. Treeannotator v2.7.5 was used to merge and annotate the sampled trees into a single tree. Figtree v1.4.4 (http://tree.bio.ed.ac.uk/software/figtree/) was used for tree visualization.

### Amylase copy number detection in ancient humans

We used the program GeneToCN (*26*) to determine amylase copy number in 73 ancient human and 8 archaic hominin genomes (fig. S33 and, tables S10 and S12) (*27*, *42*, *68–92*). The raw sequencing FASTQ formatted files were obtained through the European Nucleotide Archive (https://www.ebi.ac.uk/ena/browser/). As GeneToCN is a read-depth based copy number caller, we only utilized genomes that had at least 5X coverage both overall and in the amylase region to minimize random error. Additionally, we did not use sequences whose FASTQ files were generated by duplicate-removed bam files as we consistently found that this step generated no output for the *AMY1* copy number analysis. We used the program provided databases (https://github.com/bioinfo-ut/GeneToCN/tree/main/Kmer_db/Gene_kmers/AMY) for *AMY2A*, *AMY2B*, and *AMY1* to calculate the copy number of each gene using 25 nucleotide sequences (25-mers). We additionally used GeneToCN to calculate the amylase copy numbers from 101 present-day humans (table S5). We downloaded mapped bam files for the individuals that are available in both our diploid haplotype data and 1000 Genomes Project short-read sequencing dataset from The International Genome Sample Resource (https://www.internationalgenome.org) and HPRC (https://s3-us-west-2.amazonaws.com/human-pangenomics/index.html?prefix=working/). We validated these results using digital PCR (R^2^ = 0.95 for *AMY1*) (table S5). We used the following command for the GeneToCN:

~~~
python KmerToCN.py -db AMY_all_kmers.db -kp kmer_db_locations.txt
-s ${file} -d AMY/${individual} -o ${individual} -gm
${direct}/GenomeTester4/src/ -r AMY.txt -i
~~~

Next, we compared the difference in *AMY1*, *AMY2A*, and *AMY2B* copy numbers between all ancient human genomes practicing hunter-gatherer or agricultural lifestyles using a two-sided Wilcoxon Rank Sum Test (P = 0.037 for *AMY1*) from the `stat_compare_means()` command in the ggplot2 version 3.5.1 library (**fig. S22**). We then binned our time as “**Prior** to the major shift in agriculture”, “**During** the shift to agriculture”, and “**Post** the shift to agriculture” to compare the amylase copy number over time in both globally and in European populations only (Fig. 3C, 3D and fig. S23) using `wilcox.test(alternative = “less”)` in R package “stats”. We calculated a one tailed Wilcoxon Rank test with the alternative hypothesis that the older time period has a lower amylase copy number for *AMY1*, *AMY2A*, and *AMY2B*. Hungarian samples from the site of Szólád were not included in this analysis as their large sample size in comparison to all other archaeological sites and close kinship creates a bias in the “**Post** transitional” dataset.

### Downsampling ancient human genomes to estimate coverage bias

To determine the effect of coverage on the amylase copy number estimation, we randomly downsampled eight ancient human genomes (table S11) using seqtk (v1.4-r122) (https://github.com/lh3/seqtk) for a range from 1X to 5X coverage. We calculated the *AMY1* gene copy number five times and compared these results to the calculated copy number estimated when using the full genome coverage prior to downsampling. To consider batch effects, we chose samples from seven different studies with four different corresponding authors ranging in date from 2014 to 2024 (table S12). Additionally, we chose samples that were a mix of hunter-gatherers (n = 3) and farmers (n = 5) and had a wide range of both average genome coverage (8.2-30X) and *AMY1* gene copy number (6-17 copies) to consider unknown systematic biases coming from different individuals and sequencing settings. An example version of the script for the analysis is provided below:

~~~
random_seed=$((RANDOM % 1000 + 1))
Ratio=$(echo “ scale=3; ${Downsample} / ${Coverage}” | bc -l)
seqtk sample -s seed=${random_Seed} ${Run_ID}_1.fastq.gz
${Ratio} | gzip >
${Individual}_1_downsampled_r${Round}_d${Downsample}.fastq.gz
seqtk sample -s seed=${random_Seed} ${Run_ID}_2.fastq.gz
${Ratio} | gzip >
${Individual}_2_downsampled_r${Round}_d${Downsample}.fastq.gz
coverage=“high_coverage“
sample“${Individual}_downsampled_r${Round}_d${Downsample}“
GeneToCN.sh ${sample} ${coverage} ${directory}
~~~

### Archaic hominin genome analyses

We collected sequencing raw reads of archaic hominin genomes (n = 38) from publicly available datasets (table S10), removed adaptor sequences, and merged paired-end reads in case of overlapping using leeHom v1.2.17 (*93*). To increase sensitivity, we did not trim the reads by their base quality scores. For Chagyrskaya Neanderthal genome, we used unmapped reads (http://ftp.eva.mpg.de/neandertal/Chagyrskaya/rawBAM/) that were already merged, and adapter trimmed. The clean reads were mapped to the GRCh38 reference genome (http://ftp.1000genomes.ebi.ac.uk/vol1/ftp/technical/reference/GRCh38_reference_genome/GRCh38_full_analysis_set_plus_decoy_hla.fa), which includes decoy and HLA sequences, using “bwa aln” v0.7.17 (*94*) with the following options: `-n 0.01 -o 2 -l 16500`. Next, we employed the “computeGCBias” command from the deepTools v3.5.1 (*95*) suite to assess GC bias in archaic hominin genome dataset. Then, we used the “correctGCBias” command from deepTools v3.5.1 to obtain corrected reads. This correction process is essential for minimizing potential errors due to GC bias in downstream analyses. We evaluated the GC bias in GC-corrected reads one more time and 29 out of 38 archaic hominin genomes were excluded from downstream analyses due to moderate to significant GC bias (table S10). In addition, we calculated average read-depth for the whole genome and the amylase locus using mosdepth v0.3.5 (*96*). Based on this result, we excluded one more sample (Scladina I4-A) that has lower than 1X coverage both at the genome wide level and at the amylase locus.

Following the completion of GC bias analysis, the GC-corrected reads were realigned to the T2T-chm13 and GRCh38 assemblies to enhance the accuracy of the alignment and reduce the reference bias (*97*), utilizing “bwa mem” v0.7.17-r1188 (*94*). A reference genome with a single copy of the *AMY1* gene (*10*) was used in the final alignment stage to filter out reads that aligned to non-target chromosomes. These sequential alignment and filtering steps were designed to minimize background noise and enhance the specificity of read alignment to each amylase gene. The average read depths were calculated and normalized by genome-wide depth using mosdepth v0.3.5 (*96*) and copy number of amylase genes were calculated by using this read-depth information. For visualization of read-depth around the *AMY1* gene, we extracted GC-corrected reads mapped to the amylase locus and aligned them back to the reference genome containing a single copy of the *AMY1* gene (Perry et al. 2015). The read-depth was normalized by the average read-depth of *RNPC3* gene for comparable visualization in Fig. 3A.

### Scanning PRDM9 binding sites across the H3^r^.1 haplotype

To identify potential PRDM9 binding sites, FIMO v5.5.4 implemented in the MEME package (*98*) was employed using the degenerate 13-bp motif, “CCNCCNTNNCCNC” (*99*), and the non-redundant DNA database background frequency matrix in the H3^r^.1 haplotype. We used a consensus sequence for the H3^r^.1 haplotype to incorporate the standing nucleotide variation within the locus. Only hits with a FIMO score above 10 and a p-value below 0.00011 were considered (fig. S26 and table S13).

### Investigating mechanisms underlying structural variation formation within haplotypes

To investigate putative mechanisms underlying structural variants between two haplotypes, we aligned the haplotypes using NUCmer v3.1 (*100*). The alignments were visualized as dotplots using mummerplot v3.5 and Miropeats-style plots using the R package SVbyEye (https://github.com/daewoooo/SVbyEye/tree/master). These sequence comparisons allowed us to identify the potential breakpoints of structural variants. Once we identified the potential breakpoints, we extracted 20-kbp long sequences upstream and downstream regions and then aligned them to each other, including the corresponding GRCh38 reference sequence using MAFFT v7.522 (*56*). Based on the analysis of these breakpoints, we evaluated replication and DNA recombination-based processes that underlie the mutational steps leading to the extant structural variants. We inferred NAHR when the variations between two structural haplotypes included (i) breakpoints within paralogous segments and (ii) absence of sequence motifs linked to replication-based mutational mechanisms.

## Acknowledgements

We thank Sabriya Syed, Valerie Reyes-Ortiz, Nicholas Moskwa, Pille Hallast, Carmen Robinett, Stefan Ruhl, Victor Albert, Vincent Lynch, Derek Taylor, Leo Speidel, and John Novembre for technical help, discussions, suggestions, and feedback on the manuscript; the Human Genome Structural Variation Consortium and the Human Pangenome Reference Consortium for making their data publicly available; The Jackson Laboratory Scientific Services, including the Genome Technologies Service for expert assistance with the work described herein, and Research IT for computational infrastructure and support. We are grateful to the people who generously contributed their samples to the 1000 Genomes Project.

## Authors Contributions

O.G. and C.L. conceived the study. F.Y. performed the analysis and interpretation of the Human Genome Structural Variation Consortium (HGSVC), and Human Pangenome Reference Consortium (HPRC) Bionano Genomics optical mapping, PacBio HiFi sequencing and phased assemblies, performed haplotype-resolved assemblies of HGSVC PacBio HiFi samples using hifiasm and amylase haplotype detection using HGSVC, and HPRC datasets. C.K. performed the mutational mechanisms analyses, breakpoint characterization, linkage disequilibrium, PCA and phylogenetic analyses. C.K. contributed to evolutionary and functional analysis. K.K. performed gene annotation, singleton analysis, selection/phylogenetic analyses of amylase coding sequences/segments, and interpretation/time estimation of initial *AMY1* gene duplication events. F.Y. and K.K. performed archaic hominin genome processing. P.P. performed ddPCR validation experiments, analysis of functional site differences in amylase amino acid sequences, and data visualization. K.S. calculated amylase gene copy numbers and performed downsampling in ancient human genomes. F.Y., C.K., K.K., P.P., C.R.B., A-M.T., C.L. and O.G. drafted and critically revised the article. Final approval of the version to be published was given by F.Y., C.K., K.K., P.P., K.S., C.R.B., A-M.T, C.L. and O.G. All authors read and approved the final manuscript.

