## Supplementary Materials for "Paleolithic Gene Duplications Primed Adaptive Evolution of Human Amylase Locus Upon Agriculture"

** Contributed equally; ** Correspondence*

**The file includes:**

Supplementary Results

Supplementary Materials and Methods

Figs. S1 to S40

Tables S1 to S15

References

**Other Supplementary Materials for this manuscript include the following:**

Data Files S1 to S2

**Table of Contents**

**Supplementary Results** 3

*AMY1 and AMY2 copy number variations* 3

*Reconstruction of mutational connections among extant haplotypes* 3

*The emergence of haplotypes harboring five AMY1 copies* 5

*Variation among haplotypes harboring five AMY1 copies shaped by NAHR events* 6

*Emergence of haplotypes generated by microhomology-mediated break-induced recombination based mechanism from common haplotypes* 6

*Incorporation of haplotypes that require intermediate state into the network* 7

**Supplementary Methods** 8

*Fst calculation* ……8

*Pairwise alignments* 8

*Genomic rearrangement analysis* 8

*Visualization of alignments* 9

*Breakpoint junction characterization* 9

*Linkage disequilibrium estimation* 10

*Phylogenetic trees* 11

*Principal component analysis* 11

*Nucleotide diversity estimation* 12

*Tissue expression data analyses* 12

**Supplementary Figures** 14

**References** 62

### Supplementary Results

#### *AMY1 and AMY2 copy number variations*

Consistent with previous findings, the diploid *AMY1* copy number (n = 36 samples), ranged from 2-10; even copy numbers were more prevalent than odd copy numbers in diploid cells (fig. S29A) (*1*–*3*). In addition, while the copy number of *AMY2A* in diploid samples ranged from one to four, most samples (n = 27/36) harbored two *AMY2A* copies (fig. S29B). Similarly, the copy number of *AMY2B* in diploid samples ranged from two to three, and most samples (n = 30/36) carried two *AMY2B* copies (fig. S29B). Although variations in *AMY1* gene copy numbers are observed among amylase structural haplotypes across diverse populations, these copy number variations do not exhibit a discernible geographic specificity (p-value = 0.406, Kruskal-Wallis rank sum test).

#### *Reconstruction of mutational connections among extant haplotypes*

We constructed the mutational connections between extant haplotypes in a stepwise fashion starting from the common haplotypes *e*.g. $H1^a$.1, $H3^r$.1, $H3^r$.2, and $H3^r$.4, which make up for 70% of the haplotypes detected in our dataset. Common haplotypes provide a reasonable starting point as they are older and more likely to seed subsequent mutations (*4*).

***Emergence of $H1^a$.1:*** When compared to other common haplotypes, $H1^a$.1 differs from them by the absence of AMY1A-AMY1B segments (fig. S34). The boundaries of this deletion lie within two near identical AMY2A.2 segments with 99.82% similarity. Thus, we argue that an NAHR-driven deletion may explain the emergence of $H1^a$.1 from a haplotype that harbors these two duplications. We base this hypothesis on two observations. First, we did not find any evidence for microhomology-mediated break-induced recombination involving these rearrangement breakpoints among $H1^a$.1 and other common haplotypes. Second, we showed in the main text that haplotypes that harbor three AMY1 segments have already existed before out-of-Africa migrations, consistent with previous findings (*5*) (Fig. 3, B and C).

***Inversions among H3$^{r}$.1-H3$^{r}$.2-H3$^{r}$.4:*** The dot plots among H3$^{r}$.1, H3$^{r}$.2 and H3$^{r}$.4 show that a single inversion event can explain the structural differences among these haplotypes. Thus, a parsimonious explanation is that H3$^{r}$.2 is an intermediate haplotype between H3$^{r}$.1 and H3$^{r}$.4 (fig. S35**)**. The breakpoints between H3$^{r}$.1 and H3$^{r}$.2 involve two highly similar inverted segments that would serve as the substrates for NAHR (99.97%-100% identity and 4,355-bp in size) (fig. S30). The breakpoints between H3$^{r}$.2 and H3$^{r}$.4 involve inverted > 15-kbp segments spanning red (partially) and orange segments (fig. S35).

***Non-allelic homologous recombination driven inversion and copy number variation events that involve non-common haplotypes:*** To incorporate the haplotypes that are not explained with a single mutational step starting from common haplotypes into the connections described above, we searched two non-common haplotypes that are separated by single inversion or copy number differences with breakpoints in duplicated segments: two inverted segments leading to an inversion and two duplicated segments in the same direction leading to copy number variation (e.g. Fig. 4B). With this, we compared haplotypes to each other using pairwise dotplots to identify inversion or single copy number gain/loss events that involve two breakpoints overlapping duplicated segments. This allowed us to identify several putative mutational mechanisms (fig. S36). By combining all the mutational connections described above, we constructed a putative mutational network (fig. S27). The resulting network, despite being not definitive, provides a useful starting point for understanding the main mechanisms leading to haplotypic diversity in our amylase haplotypes.

***The emergence of haplotypes harboring five AMY1 copies***

The NAHR events between "three-copy" H3$^r$.1-H3$^r$.1 (Fig. 4B), H3$^r$1-H3$^r$.2, and H3$^r$.1-H3$^r$.4 haplotypes lead to the formation of "five-copy" H5.3, H5.4, and H5.5 haplotypes as well as H1$^a$.1 haplotype, respectively. These five-copy haplotypes differ from each other in a single inverted segment with breakpoints localizing to AMY2A.2 and intergenic.2 segments (fig. S37). This suggests an NAHR mechanism that involves inverted segments in common "three-copy" haplotypes (Fig. S28D) may also underlie the mutational steps between H5.3, H5.4, and H5.5. In other words, the initial emergence of "five-copy" haplotypes likely involves an NAHR-based copy number change from a "three-copy" haplotype. In parallel, once a "five-copy" haplotype has emerged, different structural types can readily form through a single inversion event from this "five-copy" haplotype. With this, we searched for other haplotypes in our dataset that cannot be explained by a single step NAHR duplication from common "three-copy" haplotypes but are one inversion away from existing "five-copy" haplotypes. We found that all "five-copy" haplotypes except for H5.7 and H5.9 can be explained by NAHR-driven inversions from extant haplotypes involving inverted intergenic.2 or AMY2A.2 segments (fig. S38). It is of note that a similar inversion event from H3$^r$.4 involving inverted intergenic.2 segments would generate H3$^r$.5. Adding these insights into our broader mutational network led to a relatively complete picture of haplotypes with varying numbers of *AMY1* gene copies (fig. S27).

***Variation among haplotypes harboring five AMY1 copies shaped by NAHR events***

Nine out of thirty structural haplotypes uncovered include five AMY1 segments (H5.1-H5.9) and do not vary in the copy number of individual segments, with all these haplotypes separated only by inverted sequences (Supplementary Methods). The structural rearrangements were identified using NUCmer alignments and further validated with GRIMM (*6*), which determines the minimum number of steps required for the formation of a haplotype involving inversions. One plausible mechanism to explain these observations is NAHR-mediated inversions, as we described for H3$^{r}$.1, H3$^{r}$.2, and H3$^{r}$.4 (fig. S35). For example, all the inversion breakpoints involving these five copy haplotypes involve inversions mediated by the AMY2A.2 and intergenic.2 segments (figs. S37 and S38). The other haplotype, H5.10 harboring five AMY1 segments, likely evolved through a different mechanism (fig. S36E).

***Emergence of haplotypes generated by microhomology-mediated break-induced recombination based mechanism from common haplotypes***

The microhomology-mediated break-induced recombination is a replication-based mechanism that provides a mechanistic framework to explain the formation of complex non-recurrent rearrangements and microhomologies spanning from 2 to 15-bp can be detected at the breakpoint junctions (*7*). Thus, we scanned amylase haplotypes for microhomologies at the breakpoint junctions that underlie structural variants between common haplotypes and haplotypes that could not be generated via NAHR events. We detected three haplotypes that fit this criterion: H2A2B2.1, H3$^{r}$.6 and H3B2.1 (fig. S39). Next, we pinpointed the specific junctions where the replication fork is likely to stall and reconstructed the potential mechanisms through which H2A2B2.1, H3$^{r}$.6, and H3B2.1 could evolve from H3$^{r}$.1, H3$^{r}$.2, and H3$^{r}$.5, respectively (Fig. 4C and fig. S27). Although these arrangements constitute only 4% of the haplotypes analyzed, they are particularly interesting because they underlie the copy number variation of *AMY2* genes. Specifically, the formation of H2A2B2.1 leads to the duplication of both *AMY2A* and *AMY2B*, whereas the formation of H3B2.1 results in the duplication of *AMY2B*. By integrating these putative microhomology-mediated break-induced recombination mutational steps, our mutational network expands to include haplotypes with *AMY2* copy number variation (fig. S27).

***Incorporation of haplotypes that require intermediate state into the network***

Following the approach described in "*Investigating mechanisms underlying structural variation formation within haplotypes*" in the Material and Methods section, we were able to identify putative mutational steps for all but three haplotypes: H4A0.1, H4A0.2, and H4A2.1. In all our pairwise comparisons, we found more than two breakpoints and thus cannot explain the formation of these haplotypes without invoking an intermediate haplotype. Given the small sample size of our study, it is likely that these intermediate haplotypes exist but were not detected. Thus, we integrated these haplotypes into our broader mutational network by assuming the existence of intermediate haplotypes. Specifically, we can attribute the formation of H4A0.1 to an NAHR event involving an intermediate haplotype (H5.8''), which differs from H5.8 by two inversion polymorphisms. The inverted intergenic.2 segments in H4A0.1 likely facilitate NAHR-driven inversion, leading to the formation of H4A0.2. Similarly, H4A2.1 is closest to H4A2.2, but they are still differentiated by two distinct inverted segments. Therefore, an intermediate haplotype containing one of those inversions could explain the mutational link between H4A2.2 and H4A2.1.

### Supplementary Methods

***Fst calculation***

We assessed genetic differentiation among populations using the fixation index (Fst) calculated from haplotype allele frequencies. Haplotype frequencies were estimated for each population separately (fig. S5) and for the combined dataset across all populations. We computed within-population expected heterozygosity (Hs) and the total expected heterozygosity across all populations (Ht). The Fst values were then derived using the formula Fst = (Ht - Hs) / Ht, which measures the proportion of genetic variance attributable to differences among populations.

***Pairwise alignment***

To compare the organization of the amylase haplotypes, we aligned haplotypes from 117 alleles to the GRCh38 reference genome and the chimpanzee genome (panTro6) using NUCmer v3.1 (*8*). Additional pairwise sequence alignments for the 30 distinct amylase haplotypes were generated using two algorithms: NUCmer from the MUMmer package (version 3.1) (*9*) and minimap2 (version 2.26-r1175) (*10*). NUCmer was employed with default parameters to identify and align conserved regions among the haplotypes and minimap2 was used to enhance the accuracy of the alignments using the asm5 flag.

***Genomic rearrangement analysis***

To investigate the evolutionary path of rearrangements, the GRIMM software (version 2.01) (*6*) was utilized. GRIMM enables the prediction of hypothetical paths inferring rearrangements for the evolution of genomic regions. We utilized the NUCmer-defined syntenic blocks across haplotypes as an input into GRIMM and used default parameters to predict potential evolutionary scenarios for the haplotype set of interest.

***Visualization of alignments***

Graphical representations of the genomic alignments and evolutionary paths were generated using Miropeats-style plots using the R package SVbyEye kindly provided by David Porubsky (https://github.com/daewoooo/SVbyEye/tree/master). Alignment dotplots were created with MUMmerplot (*8*) and paf2dotplot (https://github.com/moold/paf2dotplot) to detect rearrangements and structural variant formation mechanisms and to illustrate predicted rearrangement paths.

***Breakpoint junction characterization***

The breakpoint junctions were detected using MAFFT alignments (*11*), and comparative analyses of their flanking sequences were carried out to determine the mutational mechanisms of the rearrangements. For instance, for inversion polymorphisms, the proximal breakpoint junction was assigned as +A|+B and the distal junction as +C|+D for the uninverted state, while the inverted state as +A|−C and −B|+D for the proximal and distal breakpoint junctions, respectively (fig. S30). Each of the uninverted breakpoint junctions was aligned to the inverted breakpoint junction with MAFFT (v7.520) to determine positional homologies. These sequence comparisons allowed us to identify the breakpoints of structural variants. Based on the sequence composition of these breakpoints, we considered replication and DNA recombination-based processes as described in Carvalho and Lupski (Carvalho and Lupski, 2016) that underlie the mutational steps leading to the extant structural variation.

***Linkage disequilibrium estimation***

To investigate the linkage disequilibrium around the amylase region, we extracted sequences from 100-kbp upstream of the AMY2B segment and 100-kbp downstream of the AMY1C segment from the GRCh38 reference assembly. Using minimap2 (version 2.26-r1175) (*10*), we recovered the corresponding sequences from our haplotype-resolved assemblies. The flanking sequences were aligned separately using MAFFT (v7.520) (*11*) with the parameters '--auto --adjustdirection'. These alignments were then converted from multiple sequence alignment (MSA) format to variant call format (VCF) using MsaToVcf (*12*). The VCF coordinates for the two newly obtained VCF files were adjusted using command-line tools to reflect their actual positions on the GRCh38 chromosome 1 and were merged into a single VCF file using bcftools (v1.9) (*13*). We used VCFtools (v0.1.16) (*14*) to filter out indels and multiallelic SNPs, and keep SNPs with a minor allele frequency >= 5% using the options '--remove-indels --min-alleles 2 --max-alleles 2 --maf 0.05 --recode --recode-INFO-all'. The pairwise linkage disequilibrium (LD) value was estimated using PLINK (v1.90b6.21) (*15*) with the following parameters '--r2 dprime --ld-window 999999 --ld-window-kb 10000 --ld-window-r2 0'. Linkage disequilibrium estimates $R^2$ for the flanking regions around the amylase locus were visualized using the LDheatmap package (*16*) in R. We further calculated the physical distances between the SNP pairs used for LD estimation by determining the absolute differences in base pair positions between the first (BP_A) and second (BP_B) SNPs, using the PLINK output file. The LD decay across the locus was also estimated by categorizing the data into ten intervals between the first and second SNPs, measured in base pairs. These intervals were defined as follows: 0-1,000 bp, 1,001-10,000 bp, 10,001-50,000 bp, and intervals of 50,000 bp beyond the 10,001 bp threshold. The intervals between 100,001-150,000 bp and 150,001-200,000 bp remain empty because we were investigating the upstream and downstream flanking regions, and excluding the amylase locus. This analysis was independently conducted for each common structural haplotype (H1.1, H3r.1,

H3r.2, and H3r.4) (fig. S40) to explore the LD landscape and decay across the amylase locus for each haplotype. The same parameters were used throughout, with the addition of the '--keep' flag in VCFtools during the filtering process to retain the samples carrying specific haplotypes. The LD decay for the four common structural haplotypes was visualized as ridge plots in R using ggplot (v 3.5.0) for the aforementioned distance intervals.

#### ***Phylogenetic trees***

Phylogenetic trees were reconstructed for both the 50-kbp upstream and downstream of the amylase locus using sequences from haplotype-resolved assemblies. The syntenic chimpanzee sequences were mapped and extracted from the panTro6 assembly. The 50-kbp upstream and downstream sequences from the chimpanzee and humans were aligned separately using MAFFT (v7.520) (*11*) with the parameters '--auto --adjustdirection'. Maximum-likelihood trees for each of the two alignments were then reconstructed using IQ-TREE v2.2.0 (*17*) with the options: -m TEST -bb 1000 -alrt 1000. The trees were visualized as phylograms in R using ggtree (v 3.10.1) (*18*).

#### ***Principal component analysis***

Principal Component Analysis (PCA) was performed using the same sequences that were employed in reconstructing the phylogenetic trees for both 50-kbp upstream and downstream regions of the amylase locus. First, the multiple sequence alignments (MSAs) obtained via MAFFT (v7.520) (*11*) were processed using MsaToVcf (*12*) to transform these MSAs into VCF formats. Following the transformation, we only kept biallelic SNPs with a minor allele frequency >= 5% using VCFtools (v0.1.16) (*14*) with the following VCFtools parameters: '--remove-indels --min-alleles 2 --max-alleles 2 --maf 0.05 --recode --recode-INFO-all’. The PCA was carried out using PLINK (v1.90b6.21) (*15*) and the results were visualized using R and the ggplot (v 3.5.0) R-package.

***Nucleotide diversity estimation***

To estimate the nucleotide diversity (π), we first aligned sequences from each structural haplotype separately. Specifically, alignments were conducted for 19 alleles from H1$^{a}$.1, 18 alleles from H3$^{r}$.1, 23 alleles from H3$^{r}$.2, and 21 alleles from H3$^{r}$.4. After alignment, each alignment was converted from multiple sequence alignments (MSAs) files into VCF files using MsaToVcf (*12*). We then used VCFtools (v0.1.16) (*14*) to calculate the pi nucleotide diversity across the entire region for each haplotype. Additionally, we performed the same analysis separately for African and non-African samples using the ‘--keep’ flag in VCFtools to assess nucleotide diversity within and outside of Africa.

***Tissue expression data analyses***

The expression data of amylase genes were downloaded from Genotype-Tissue Expression (GTEx) portal (*19*), representing 54 non-diseased tissue sites collected mostly from deceased people. We directly used the  TPM (transcript per million) data provided by the portal (https://gtexportal.org/home/downloads/adult-gtex/overview) to visualize gene expressions in three tissues where the highest values are observed. Since major salivary glands were not included in the GTEx portal, we used the TPM data available as a supplementary table in Saitou et al. (2020) (*20*), which includes adult major salivary gland transcriptomes. Note that even though the TPM data used here included *AMY1A, AMY1B,* and *AMY1C* transcripts as independent genes, our genomic analysis suggests that they cannot be easily discriminated against. Thus, the collective expression of these three transcripts should be considered representative of *AMY1* expression in each tissue.

### Supplementary Figures

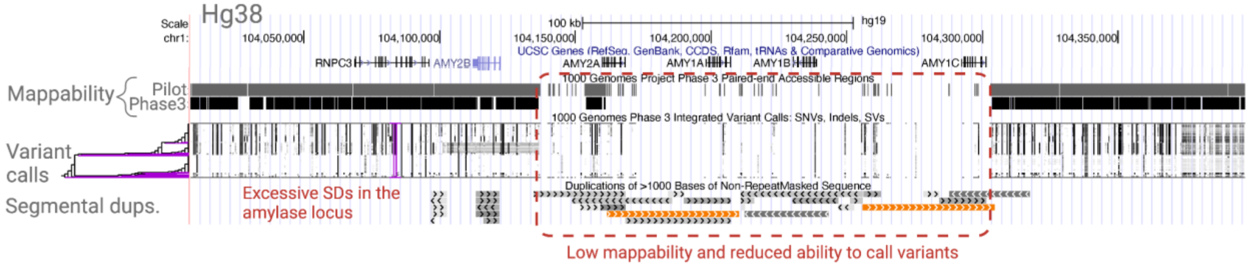


**Fig. S1. A UCSC genome browser view of the human amylase locus.**
This figure shows the mappability status from short-read sequencing data and variant calls from the 1000 Genomes Project, along with the segmental duplication track. The mappability, variant call, and segmental duplication tracks are all available on the UCSC Genome Browser. The default settings were used to visualize.

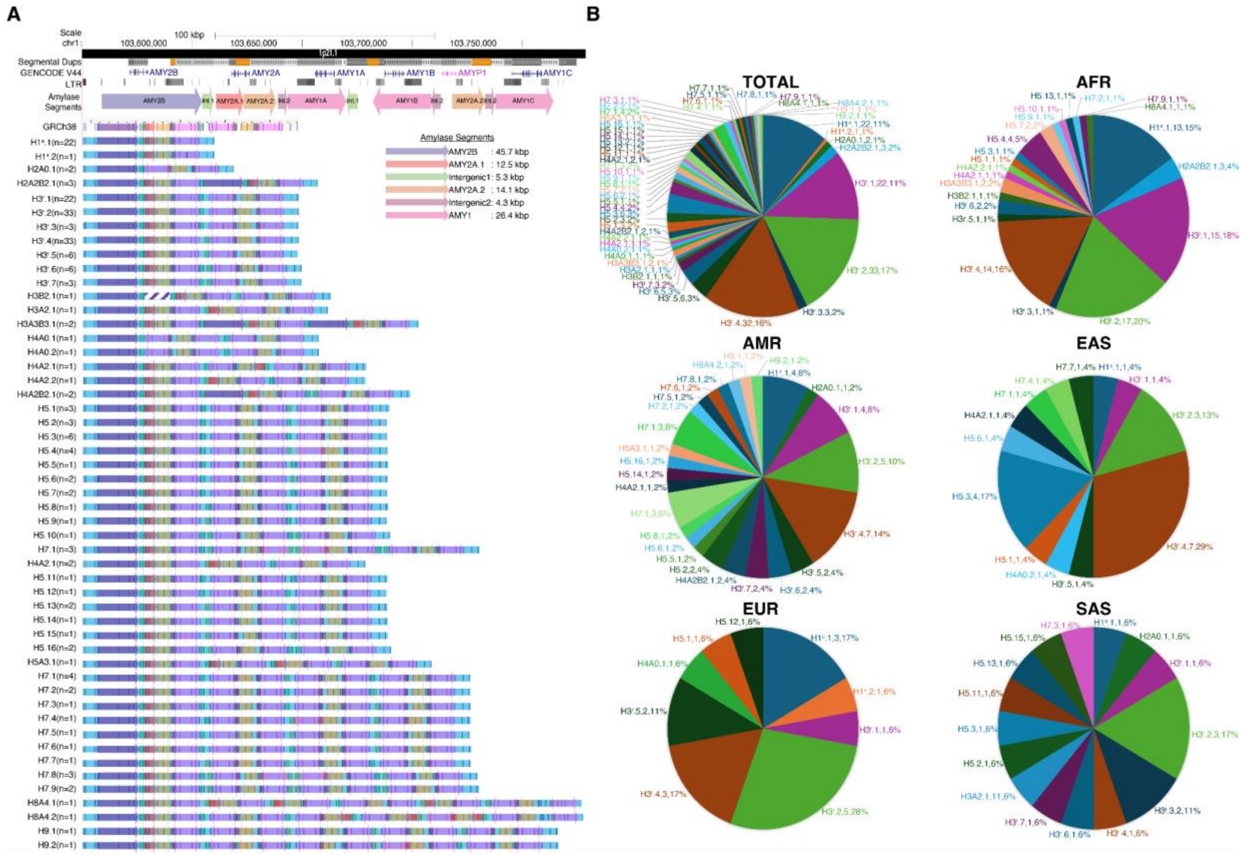


**Fig. S2. Structural amylase haplotypes detected in the human amylase locus.** (A) The top panel represents the positions of segmental duplications, GENCODE V44 gene annotations, Long-Terminal Repeats (LTRs), and amylase segments (colored arrows). The GRCh38 reference assembly haplotype structures are presented as *in silico* maps within rectangles with a white background and vertical blue lines. Amylase haplotypes (n= 51), H1ᵃ.1 - H9.2, are depicted with colored arrows indicating amylase segments and the distinct structure of each haplotype. A black line in the second AMY1 segment of H1ᵃ.2 represents the polymorphic label present in three alleles. Diagonal stripes in the second AMY2B segment of H3B2.1 indicate that the segment is a partial copy of an AMY2B segment. (B) Distribution is represented in consolidated sets of samples (TOTAL) and across populations (African - AFR, American - AMR, East Asian - EAS, European - EUR, South Asian - SAS). Nomenclature in pie charts is as follows: Haplotype ID, number of alleles, allele frequency within each continental population.

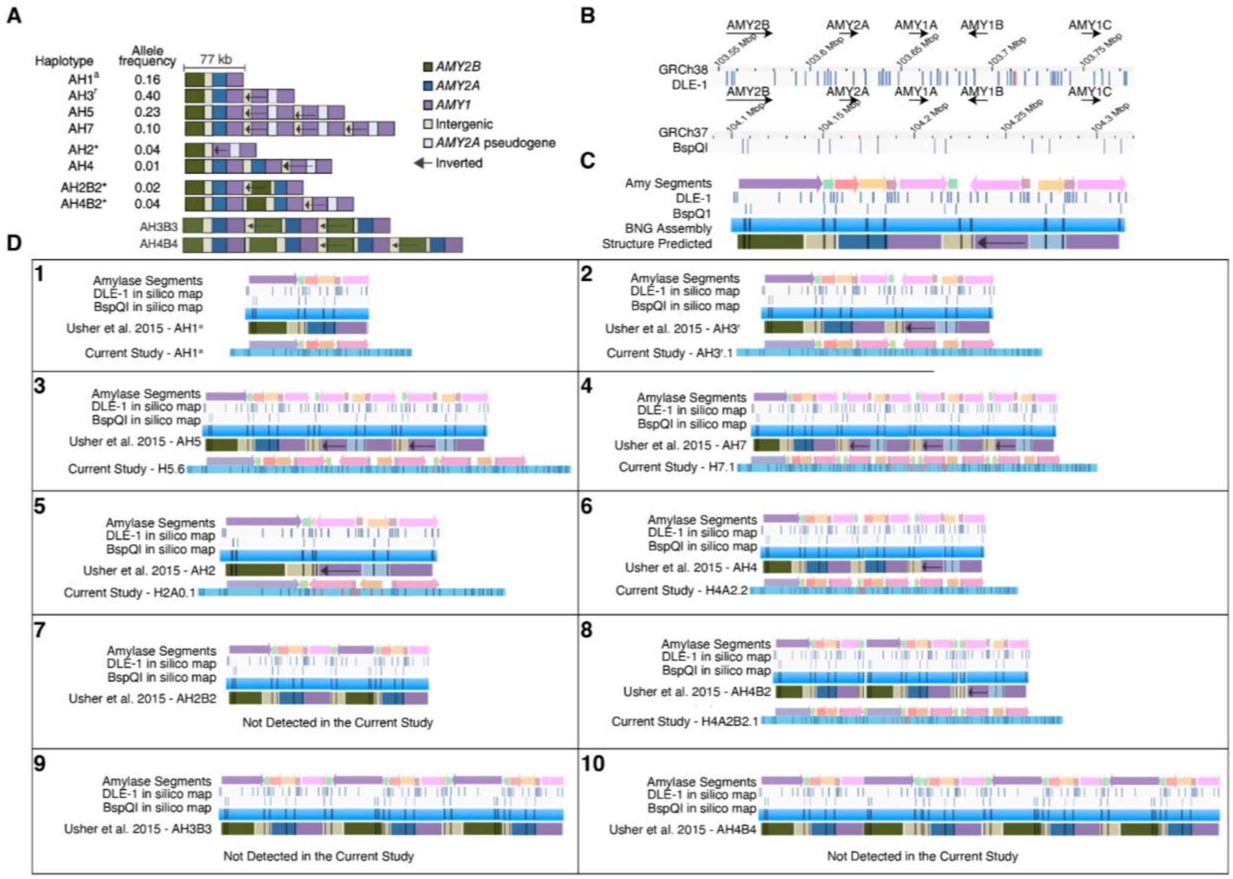


**Fig. S3. Comparison of the amylase structural haplotypes from our study with those from a previous study.**
(A) The amylase structural haplotypes (n = 10) detected previously by Usher and colleagues (*3*). (B) and (C) represent the labeling pattern of the amylase locus in GRCh38 and GRCh37 human reference genome assemblies *in silico* maps with BspQ1 and DLE-1 enzymes, respectively, generated as part of the optical genome mapping data analysis. (D) Pairwise comparison of the amylase structural haplotypes detected by Usher and colleagues with those detected in our study.

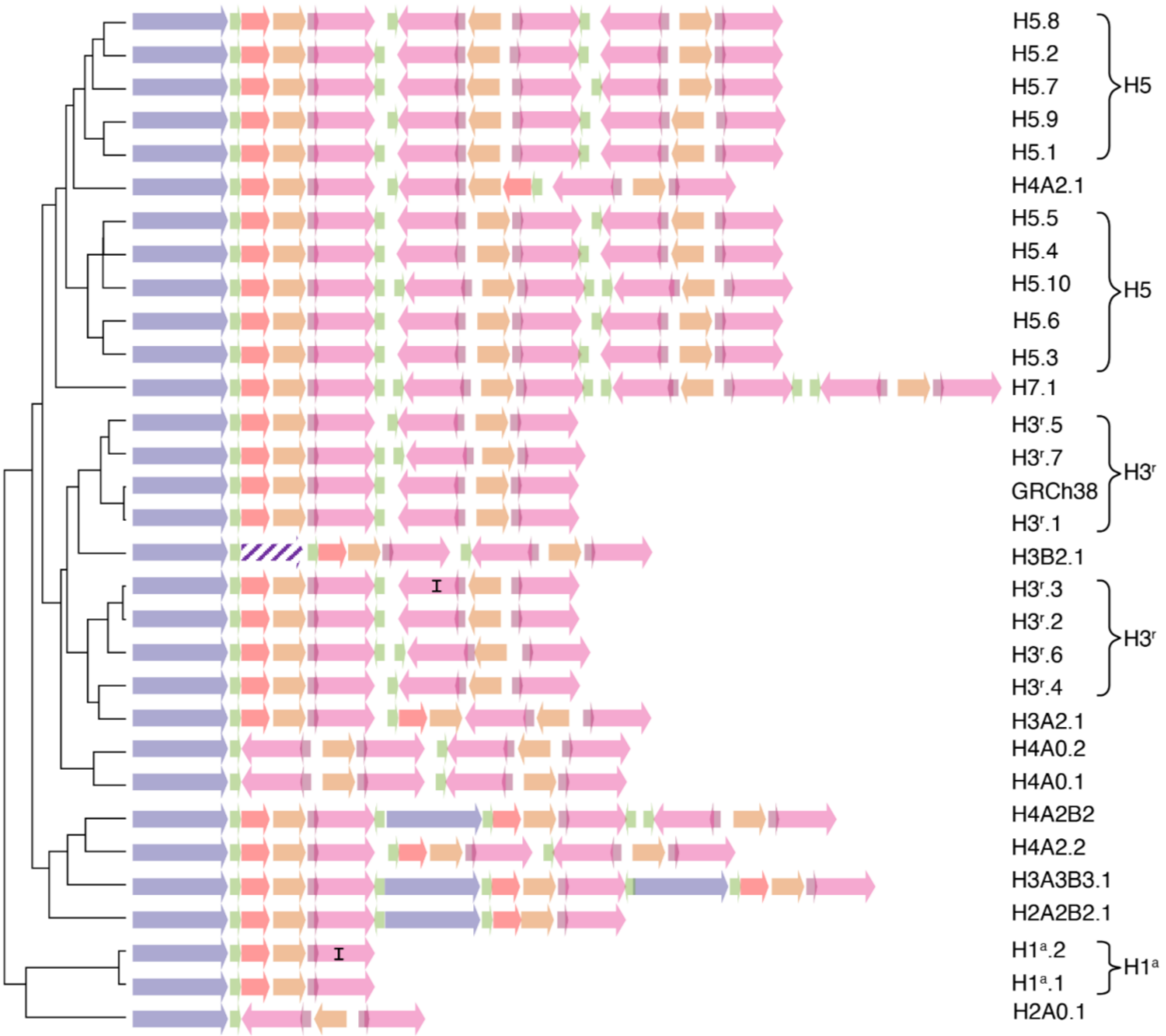


**Fig. S4. Amylase structural haplotype clusters.**
These clusters were generated based on a tree constructed using the Jaccard similarity index, as part of the pgr-tk v0.5.1 package (*21*). Haplotype IDs are displayed on the right side of the figure.

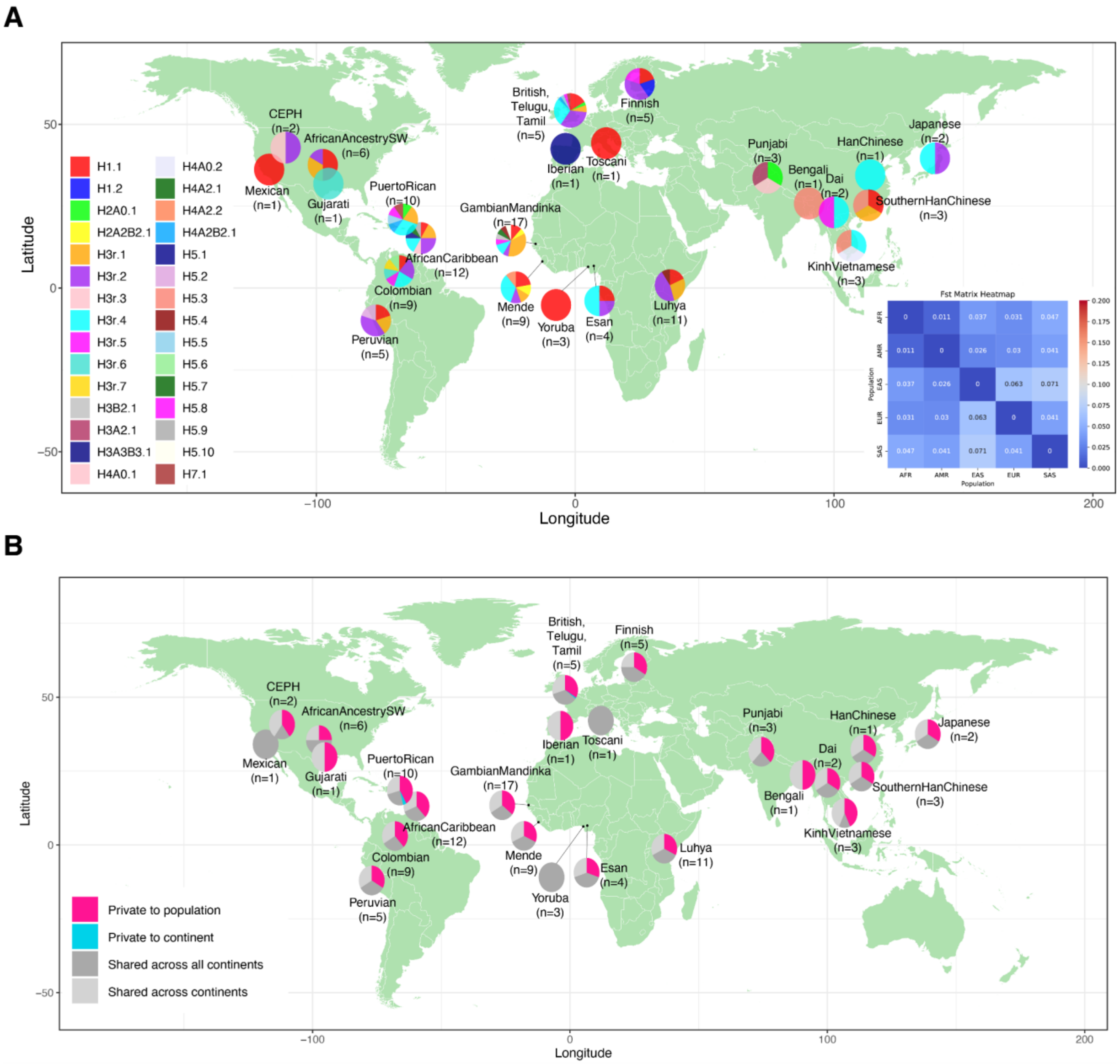


**Fig. S5. Distribution of amylase haplotypes across different continental populations.** (A) Geographical distribution of high-confidence haplotypes (n = 30). The inset heatmap shows the Fst values across populations based on the frequency of amylase haplotypes. (B) Pies are divided into four slices, representing haplotypes private to a population (pink), private to a continental area (brown), shared across continental areas (light gray), and shared across all continents (dark gray).

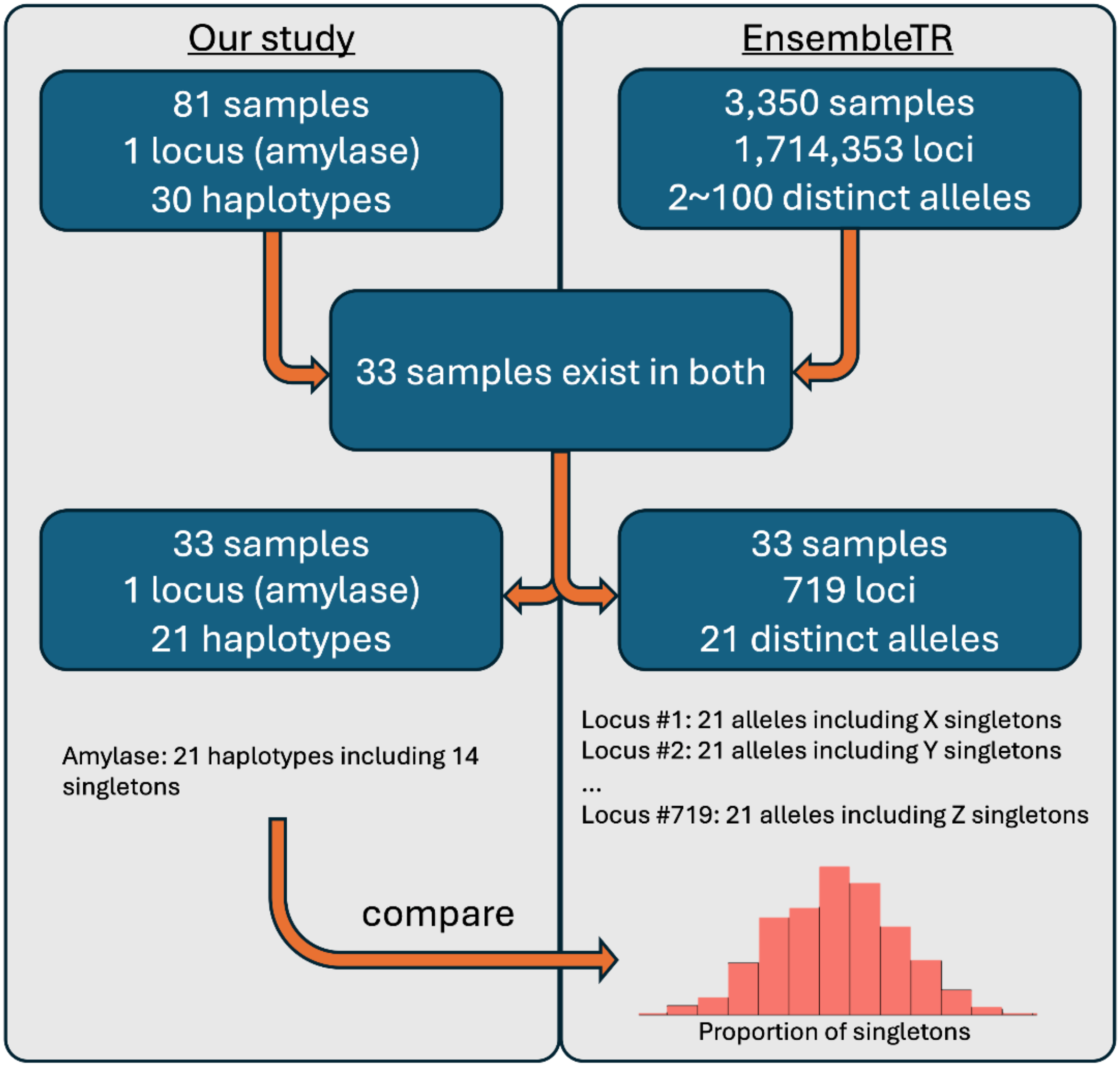


**Fig. S6. Overview of the integration and analysis of EnsembleTR samples in the study.** The flowchart details the relationship and overlap between our study's samples and the EnsembleTR dataset, leading to the identification and analysis of specific tandem repeat loci.

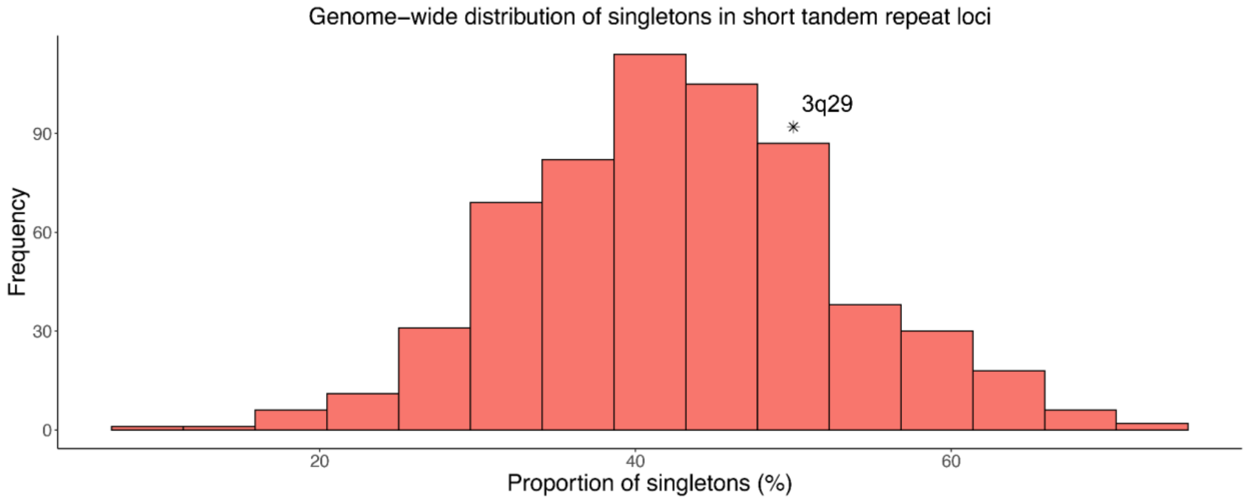


**Fig. S7. The proportion of singletons for tandem repeat loci across the genome and the 3q29 locus**

The asterisk (*) represents the proportion of singletons among all distinct alleles (50%; 11 out of 22) detected at the 3q29 locus.

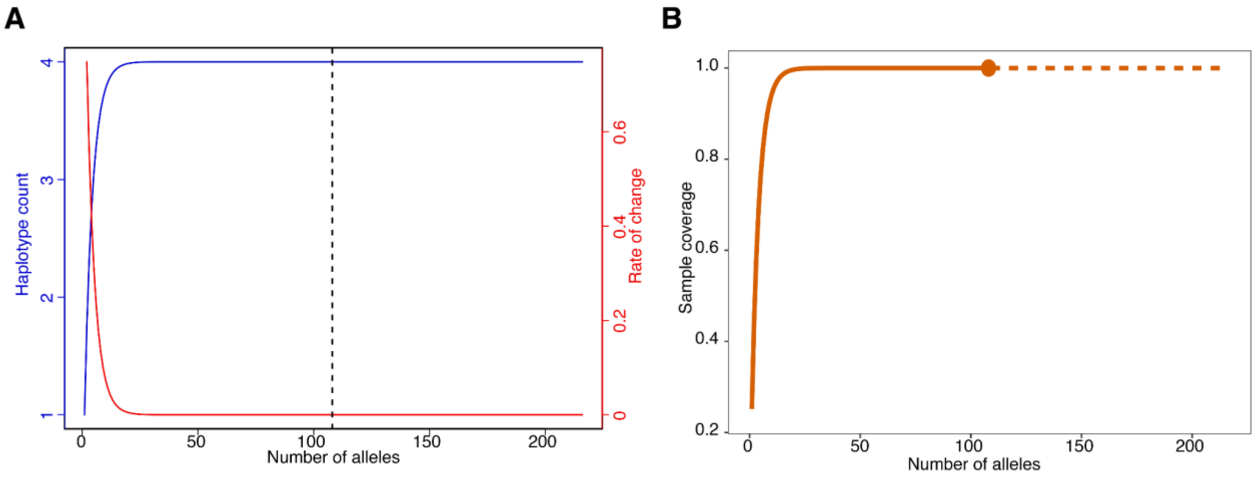


**Fig. S8. Rarefaction and extrapolation sampling curve based on four common amylase haplotypes.**
(A) Displaying how the rate of change (red line) decreases as the number of distinct haplotypes (blue line) gets closer to an asymptote with the increase in the number of alleles. The rate of change indicates the number of novel haplotypes discovered per unit increase in the number of analyzed alleles. The dashed line shows our data point with the rate of change at 1.33e-15. (B) Sample completeness curve based on four common amylase haplotypes. The dot indicates that 100% of samples belong to one of four common haplotypes in our dataset.

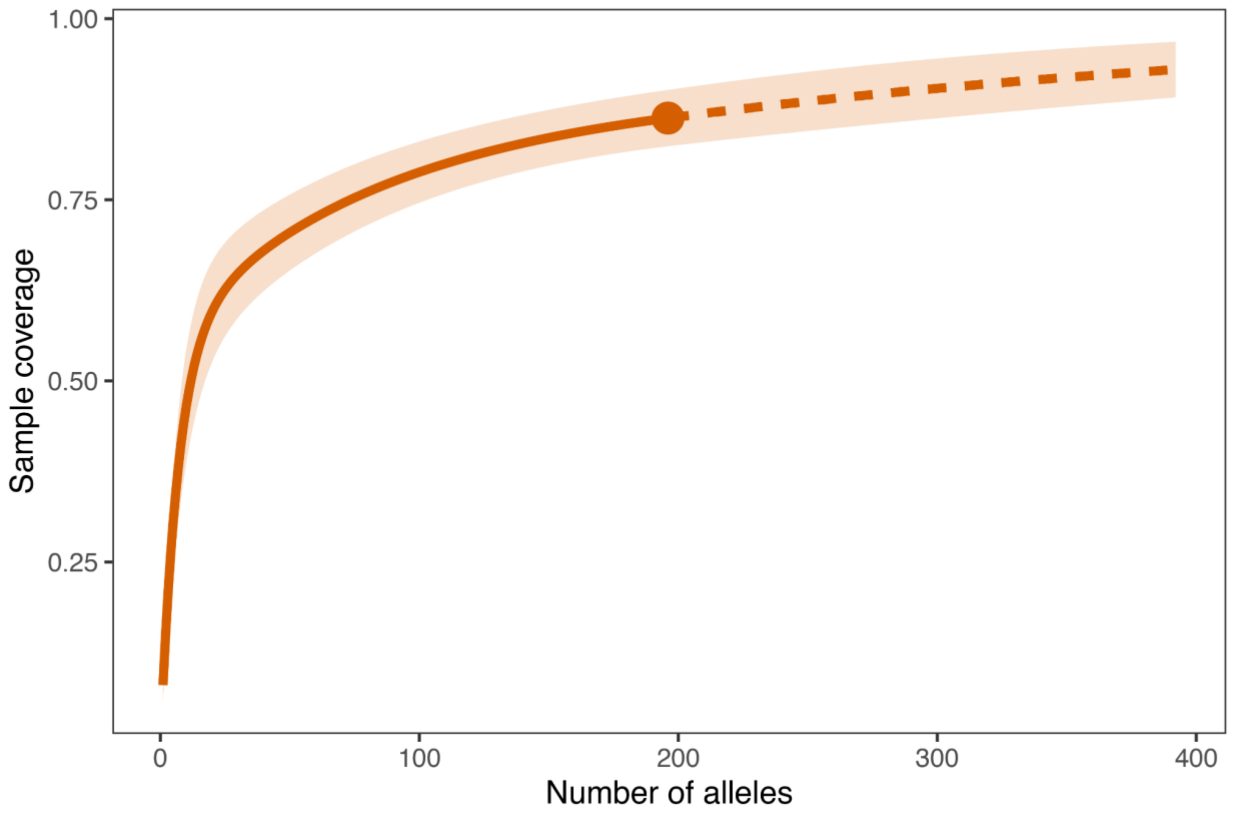

**Fig. S9. Sample completeness curve based on 51 amylase haplotypes.**
The dot indicates that 86.27% of samples belong to one of the 51 haplotypes in our dataset. The shaded area represents the 95% lower and upper confidence limits.

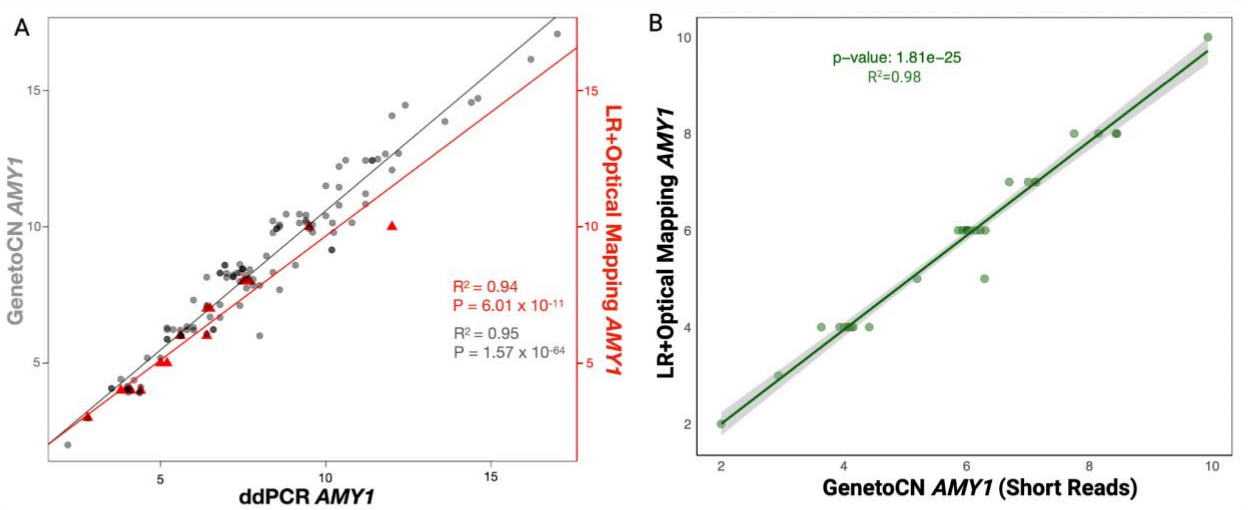


**Fig. S10. Validation of *AMY1* gene copy numbers.**
(A) Comparison of the *AMY1* gene copy numbers from long-read (LR) and optical genome mapping-based annotation predictions (red triangles, 18 individuals) and read-depth based methods, GeneToCN (gray dots, 101 individuals) (*22*), using digital droplet PCR (ddPCR). The x-axis represents the mean ddPCR *AMY1* gene copy number estimations (two rounds each), excluding pseudogenes (duplicate replicates). The right-hand side y-axis (red) represents *AMY1* gene copy number predictions from LR and optical genome mapping techniques. The left-hand side y-axis (gray) represents *AMY1* gene copy number predictions from short-read, read-depth

based (GeneToCN) techniques. P-values for the significance of correlation and linear regression R-squared values are provided in red and gray, respectively. (B) *AMY1* copy number detection using GeneToCN for samples sequenced with both short reads (x-axis) and long reads (y-axis) for 32 individuals that have their amylase copy number supported by both long-read sequencing and optical genome mapping. The p-value for the significance of correlation and linear regression R-squared values are provided in green.

444

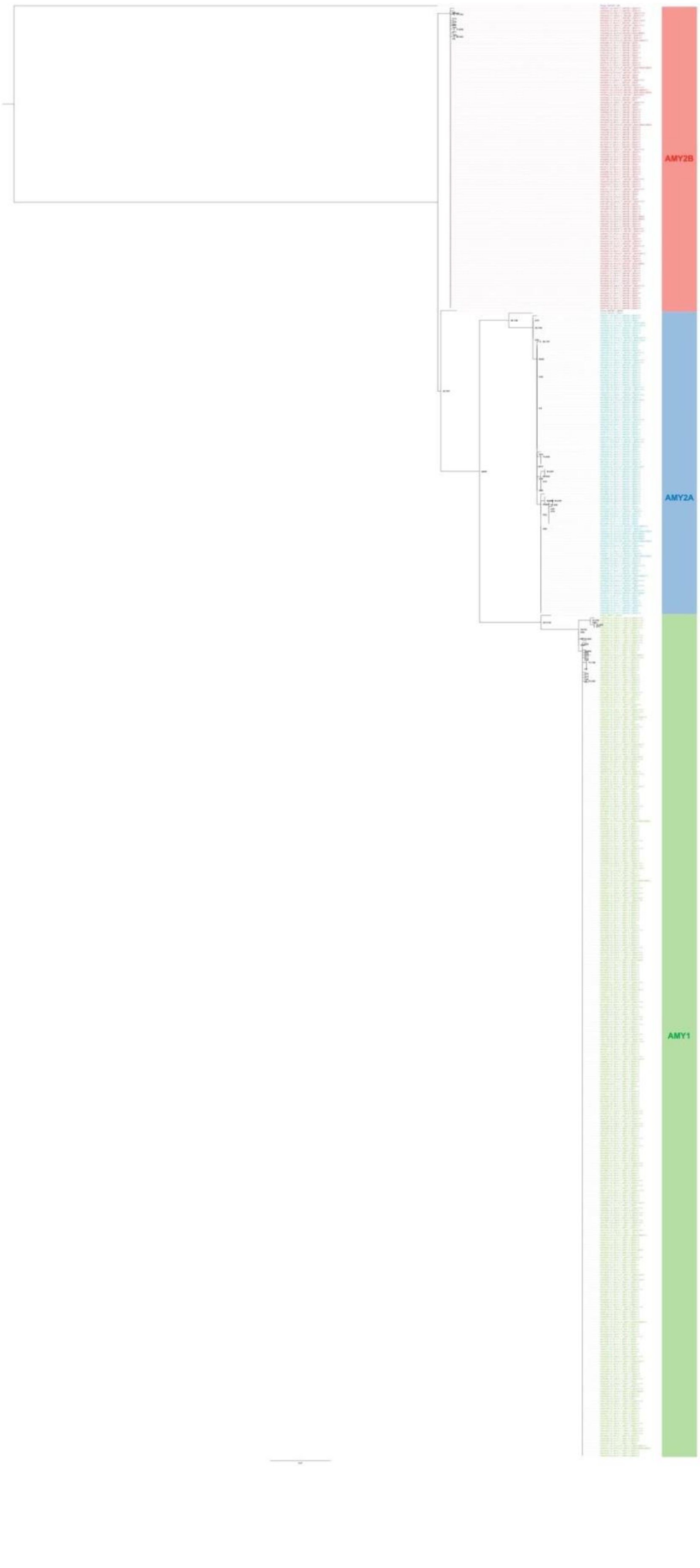

**Fig. S11. Maximum likelihood tree for coding sequences of all amylase genes.**
Amylase coding sequences from sheep (Sheep_AMY2B.1_2B), and chimpanzee (Chimpanzee_AMY2B.1_2B2A1, Chimpanzee_AMY2A.1_2B2A1, and Chimpanzee_AMY1.1_2B2A1) were used as outgroups for all amylase genes (*AMY2B*, *AMY2A*, and *AMY1*, respectively). The number on the node represents Shimodaira-Hasegawa approximate Likelihood Ratio Test (SH-aLRT) support (%) / ultrafast bootstrap support (%). The support for branches is calculated using the SH-aLRT. The SH-aLRT provides a percentage score that indicates the level of support for a particular branching (i.e., a split in the tree). A higher percentage indicates more robust support for the reliability of that branch, suggesting that the data strongly support the hypothesis that the sequences on either side of the branch are more closely related to each other than to sequences on different branches.

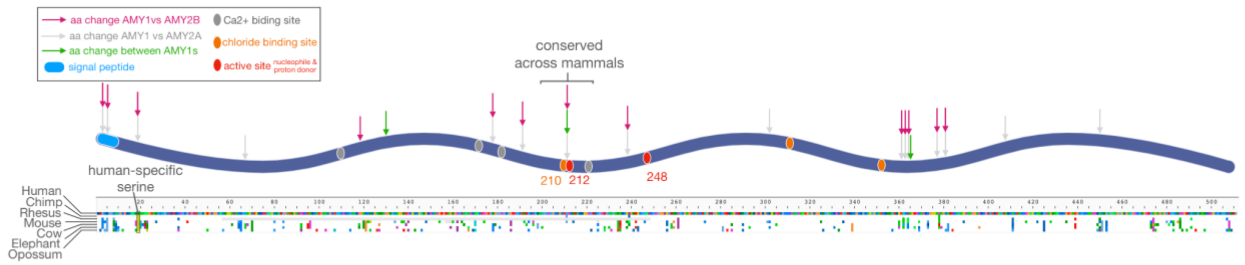


**Fig. S12. Positions of amino acid changes across 511 residues of amylase in the context of functional sites.**

Upper panel: The blue wavy line represents all aligned amylase peptide sequences. Colored ovals show approximate locations of previously defined functional domains in UniProtKB (Accession: P0DUB6), as noted in the boxed legend in the upper left. Numbers indicate specific functional domain sites. Arrows represent sites of amino acid changes when comparing *AMY1*, *AMY2A*, and *AMY2B* sequence alignments (colors indicated in the boxed legend). Lower panel: Alignment of AMY1-like predicted peptide sequences from different mammalian species (names labeled on the left). Colored squares indicate amino acid differences according to RasMol amino acid color scheme, while white space indicates sequence conservation.

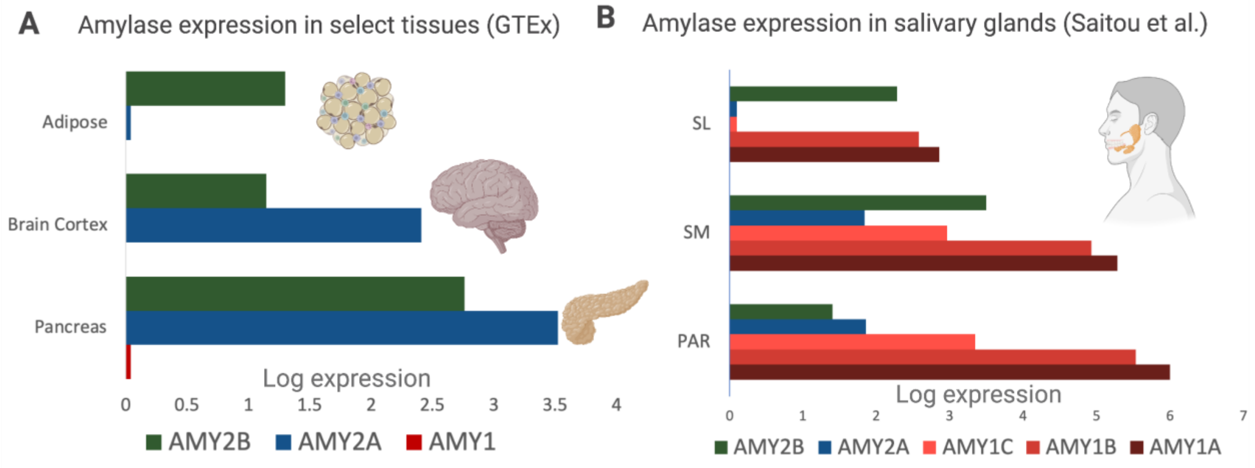

**Fig. S13. Expression of amylase genes across human tissues.**
(A) The panel shows the log expression for *AMY2B*, *AMY2A*, and *AMY1 genes* in adipose, brain cortex, and pancreas tissues. (B) The panel shows the expression of amylase genes in major salivary glands. This dataset is available through Saitou et al. (*20*). SL: Sublingual Gland, SM: Submandibular Gland , PAR: Parotid Gland.

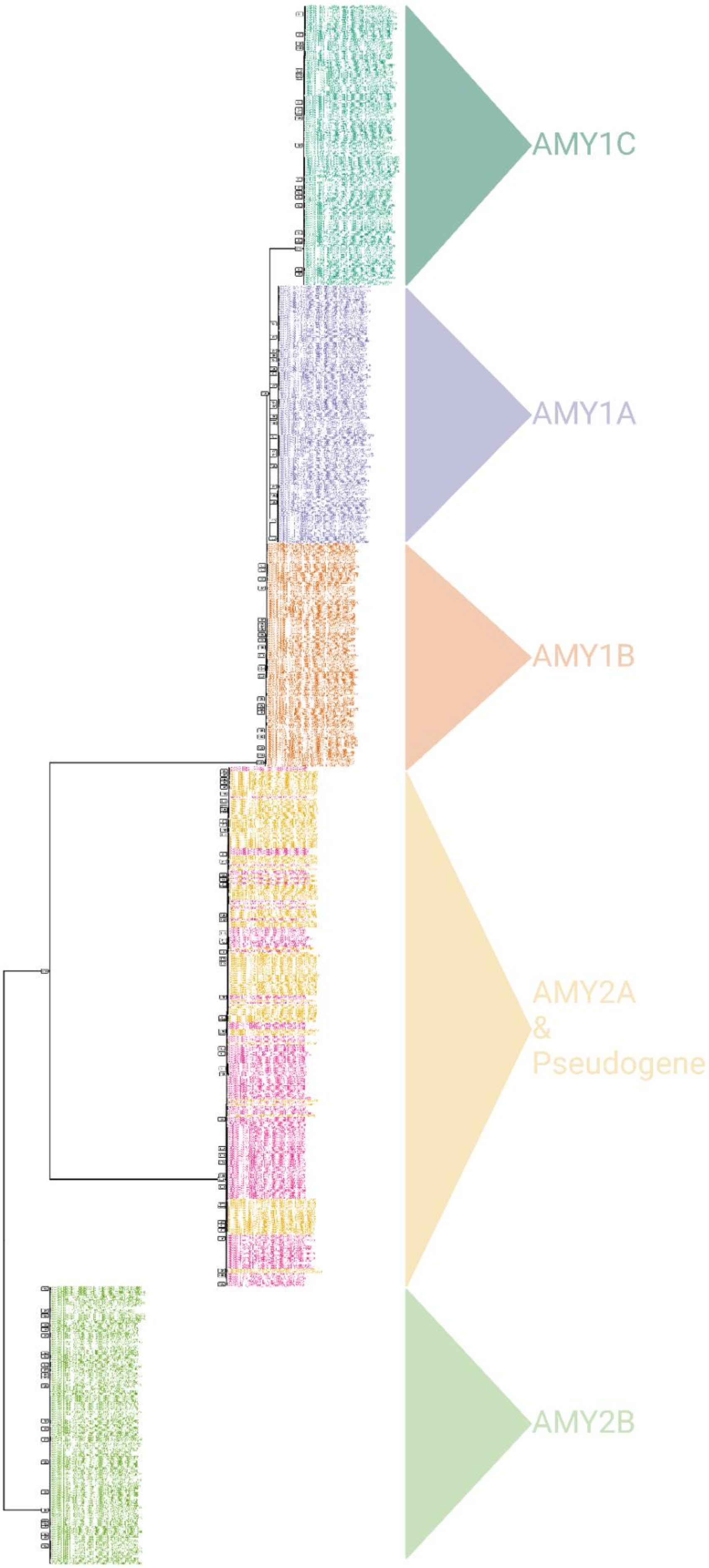




539

**Fig. S14. Maximum likelihood phylogenetic tree for the amylase segments that harbor the five distinct copies of the amylase genes and pseudogenes.**

The pseudogene originated from an ancestral incomplete gene duplication from the AMY2A segment. Pink color in the tree represents pseudogene copies. The support for branches is calculated using the Shimodaira-Hasegawa approximate Likelihood Ratio Test (SH-aLRT). The SH-aLRT provides a percentage score that indicates the level of support for a particular branching (i.e., a split in the tree). A higher percentage indicates more robust support for the reliability of that branch, suggesting that the data strongly support the hypothesis that the sequences on either side of the branch are more closely related to each other than to sequences on different branches.

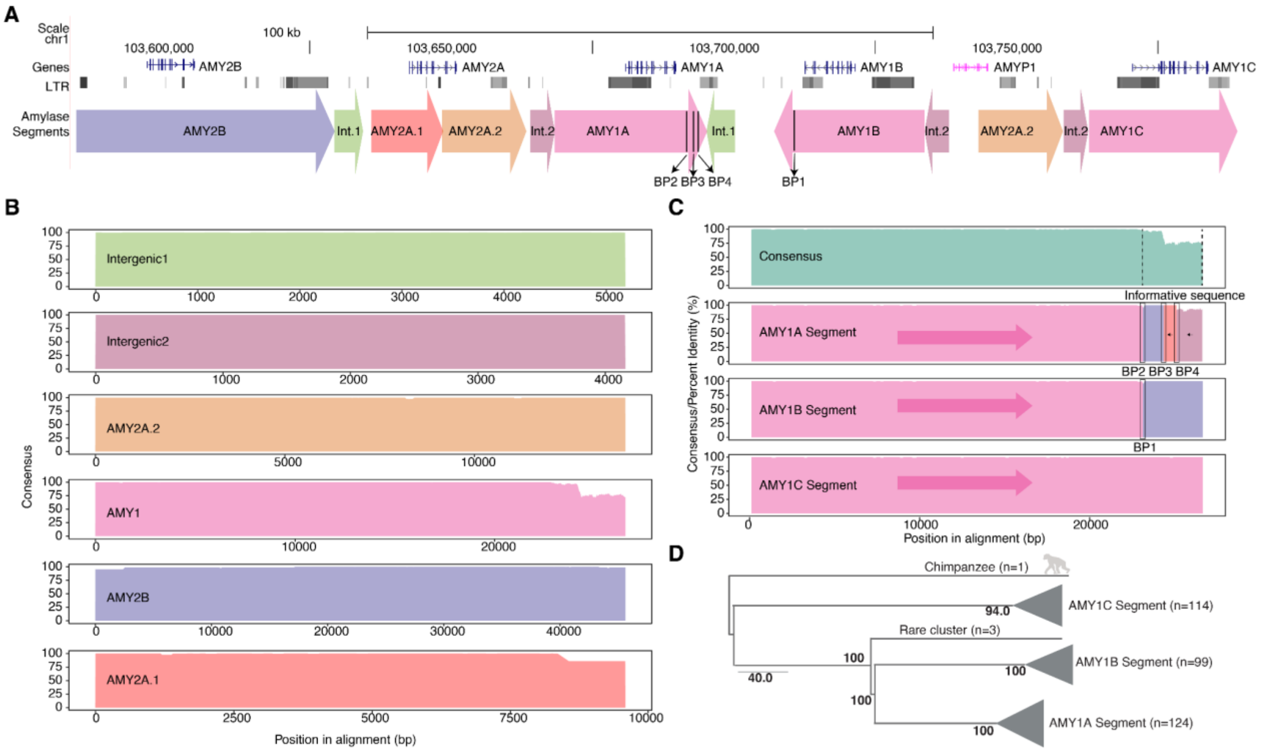


**Fig. S15. Characterization of the amylase segments.**

(A) The reference panel displays the position and orientation of each amylase segment, GENCODE V44 gene annotations, and long-terminal repeats (LTRs). The black lines in AMY1 segments represent the position of breakpoints depicted in panel C. (B) Consensus of multiple sequence alignment for Intergenic1, Intergenic2, AMY2A.2, AMY1, AMY2B, and AMY2A.1 amylase segments. The consensus is calculated by averaging frequencies of major alleles (including a gap) across a 200-bp window with 10-bp step. The x-axis represents position in alignment (bp). (C) Consensus of AMY1 segment alignment (top) and sequence composition of each AMY1 segment cluster (bottom). The y-axis represents the average % identity of a 200-bp window from the consensus sequence of each AMY1 segment cluster versus the best hit from the BLAT search against the pseudo reference sequence (see "Detecting sequence origin and breakpoints in AMY1 segments" in Materials and Methods). Dashed lines in the consensus plot (top) represent an interval (from 22,850 to 26,730 in the AMY1 segment alignment, Data File-S2) harboring an informative sequence. The inverted black arrows at the 3' end of the AMY1A segment indicate that the best hits from AMY2A.1 and Intergenic2 segments are oriented in the opposite direction to the AMY1A segment. The dashed boxes in the AMY1A and AMY1B segments indicate the regions where the possible breakpoints for each recombination event are located. (D) Maximum likelihood tree consisting of three AMY1 segment clusters (AMY1C, AMY1B, and AMY1A), rooted by the chimpanzee (mPanTro3) AMY1 segment. The tips within each cluster are collapsed for visualization. The number in each node represents bootstrap support (%). The branch length may not reflect the actual distance between tips as the 3' end of AMY1 segments are not well aligned.

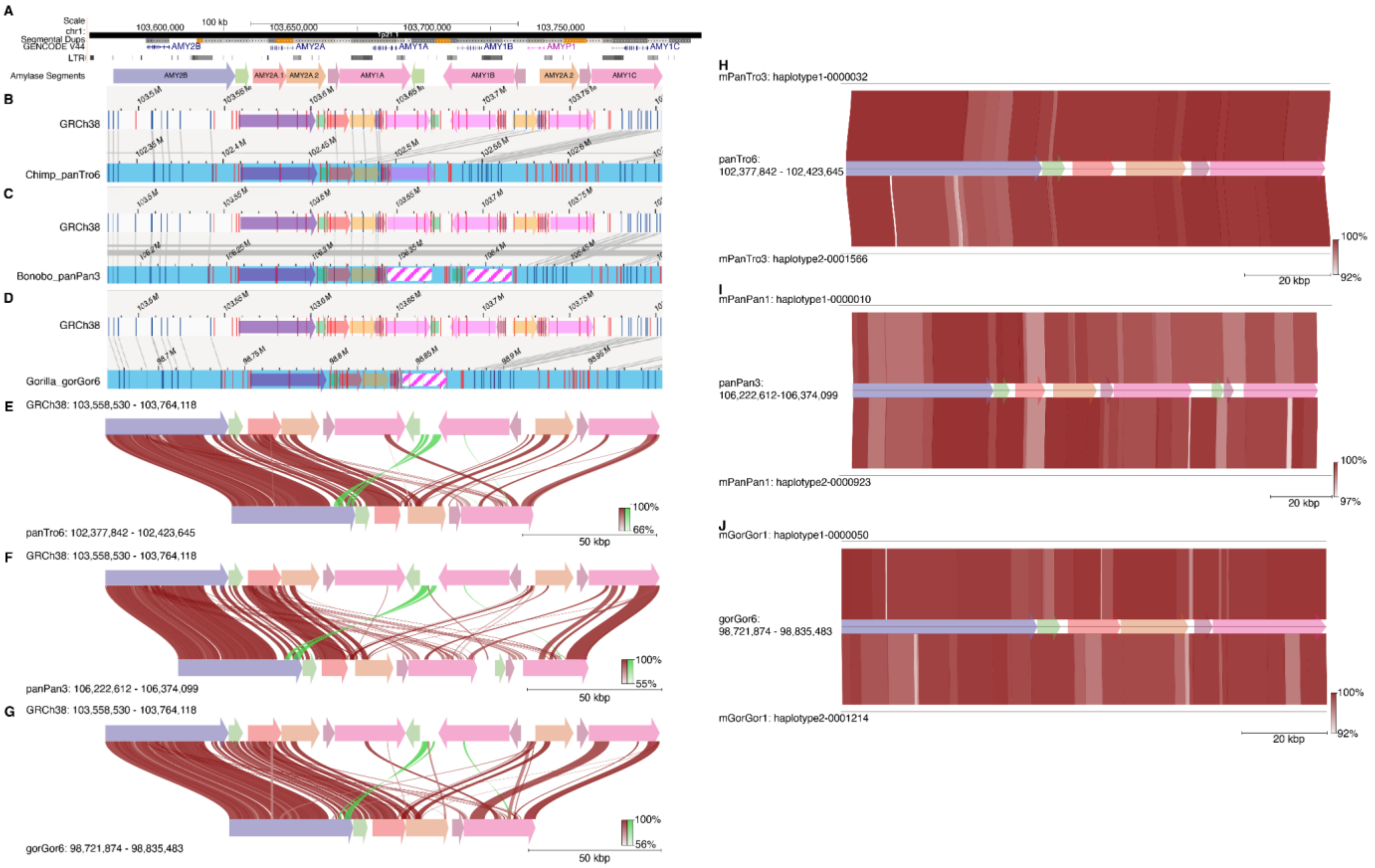


**Fig. S16. Structure of amylase locus in nonhuman primates: chimpanzee, bonobo, and gorilla.**

(A) GENCODE V44 gene annotations, long-terminal repeats (LTRs), and segmental duplications are shown as tracks. The lower panel displays amylase segments (colored arrows). (B) *In silico* map alignment between GRCh38 (white background with blue vertical lines) and chimpanzee (blue background with blue vertical lines) reference genomes. (C) I*n silico* map alignments between GRCh38 and bonobo reference genomes. (D) I*n silico* map alignments between GRCh38 and gorilla reference genomes. Striped AMY1 segments in bonobo and gorilla represent segments in which the orientation is unclear from the *in silico* maps. Lines between GRCh38 and nonhuman primate *in silico* maps show aligned labels. Pairwise sequence alignments of the amylase locus from GRCh38 and (E) chimpanzee - panTro6, (F) bonobo - panPan3, and (G) gorilla - gorGor6 are visualized as synteny plots using pygenomeviz (*23*). Pairwise sequence alignments of the amylase locus from (E) panTro6 to mPanTro3, (F) panPan3 to mPanPan1, and (G) gorGor6 to mGorGor1 are visualized as synteny plots using pygenomeviz (*23*).

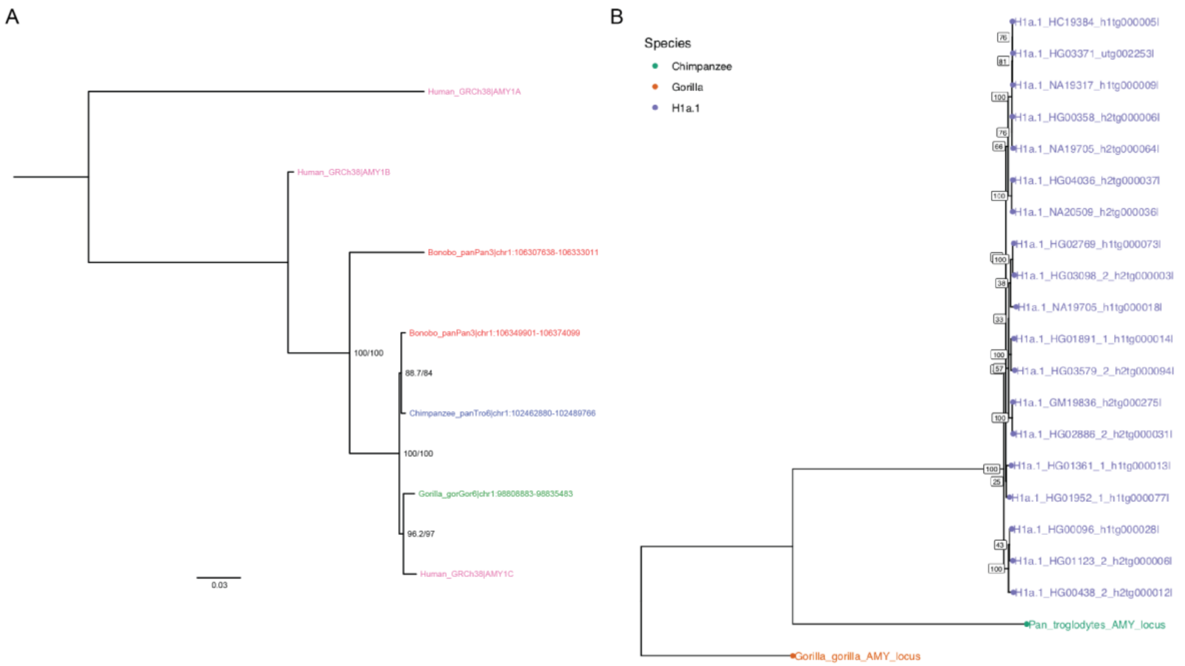


**Fig. S17. Phylogenetic tree showing the relationship of AMY1 segments found in great apes.** (A) Maximum likelihood tree for the 3' end interval of the AMY1 segment (from 22,850-bp to 26,730-bp in the AMY1 segment alignment, Data File-S2) in nonhuman primates (Human; GRCh38, Chimpanzee; panTro6, Bonobo; panPan3, and Gorilla; gorGor6). The tree is rooted by the midpoint. The number on the node represents Shimodaira-Hasegawa approximate Likelihood Ratio Test (SH-aLRT) support (%) / ultrafast bootstrap support (%). The SH-aLRT provides a percentage score that indicates the level of support for a particular branching (i.e., a split in the tree). A higher percentage indicates more robust support for the reliability of that branch, suggesting that the data strongly support the hypothesis that the sequences on either side of the branch are more closely related to each other than to sequences on different branches. (B) Phylogenetic tree for the 19 present-day human H1$^{a}$.1 haplotypes. The syntenic amylase loci of chimpanzee (Pan troglodytes; NHGRI_mPanTro3-v2.0_pri) and gorilla (Gorilla gorilla gorilla, NHGRI_mGorGor1-v2.0_pri) were used to construct the phylogenetic tree.

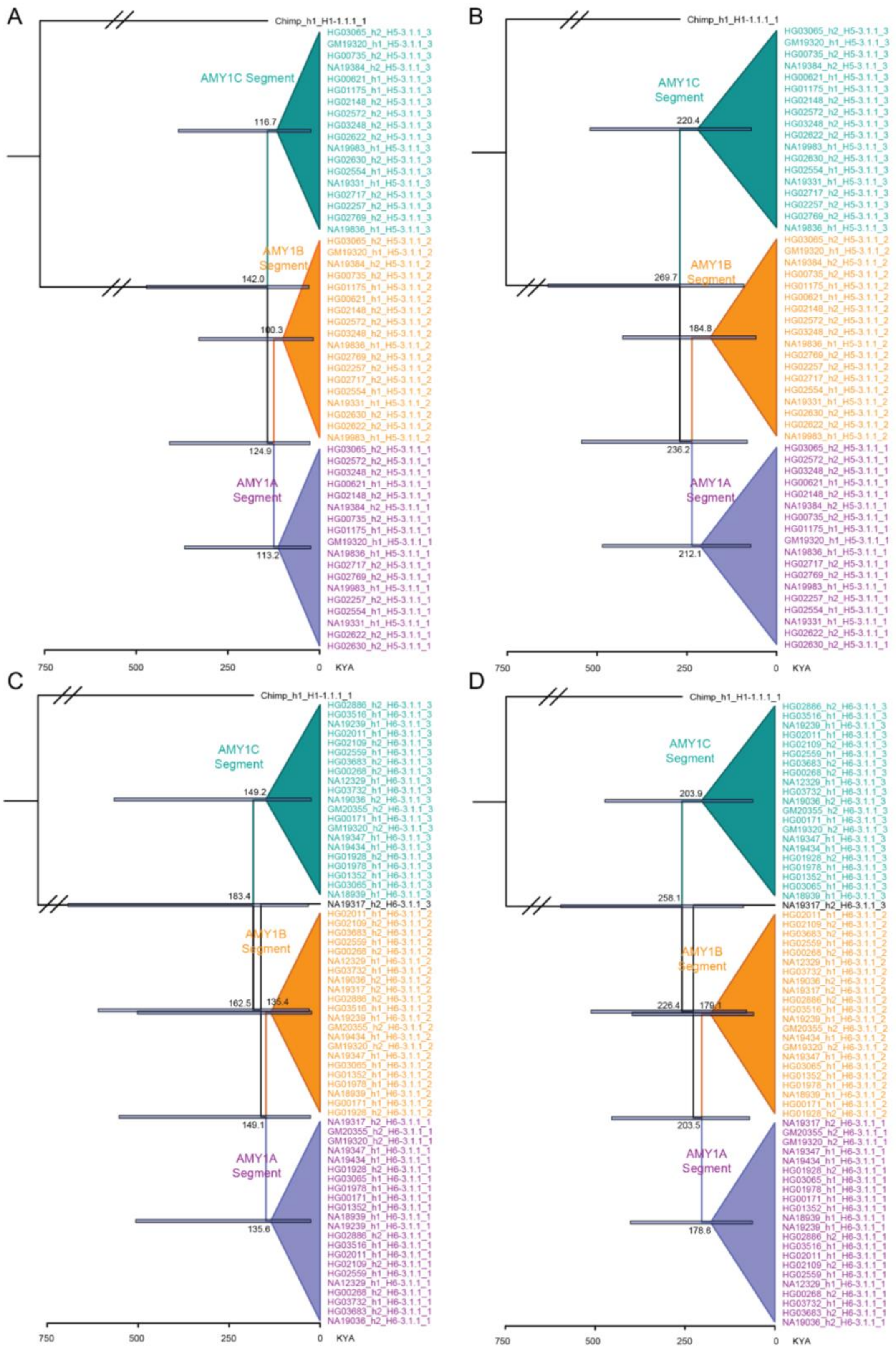


**Fig. S18. Dated Bayesian phylogenetic trees for H3$^r$.1 and H3$^r$.2 AMY1 segments.**
(A) H3$^r$.1 AMY1 segment tree using the "Coalescent Bayesian Skyline" prior and "Optimized Relaxed Clock" model. (B) H3$^r$.1 AMY1 segment tree based on the "Coalescent Constant

Population” prior and “Optimized Relaxed Clock” model. (C) H3$^{r}$.2 AMY1 segment tree using the “Coalescent Bayesian Skyline” prior and “Optimized Relaxed Clock” model. (D) H3$^{r}$.2 AMY1 segment tree based on the “Coalescent Constant Population” prior and “Optimized Relaxed Clock” model. The labels on nodes indicate point estimates of divergence time. The bars on nodes indicate a 95% Highest Posterior Density (HPD) interval. Each tree is rooted by the chimpanzee AMY1 segment (Chimpanzee_h1_H1a.1_1) and child nodes are collapsed by their AMY1 segment categorization (AMY1C, AMY1B, and AMY1A). KYA: thousand years ago.

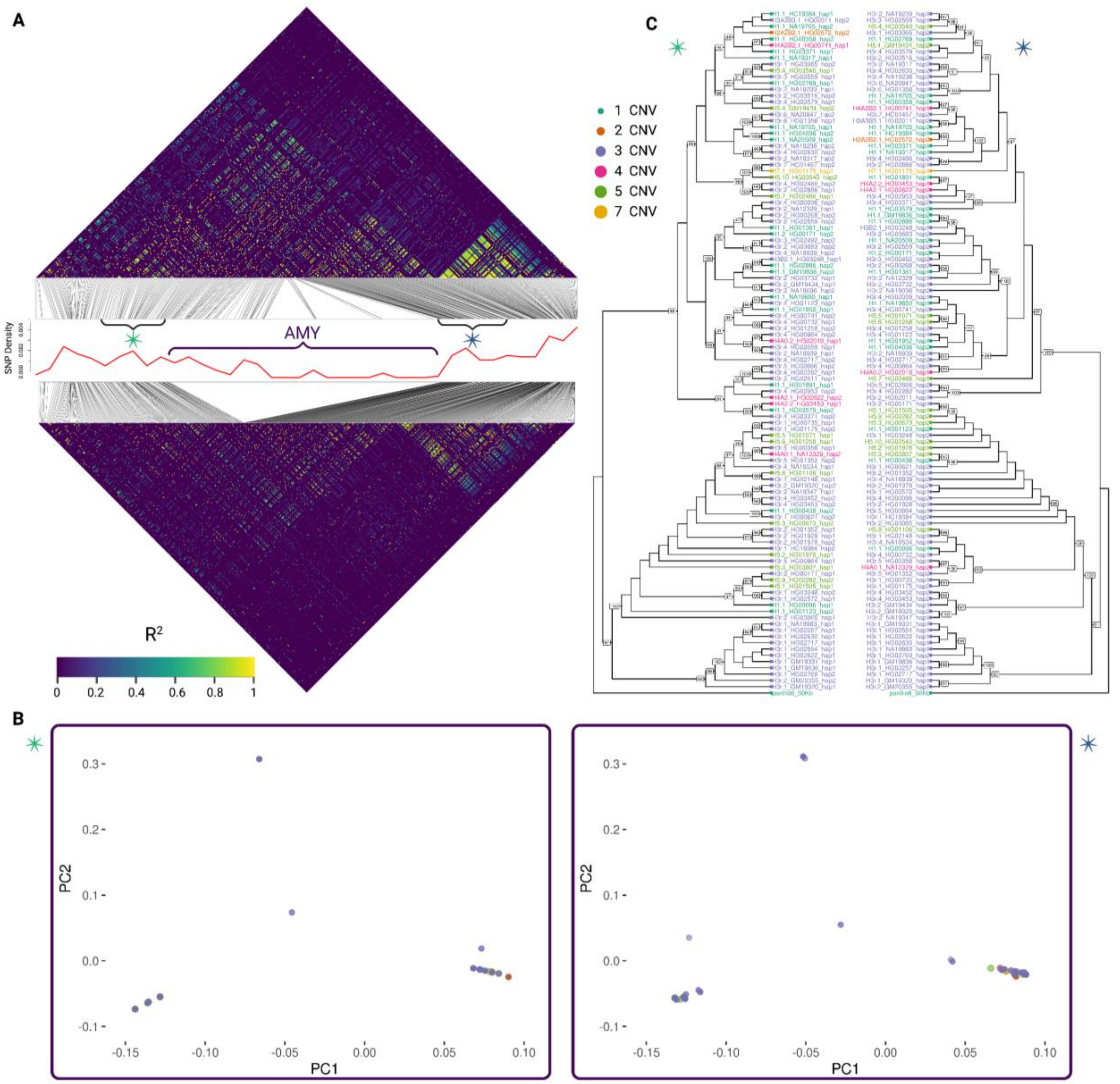


**Fig. S19. Linkage disequilibrium and phylogenetic clustering of the amylase structural haplotypes.**

(A) Displays linkage disequilibrium (LD) $R^2$ estimates for the amylase locus and its flanking sequences, incorporating 100-kbp upstream and downstream regions of the amylase locus. The upper LD heatmap was constructed using the 1000 Genomes Project Phase 3 variant calls. The lower LD heatmap was constructed using haplotype-resolved diploid assemblies from this study. (B) Principal component analysis (PCA) of assemblies from our study, constructed from the 50-kbp upstream (left) and downstream regions (right). (C) The two phylogenetic trees (depicted as cladograms for simplicity) representing the genomic regions 50-kbp upstream (left) and downstream (right) of the amylase locus illustrate the evolutionary relationships within each of these two regions. The numbers on the nodes represent the bootstrap support.

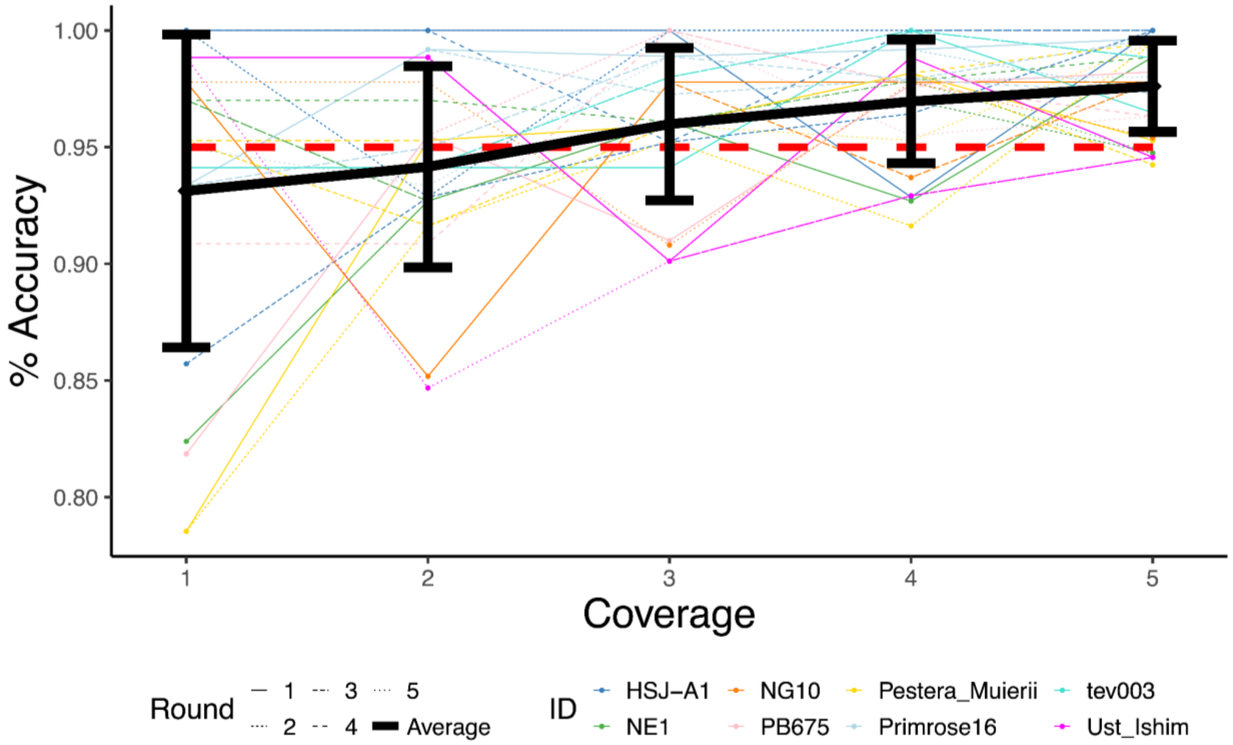


**Fig. S20. Downsampling of eight ancient genomes.**
The effect of coverage on the percent accuracy of the *AMY1* copy number estimation using the full genome data (here termed "% Accuracy"). The genomes were chosen to account for batch effects: four different laboratories, a mix of hunter-gatherers and agriculturalists, an estimated 6-17 copies of the *AMY1* gene, and dates ranging from 34,000 years ago to within the last 1,000 years. Each genome was randomly downsampled to 1-5X coverage, five separate times. Color denotes genome, and line shape denotes round of downsampling. The black line represents the average accuracy from all downsampling results for all genomes at each coverage level. The error bars represent the standard deviation of % accuracy results for each coverage.

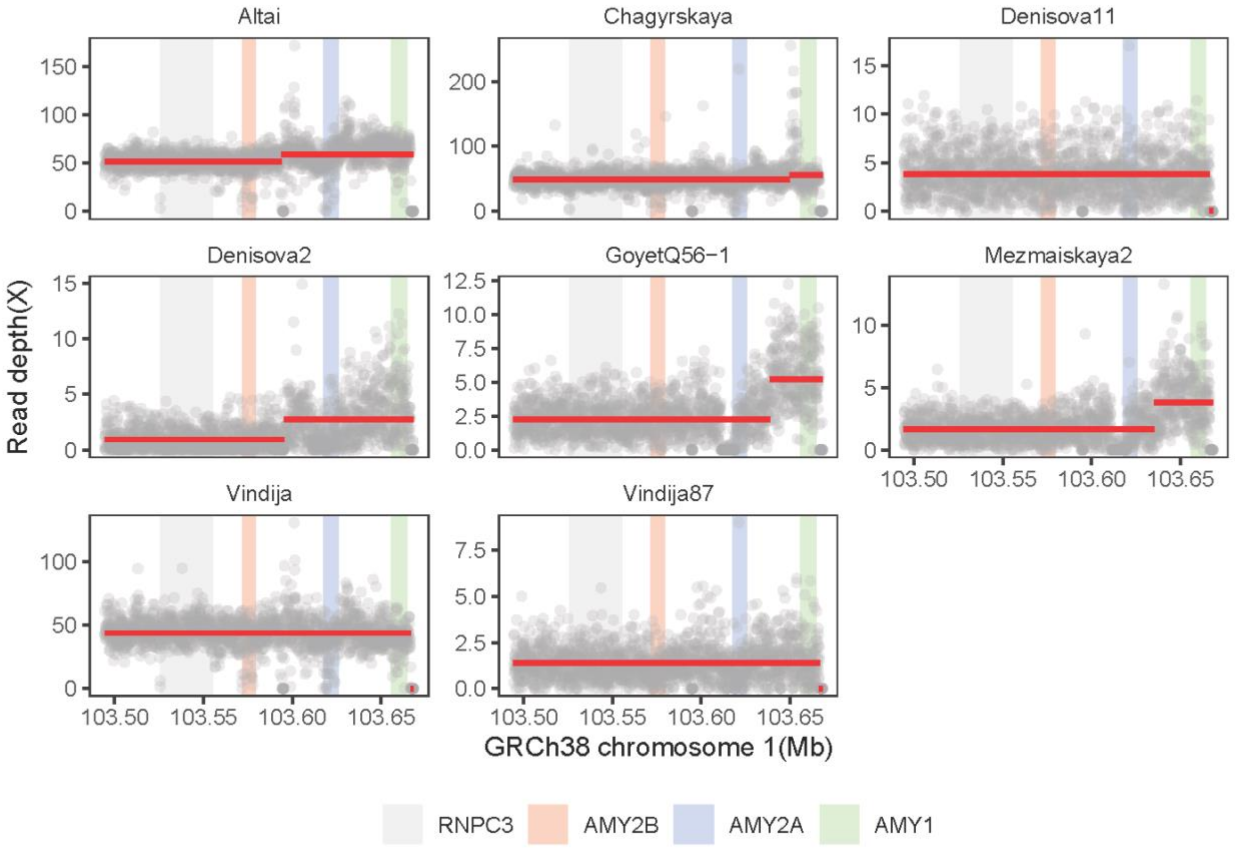


**Fig. S21. Read-depth plots for the amylase locus of archaic hominin genomes.**
The GC-bias corrected read-depth for the amylase locus spanning the *RNPC3* (gray), *AMY2B* (red), *AMY2A* (blue), and *AMY1* (green) genes. Each individual data point represents an average read-depth over a non-overlapping 100-bp window. The red line indicates the average read-depth of each segmented region determined by the changepoint analysis method "AMOC". Only uniquely mapped reads were used for this visualization.

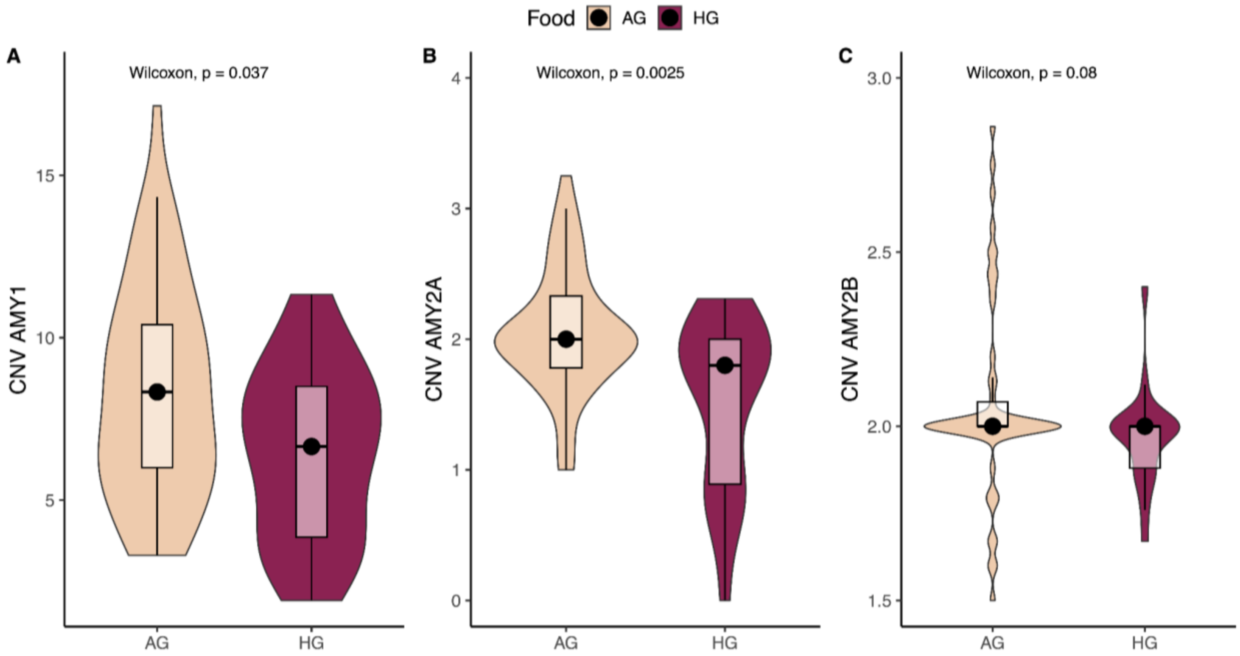


**Fig. S22. *AMY1* copy number in ancient human genomes by diet.**
A violin plot comparison of (A) *AMY1*, (B) *AMY2A*, and (C) *AMY2B* copy numbers between ancient human hunter-gatherers (HG) and agriculturalists (AG) (n = 73). Wilcoxon p-values testing whether there is a significant difference in gene copy number between the two dietary habits are given. The black circle in each boxplot represents the median of each dataset.

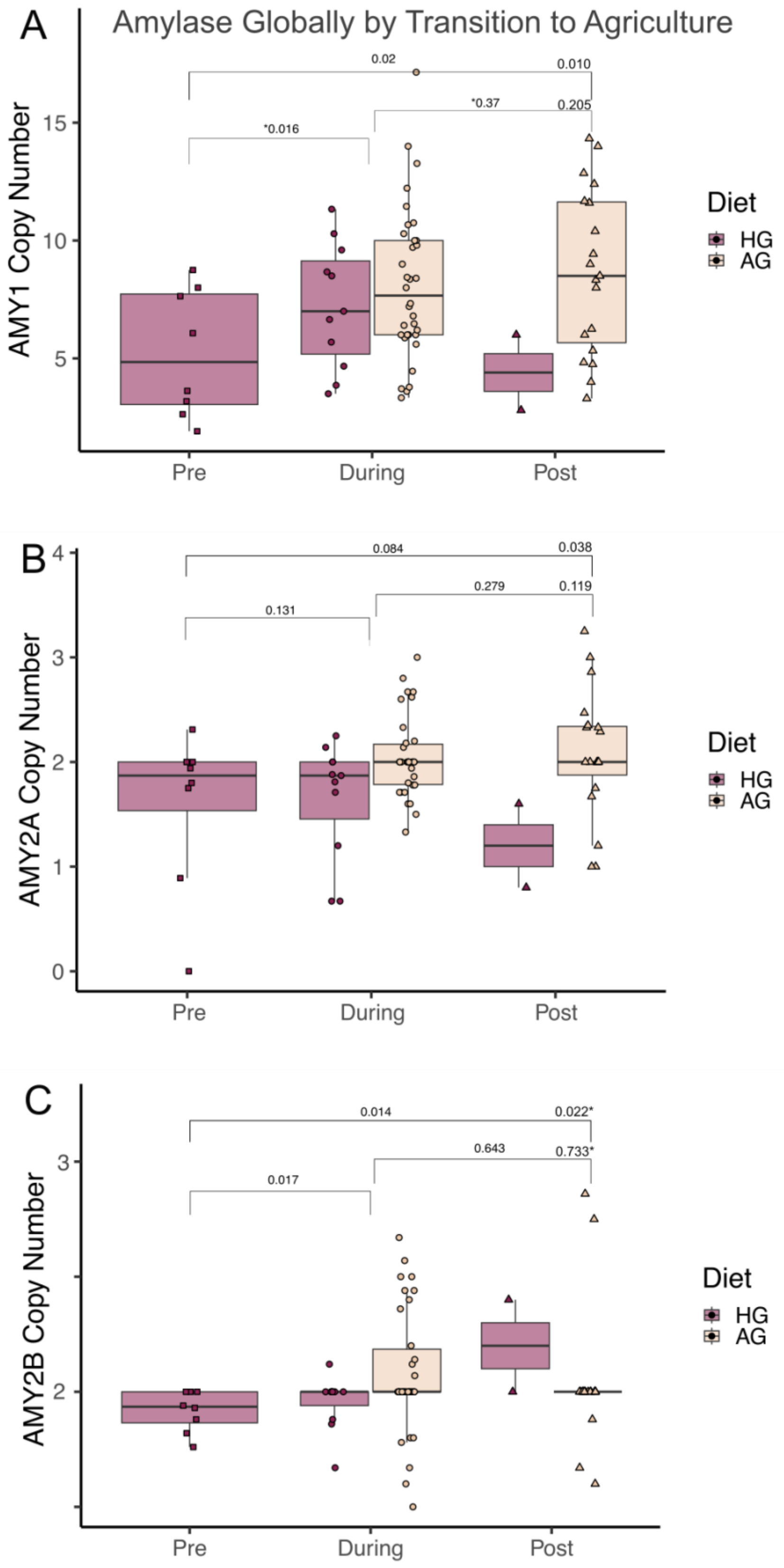
A
Amylase Globally by Transition to Agriculture
AMY1 Copy Number
5
10
15
0.02
0.010
*0.016
*0.37
0.205
Pre
During
Post
Diet
HG
AG
B
AMY2A Copy Number
0
1
2
3
4
0.084
0.038
0.131
0.279
0.119
Pre
During
Post
Diet
HG
AG
C
AMY2B Copy Number
2
3
0.014
0.022*
0.017
0.643
0.733*
Pre
During
Post
Diet
HG
AG

**Fig. S23. *AMY1* copy number calculated globally in ancient human genomes using GeneToCN as compared to time.**

"During" represents 9,000-3,000 years before present (BP), as this encompasses the majority of the transition to agriculture worldwide plus the additional time it would have taken for the copy number to adjust at the population level. Modern samples younger than ~500 years old are not included. The copy numbers between time periods are compared based on a one-sided Wilcoxon test with the alternative hypothesis that older time periods have a lower copy number. Hunter-gatherers (HG) and agriculturalists (AG) were combined in the statistical tests and separated only for visualization. Two hunter-gatherer individuals in the "Post" category are ~1,000 year-old individuals from maritime communities (Kawéskar and Yámana territory) in Chilean Patagonia. The believed descendants of these populations practiced hunter-gathering until the arrival of Europeans in the 16th century and therefore, never transitioned to an agriculturally dependent lifestyle. The stars indicate p-values from additional Wilcoxon tests calculated between "Post" and "During" and between "Post" and "Pre" without these two hunter-gatherers.

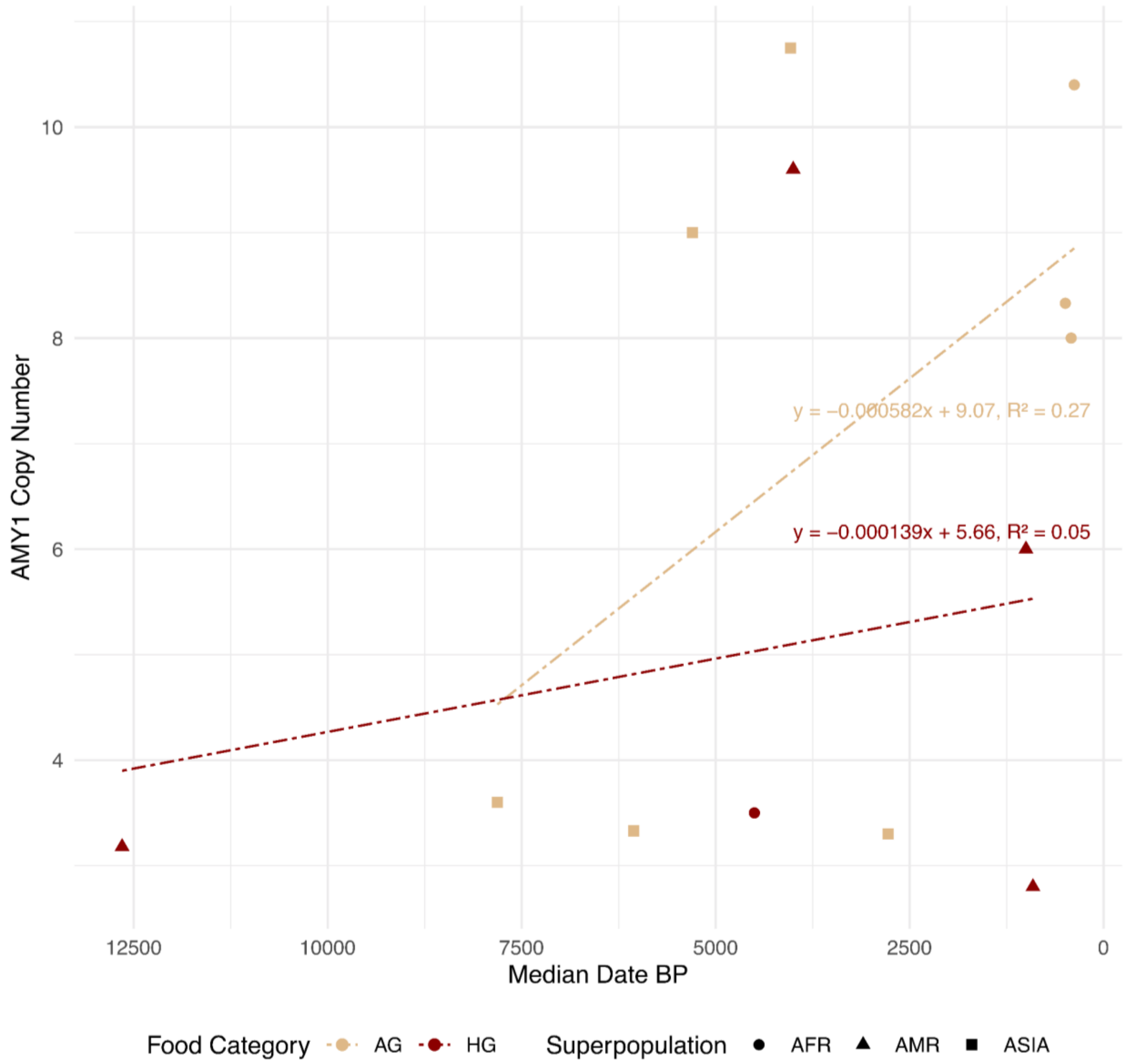


**Fig. S24. The temporal distribution of *AMY1* gene copy number variation in ancient human non-European populations, analyzed using the GeneToCN tool.**
Data points are colored by subsistence strategy: hunter-gatherers (HG) in red and agriculturalists (AG) in beige. The scatter plot shows individual samples plotted against their median dating in years before present (BP). Linear regression trendlines are included for both hunter-gatherers and agriculturalists, with the respective regression equations and $R^2$ values displayed to indicate the strength and direction of the trends over time.

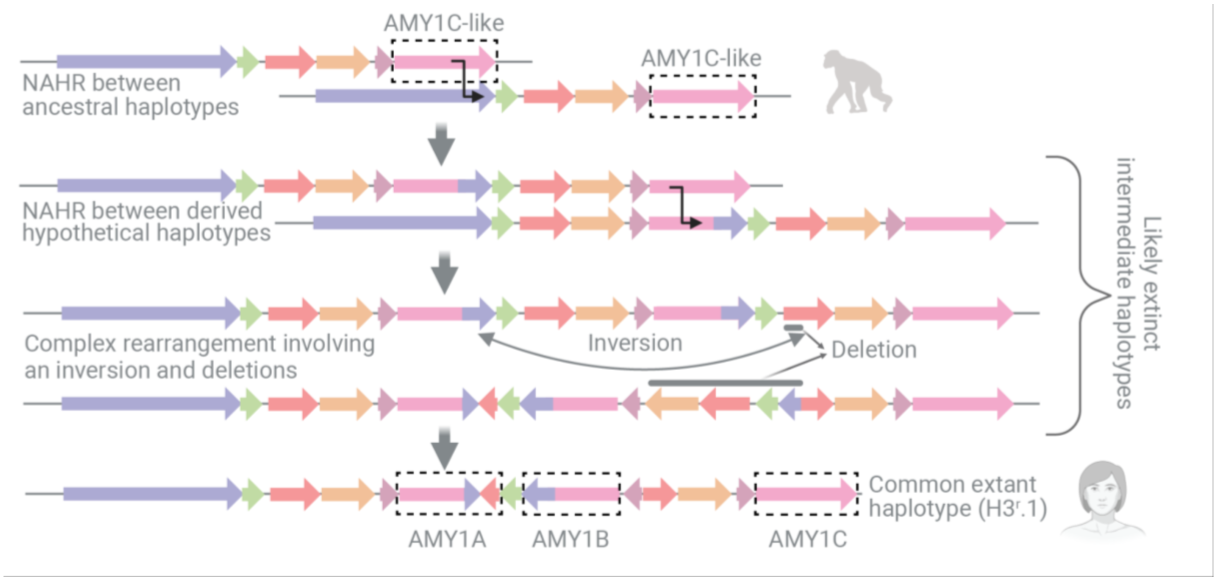


**Fig. S25. Proposed mechanism for the *AMY1* gene.**

We hypothesize that the initial duplication of the *AMY1* gene was facilitated by non-allelic homologous recombination (NAHR) between the ancestral AMY1 segment and the AMY2B segment. This would result in a haplotype consisting of one AMY2B segment but two AMY2A and AMY1 segments. The newly duplicated AMY1 segment appears chimeric, having acquired a part of the AMY2B segment (indicated in purple). This intermediate haplotype would then undergo a second NAHR event between the proximal and distal AMY1 segments, leading to the duplication of the *AMY1* gene with a haplotype carrying three AMY1 segments. A subsequent inversion-deletion event, as illustrated, would then produce a haplotype with three copies of the *AMY1* gene similar to the extant H3$^r$.1 haplotype.

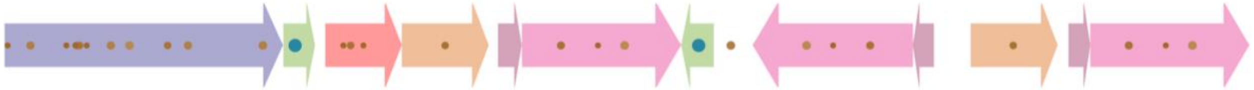

**Fig. S26. PRDM9 binding sites including degenerate sites, for the consensus $H3^r$.1 haplotype sequence were identified using FIMO.**

We used the non-redundant DNA database (NRDB) as background frequencies across the amylase locus (using the $H3^r$.1 consensus sequence as a template). Larger blue circles indicate perfect matches, and smaller brown circles indicate near-perfect matches (one mismatch permitted). For all matches, we require the nominal p-value to be equal to or less than 0.0001. Individual p-values and match scores can be found in **supplementary table S13**.

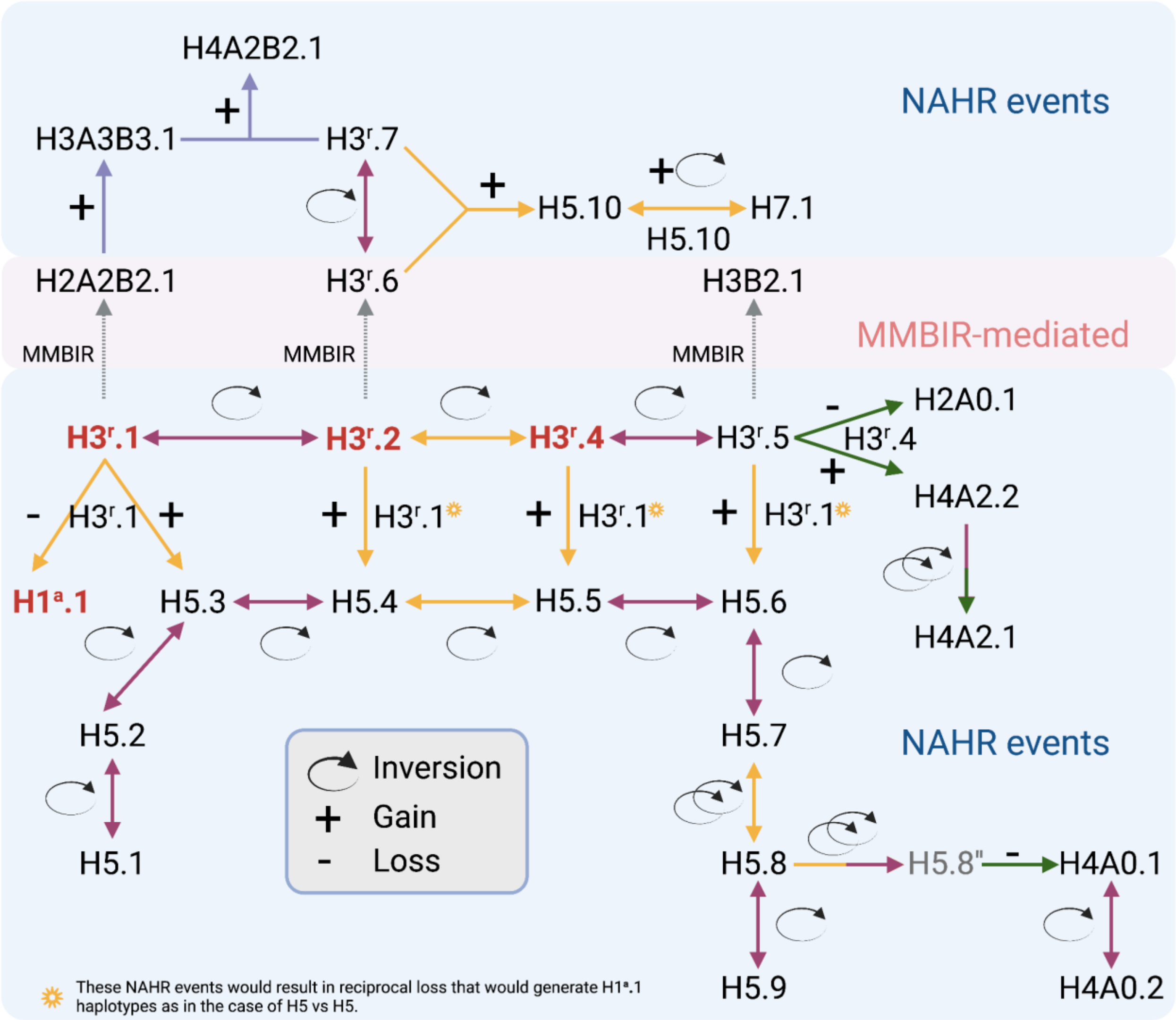


**Fig. S27. Model for mutational events network involving amylase structural haplotypes.** This mutational events network exhibits common haplotypes (H1$^{a}$.1, H3$^{r}$.1, H3$^{r}$.2, and H3$^{r}$.4 - red colored text), microhomology-mediated break-induced replication events (H2A2B2.1, H3$^{r}$.6, H3B2.1), recurrent inversions among five-copy haplotypes (H5.1, H5.2, H5.3, H5.4, H5.5, H5.6, H5.7, H5.8 and H5.9) and other haplotypes (H2A0.1, H3$^{r}$.5, H3$^{r}$.6, H3$^{r}$.7, H4A0.1, H4A0.2, H4A2.1, H4A2.2, H4A2B2.1, H5.10 and H7.1). The colors of the arrows indicate the differently-colored sequences that are involved in the non-allelic homologous recombination events, either as recombination substrates for copy number gains and losses or as inverted repeats that would generate inversions. Specifically, maroon arrows represent inversions mediated by the intergenic2 inverted segments, orange arrows represent either inversions mediated by the inverted AMY2A.2 segments or crossover across the directly oriented AMY2A.2 segments, and subsequent copy-number gains or losses. Further details about the comparison of haplotypes and mechanistic inferences can be found in the **Supplementary Materials and Methods**.

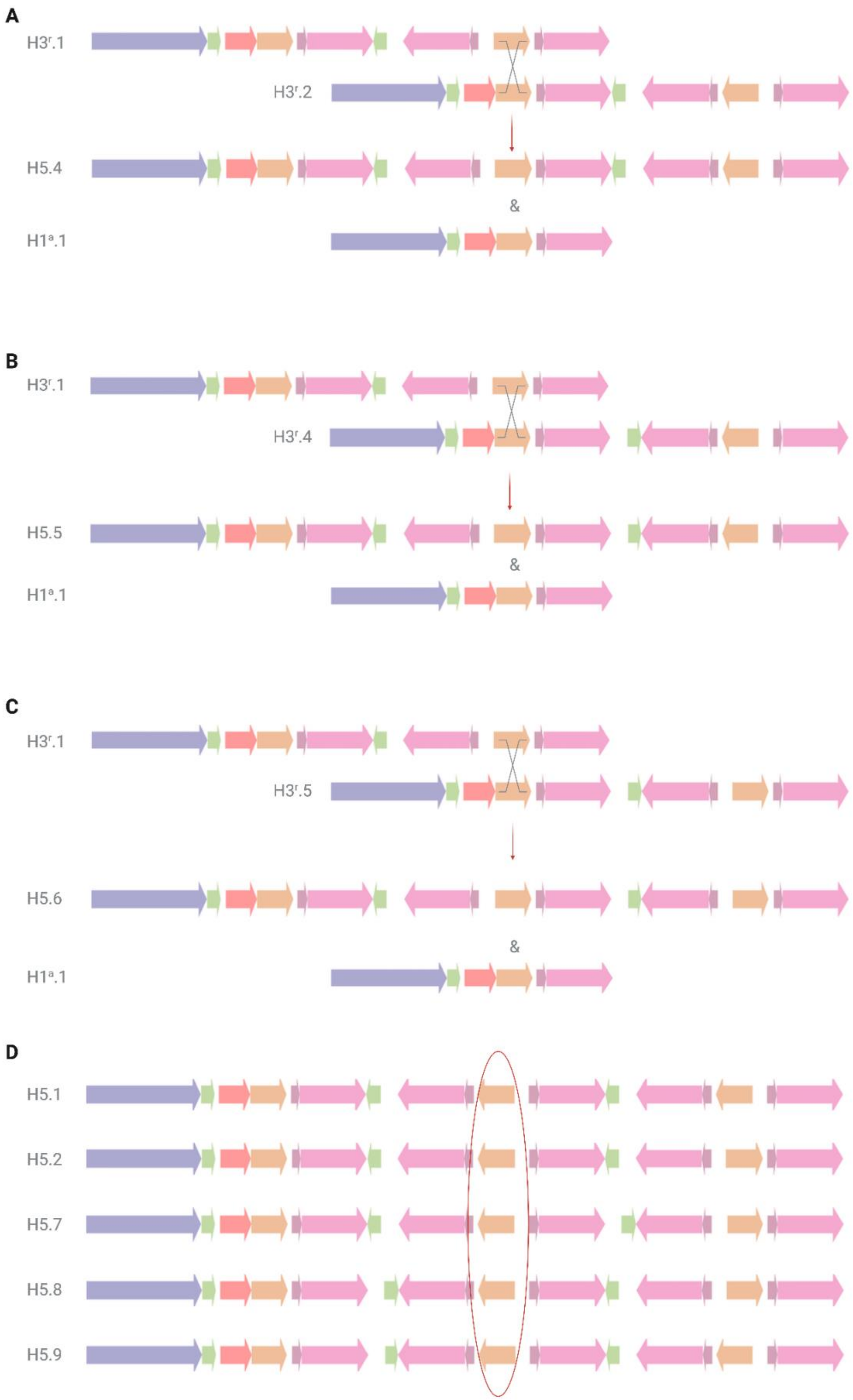

A
$H3^{r}.1$
$H3^{r}.2$
H5.4
&
$H1^{a}.1$
B
$H3^{r}.1$
$H3^{r}.4$
H5.5
&
$H1^{a}.1$
C
$H3^{r}.1$
$H3^{r}.5$
H5.6
&
$H1^{a}.1$
D
H5.1
H5.2
H5.7
H5.8
H5.9

**Fig. S28. Non-allelic homologous recombination events among common amylase haplotypes.** Non-allelic homologous recombination (NAHR) among common haplotypes harboring three *AMY1* gene copies ($H3^r$.1, $H3^r$.2, $H3^r$.4, $H3^r$.5) with breakpoints in the AMY2A.2 segment (orange) were represented. Panels (A) to (C) represent the copy number gain and loss and resulting structural haplotypes facilitated by the crossover between: (A) $H3^r$.1 and $H3^r$.2, (B) $H3^r$.1 and $H3^r$.4, (C) $H3^r$.1 and $H3^r$.5. (D) This panel indicates the relative positions and orientations of the AMY2A.2 segments. Similar crossover events involving AMY2A.2 segments may not be feasible in all $H3^r$ haplotypes.

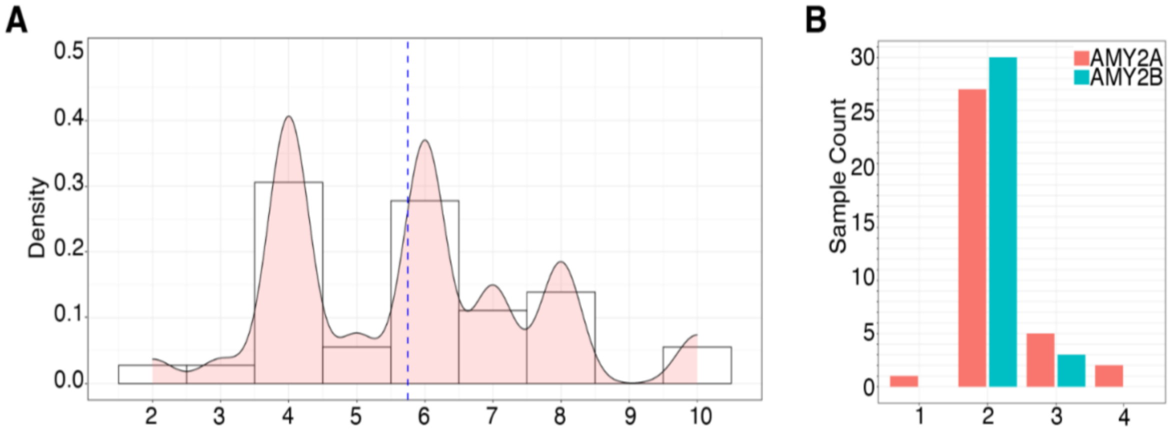


**Fig. S29. Copy number distribution of *AMY1* and *AMY2* genes.**
Copy numbers of *AMY1* and *AMY2* genes were detected using samples (n = 36) with haplotype structures on both chromosomes that were supported by optical genome mapping and long-read sequencing based assemblies. (A) Density plot of *AMY1* copy numbers. Copy numbers are shown on the x-axis and density is shown on the y-axis. (B) Bar plots of *AMY2A* and *AMY2B* copy numbers. Copy numbers are shown on the x-axis and sample count is represented on the y-axis.

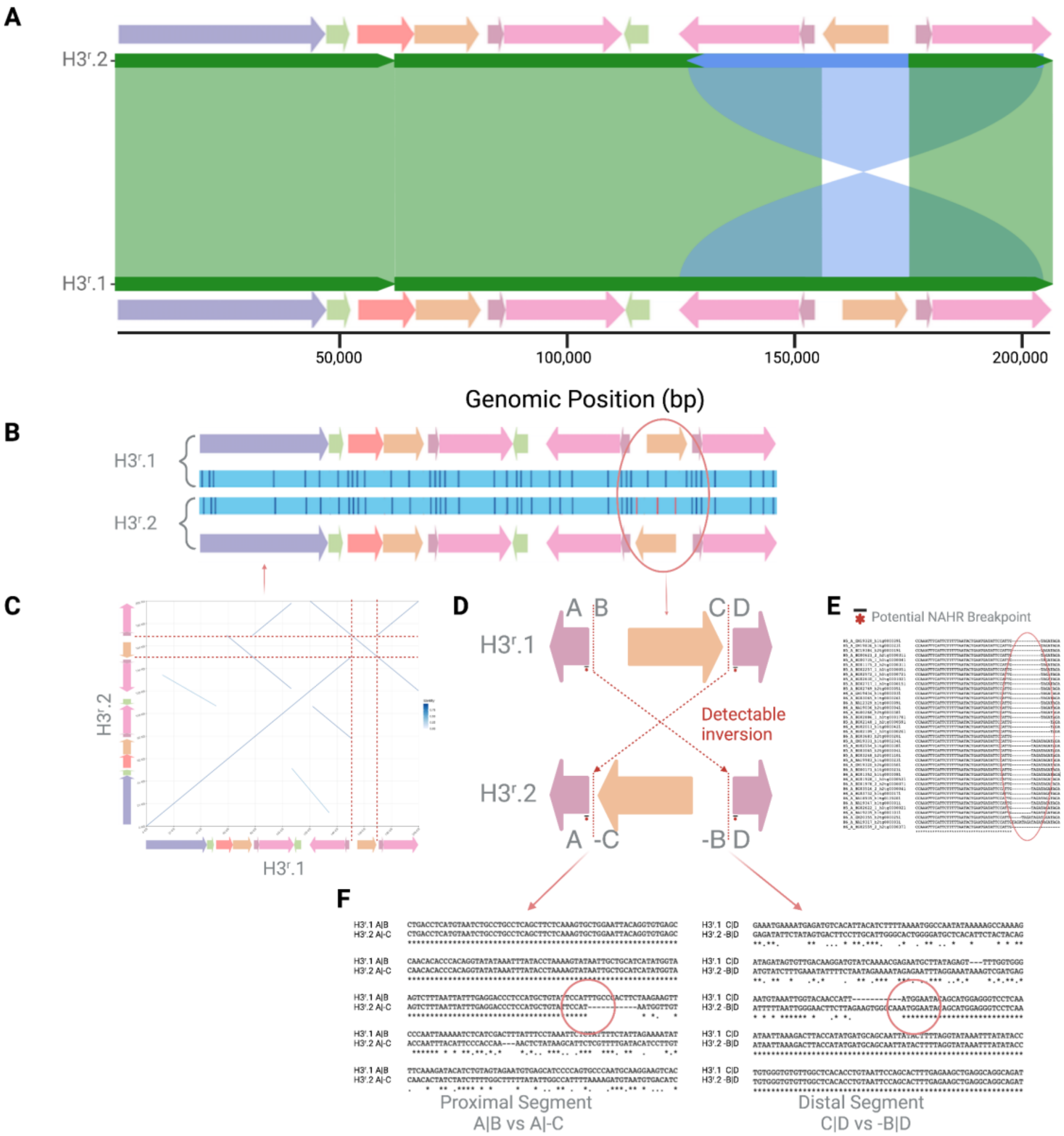


**Fig. S30. Schematic representation of the breakpoint detection analysis.**
(A) MIROPEAT plot for the global alignment comparison of two common amylase haplotypes, H3$^{r}$.1 and H3$^{r}$.2. The plot suggests that the two haplotypes harbor the same number of segments yet differ by an inversion of the terminal AMY2A.2 segment. (B) Optical genome mapping data confirms the presence of the inverted AMY2A.2 segment. (C) Dotplot visualization of the global alignment of the two haplotypes provides a fine-scale look at the breakpoints. (D) A schematic diagram for the inversion event between H3$^{r}$.1 and H3$^{r}$.2 haplotypes, with the proximal and distal breakpoints, provides the nucleotide resolution of the inversion breakpoints. The detectable inversion breakpoints are depicted with dotted lines. The proximal breakpoint in H3$^{r}$.1 is partitioned into fragments A and B, and the distal breakpoints into fragments C and D. The inversion brings fragment A into physical proximity to -C and fragment D into close linkage with -B in the H3$^{r}$.2 haplotype. (E) Local alignment of the A fragment across the reconstructed H3$^{r}$.1 and H3$^{r}$.2 haplotypes. (F) (left) Local alignment of the proximal breakpoint A|B in H3$^{r}$.1 and A|-

C in $H3^r$.2. The red circle highlights the detectable inversion breakpoints. (right) Local alignment of the distal breakpoint C|D in $H3^r$.1 and -B|D in $H3^r$.2. The red circle highlights the detectable inversion breakpoints.

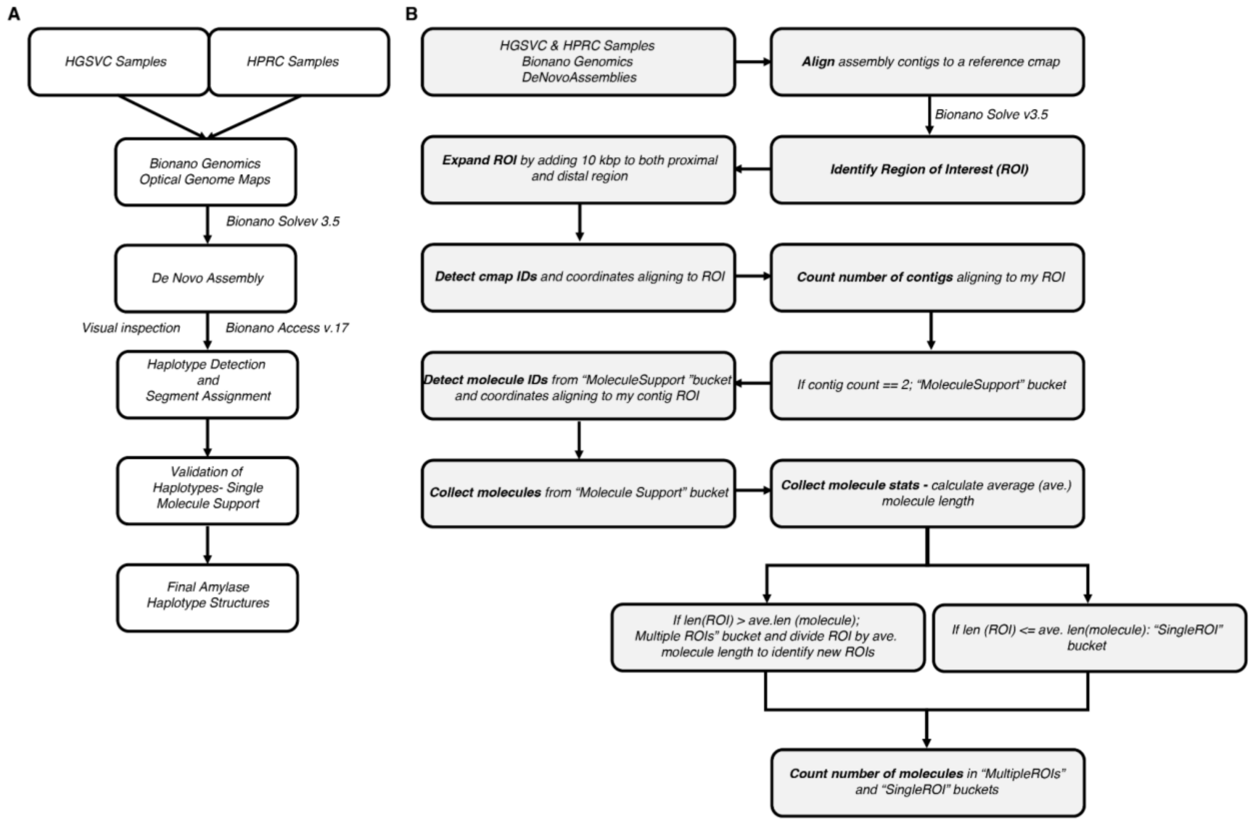

**Fig. S31. Analysis pipelines used to process HGSVC and HPRC datasets.**
(A) *De novo* assembly pipeline used for optical genome mapping datasets included in this study. (B) Optical genome mapping molecule support pipeline describing a step-by-step process to assess the molecule support for each sample. HGSVC: Human Genome Structural Variation Consortium, HPRC: Human Pangenome Reference Consortium.

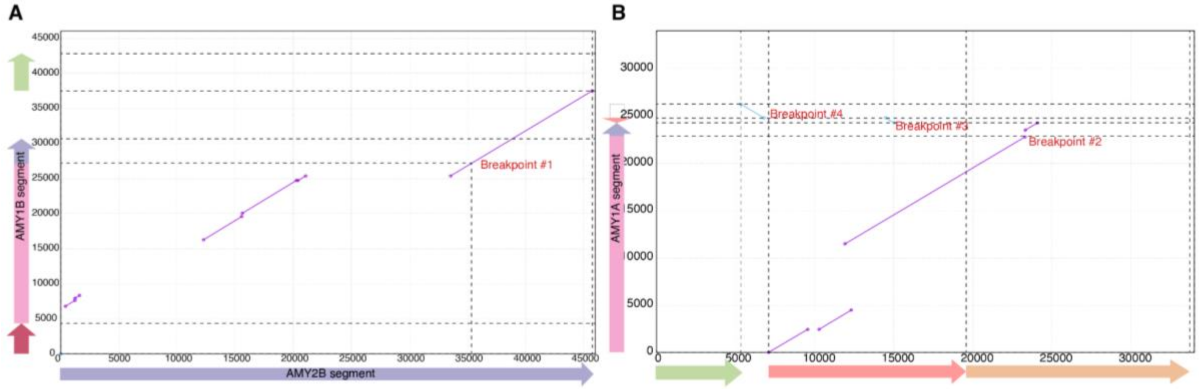


**Fig. S32. Rearrangement breakpoints detected in AMY1 segments in the context of structural haplotypes.**
(A) Dotplot showing the shared sequences (purple lines) between consensus sequences of the AMY2B segment and the AMY1B segment and its flanking region (y-axis: 30,000 - 45,000 bp). The AMY1B segment was annotated by sequence composition depicted in fig. S15. (B) Dotplot showing the shared sequences (purple and blue-inverted lines) between consensus sequences of AMY2A.1 segment and its flanking region, and the AMY1A segment. The AMY1A segment was annotated by sequence composition depicted in fig. S15.

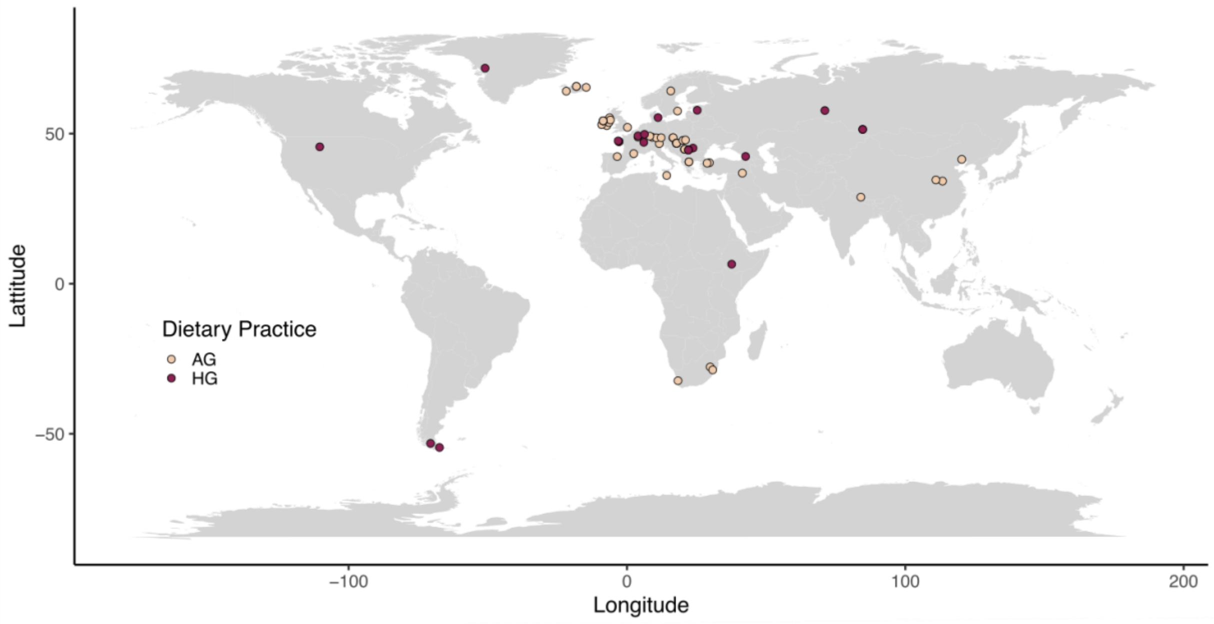


**Fig. S33. Map of the world for 73 ancient humans used in this study.**
Coloring is based on dietary habits with agriculturalists (AG) in beige and hunter-gatherers (HG) in maroon. The majority of the available samples are located in Europe.

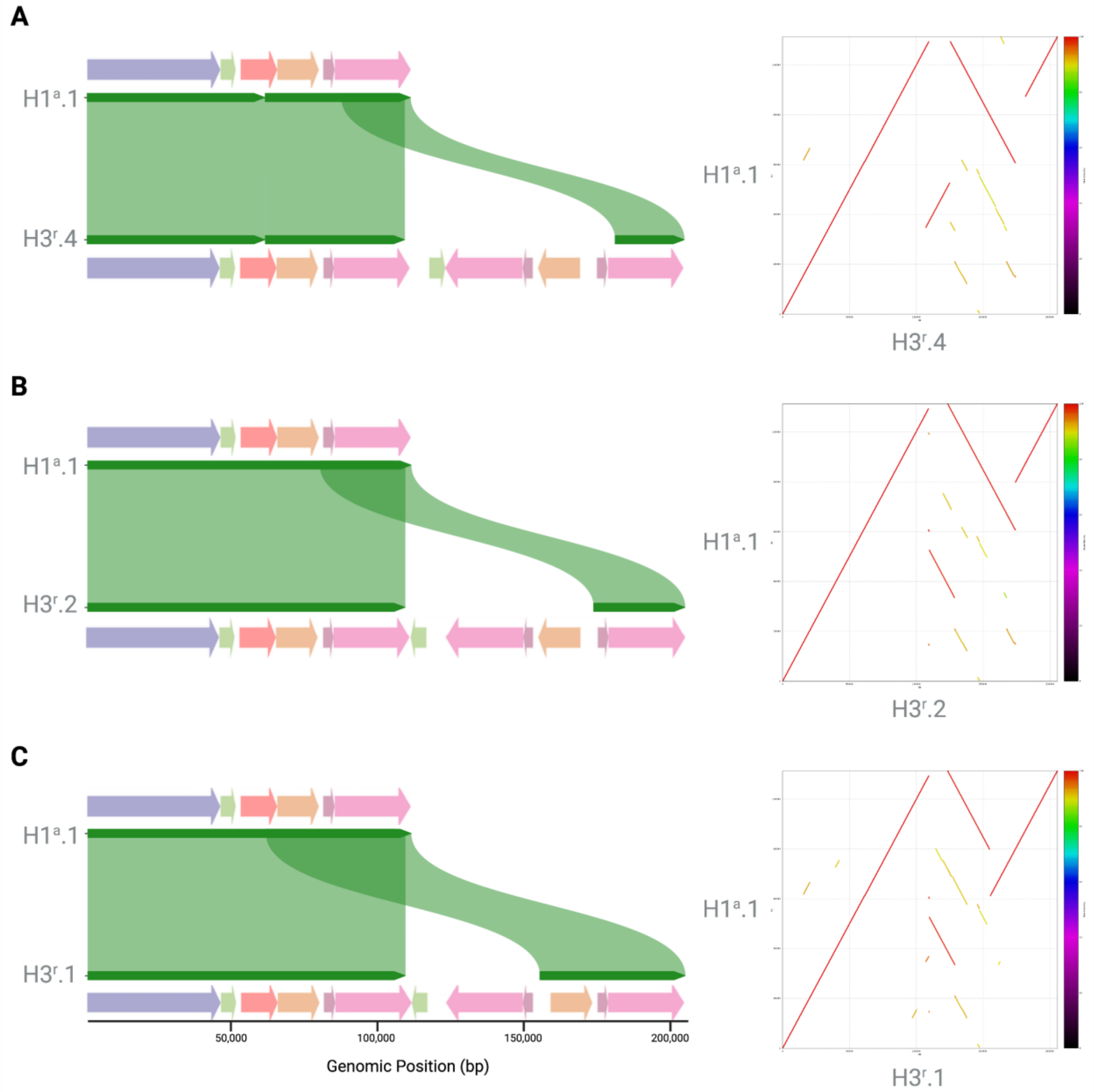


**Fig. S34. Comparison of the H1$^a$.1 haplotype to other common haplotypes.**
MIROPEAT (left) and dotplot (right) visualizations for the (A) H1$^a$.1 and H3$^r$.4, (B) H1$^a$.1 and H3$^r$.2, (C) H1$^a$.1 and H3$^r$.1 haplotypes. H1$^a$.1 differs from other common haplotypes by the absence of the AMY1A and AMY1B segments. These specific comparisons indicate that the emergence of the H1$^a$.1 haplotype was mediated by an NAHR-driven deletion between the two directly oriented AMY2A.2 segments, as represented by panel C, and can involve other three-copy haplotypes with similar directly oriented AMY2A.2 segments.

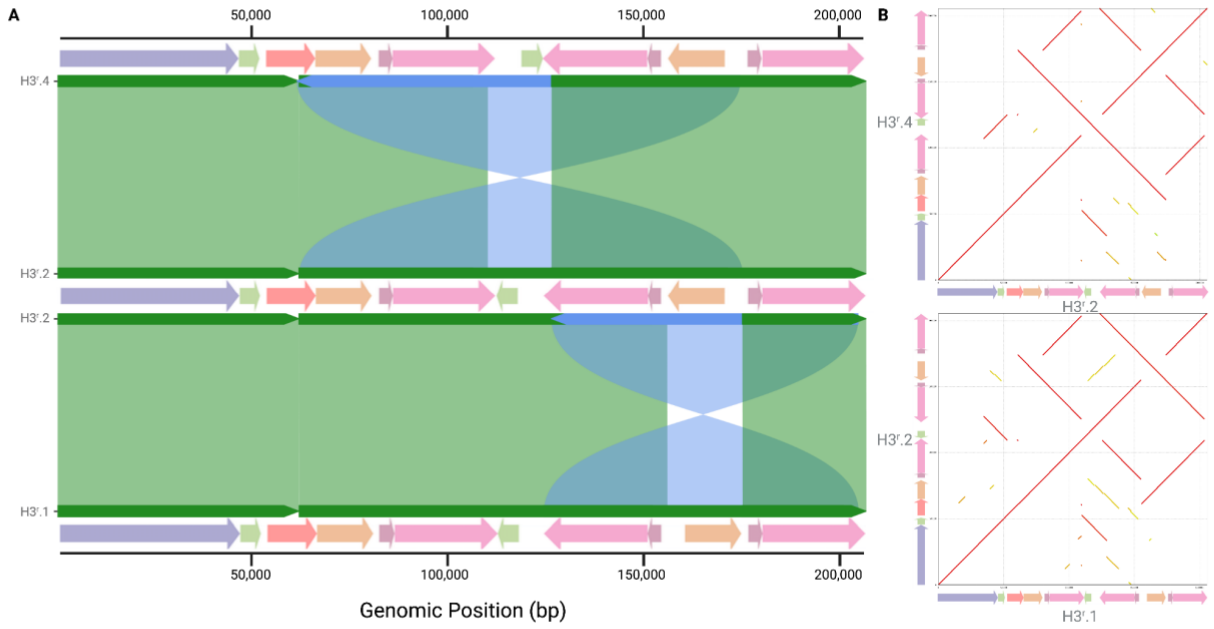


**Fig. S35. The comparison of the structural similarities among three common haplotypes, H3$^r$.1, H3$^r$.2, and H3$^r$.4, harboring the same copy number of amylase segments.** (A) MIROPEAT visualization in a pairwise manner allows the identification of two large inversions differentiating two haplotypes. (B) Dotplot visualization provides a more detailed look into the breakpoints of both inversions.

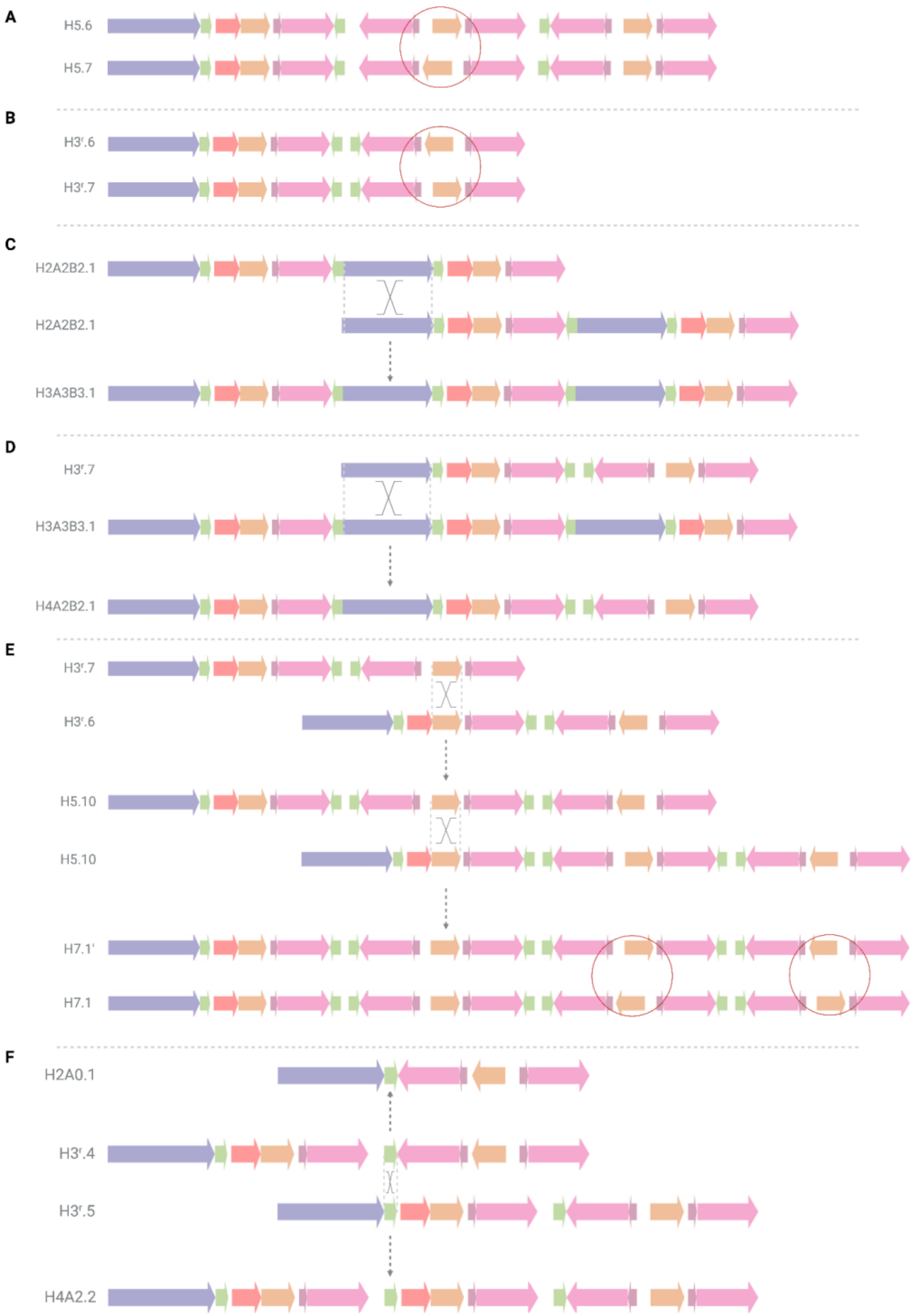


**Fig. S36. Inversions and copy number gains and losses via non-allelic homologous recombination events.**
(A) Potential NAHR event involving the inverted intergenic2 segment in the H5.6 haplotype that could lead to the inversion of the AMY2A.2 segment and thus the formation of the H5.7 haplotype. (B) Potential NAHR event involving the inverted intergenic.2 segment in the H3$^r$.6 haplotype that could lead to the inversion of the AMY2A.2 segment and thus the formation of the H3$^r$.7

haplotype. (C) The crossover between two H2A2B2.1 haplotypes resulting in the H3A3B3.1 haplotype and the observed copy number gains. (D) The crossover via NAHR between H2A2B2.1 and $H3^{r}$.7 haplotypes leading to copy number gains and generating the H4A2B2.1 haplotype. (E) The NAHR event between the forward-oriented AMY2A.2 segments of the $H3^{r}$.7 and $H3^{r}$.6 haplotypes produces the H5.10 haplotype (top). A second proposed NAHR event between two likely recently emerged H5.10 haplotypes could result in the formation of a hypothetical H7.1’ haplotype (middle). The two subsequent NAHR inversions involving the intergenic2 segments flanking the two distal AMY2A.2 segments could explain the formation of the H7.1 haplotype (bottom). (F) The NAHR-driven deletion may explain the emergence of the H2A0.1 haplotype from a crossover between the intergenic1 segments of the $H3^{r}$.4 and $H3^{r}$.5 haplotypes. Similarly, an unequal crossover between the intergenic1 segments of the aforementioned haplotypes could result in copy number gain and the formation of H4A2.2.

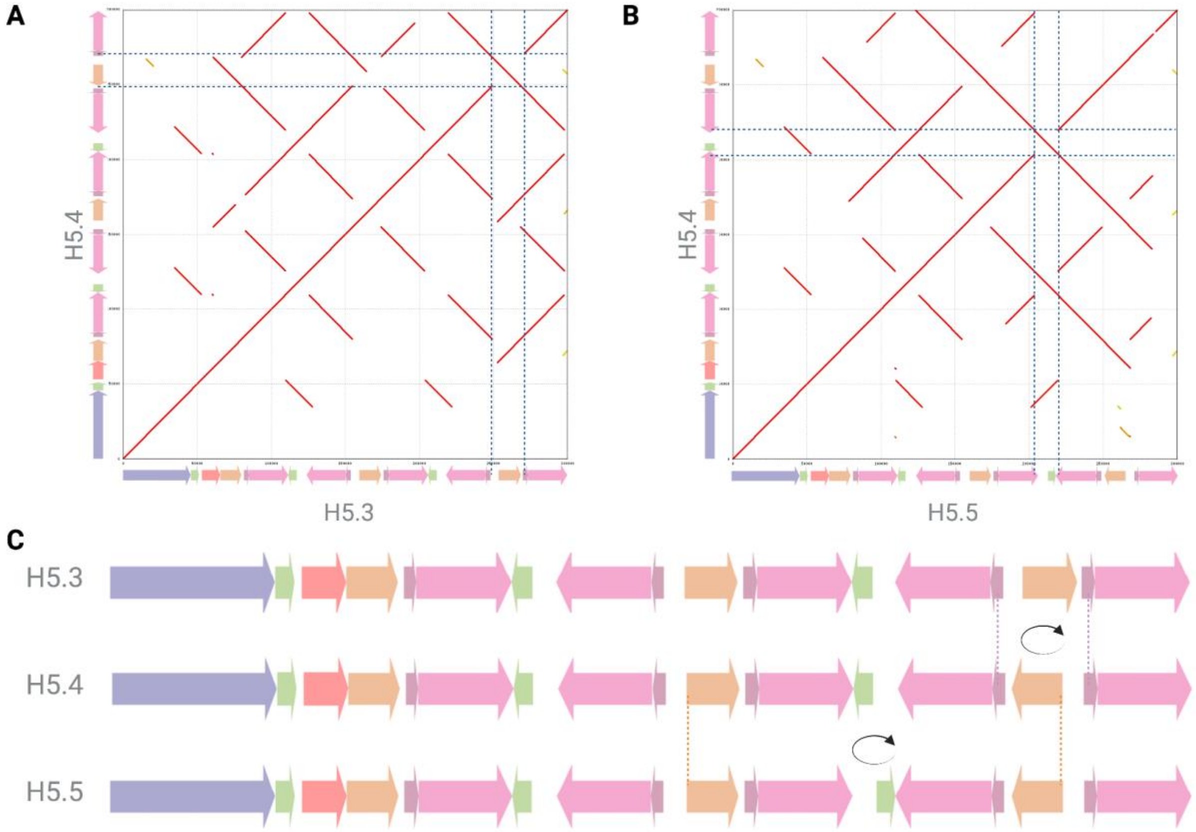


**Fig. S37. Structural differences among five-copy haplotypes.**
(A) Dotplot comparing H5.3 to H5.4 indicates a single NAHR-driven inversion, verified by local sequence alignments. The inversion spans the terminal AMY2A.2 segment and was likely mediated by the intergenic2 inverted repeats flanking the AMY2A.2 segment. The specific breakpoints are shown using dotted gray lines. (B) The predicted inversion breakpoints are highlighted in panel C. (C) Visual representation of the proposed stepwise NAHR-mediated inversions. The dotted lines indicate the breakpoints of inversions among these haplotypes. Note that H5.4 is an intermediate haplotype between H5.3 and H5.5 when inverted segments are considered.

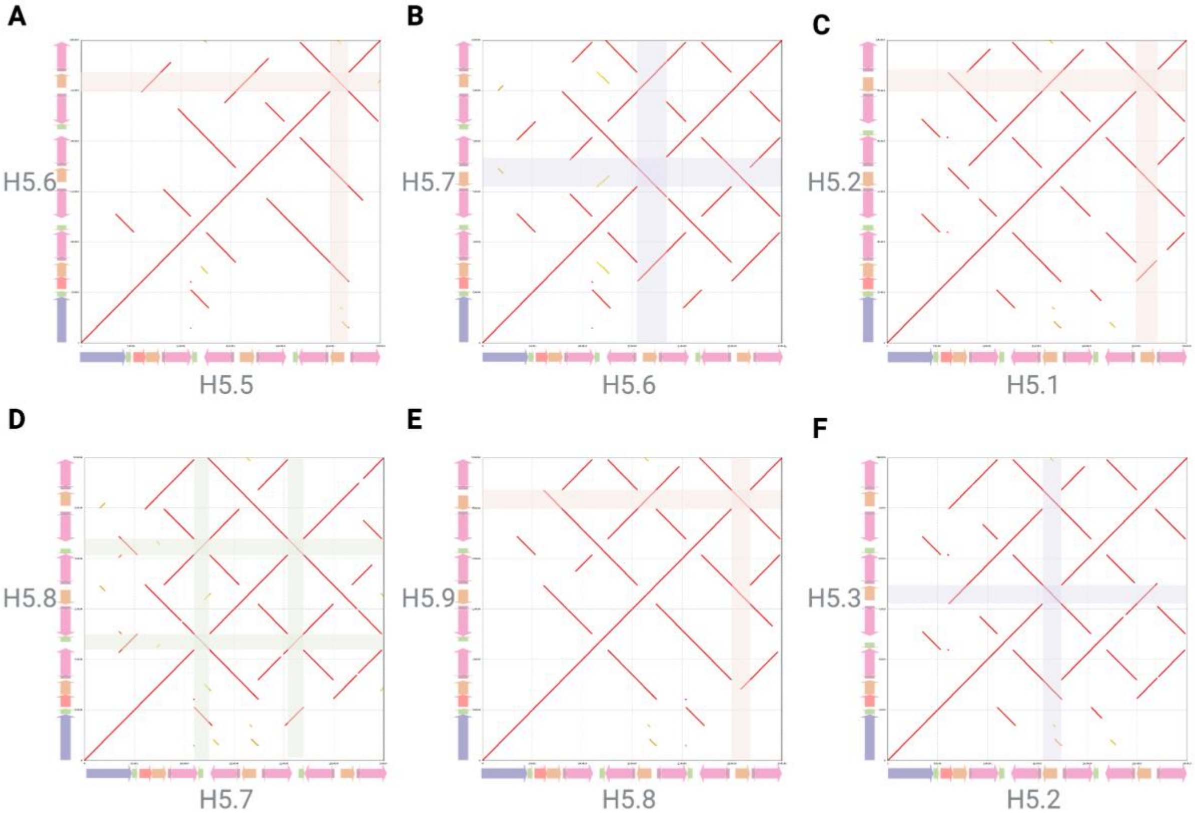


**Fig. S38. Recurrent inversions among intermediate and rare five-copy haplotypes.**
(A) Comparison of H5.5 to H5.6 indicates that the two haplotypes are separated by a single inversion of the terminal AMY2A.2 segment mediated by the two flanking intergenic2 segments. (B) Comparison of H5.6 to H5.7 haplotype indicates a single NAHR inversion of the second AMY2A.2 segment. The inversion is mediated by the flanking intergenic.2 inverted repeats. (C) Comparison of H5.1 to H5.2 reveals a single inversion of the terminal AMY2A.2 segment mediated by the two flanking intergenic2 segments. (D) Comparison of H5.7 to H5.8 haplotypes reveal two complex NAHR inversion events, spanning the proximal intergenic1 segment and the distal intergenic1 segment. The first inversion was mediated by the two inverted AMY2A.2 segments, spanning the first AMY1A segment to the first AMY1B segment. Similarly, the second inversion was mediated by the two inverted AMY2A.2 segments spanning the second AMY1A segment to the second AMY1B segment. (E) Comparison of H5.8 to H5.9. The two haplotypes are separated by a single inversion of the terminal AMY2A.2 segment mediated by the two flanking intergenic.2 segments. (F) A comparison of the H5.2 to H5.3 haplotype revealed a single NAHR inversion of the second AMY2A.2 segment. The NAHR inversion is mediated by the flanking intergenic.2 inverted repeats.

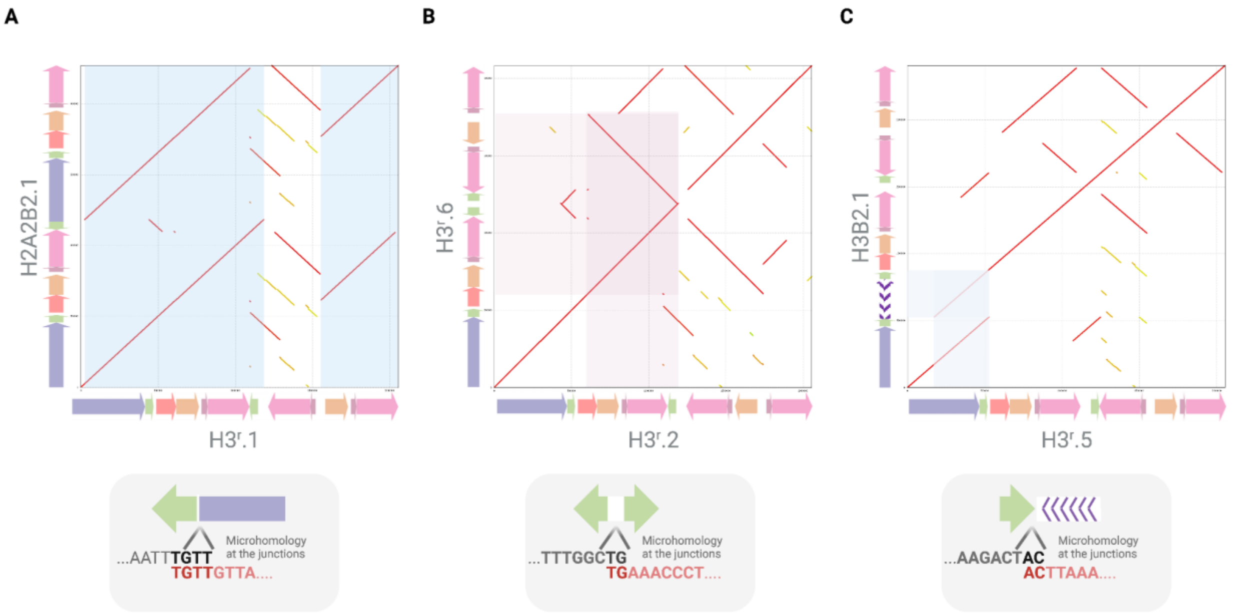


**Fig. S39. Microhomology-mediated break-induced replication events.**
Events explaining the generation of haplotypes that harbor *AMY2* gene duplications among others. (A) Comparison of the H2A2B2.1 haplotype to the H3$^{r}$.1 haplotype. The microhomology-mediated break-induced replication (MMBIR) results in the gain of an additional copy of the AMY2B segment. Four nucleotides of microhomology internal to the breakends were identified at the breakpoint junction as indicated below the dotplot. (B) Comparison of H3$^{r}$.6 to H3$^{r}$.2. The H3$^{r}$.6 haplotype has gained an additional copy of the intergenic.1 and AMY1A segments inserted with the reverse orientation, while losing the AMY1B segment. Two nucleotides of microhomology internal to the breakends were identified at the breakpoint junction and indicated below the dotplot. (C) Comparison of H3B2.1 to H3$^{r}$.5 displaying that the H3B2.1 haplotype has a partial copy of the AMY2B segment. This MMBIR- based event is facilitated by the two-nucleotide microhomology internal to the breakends shown below the dotplot.

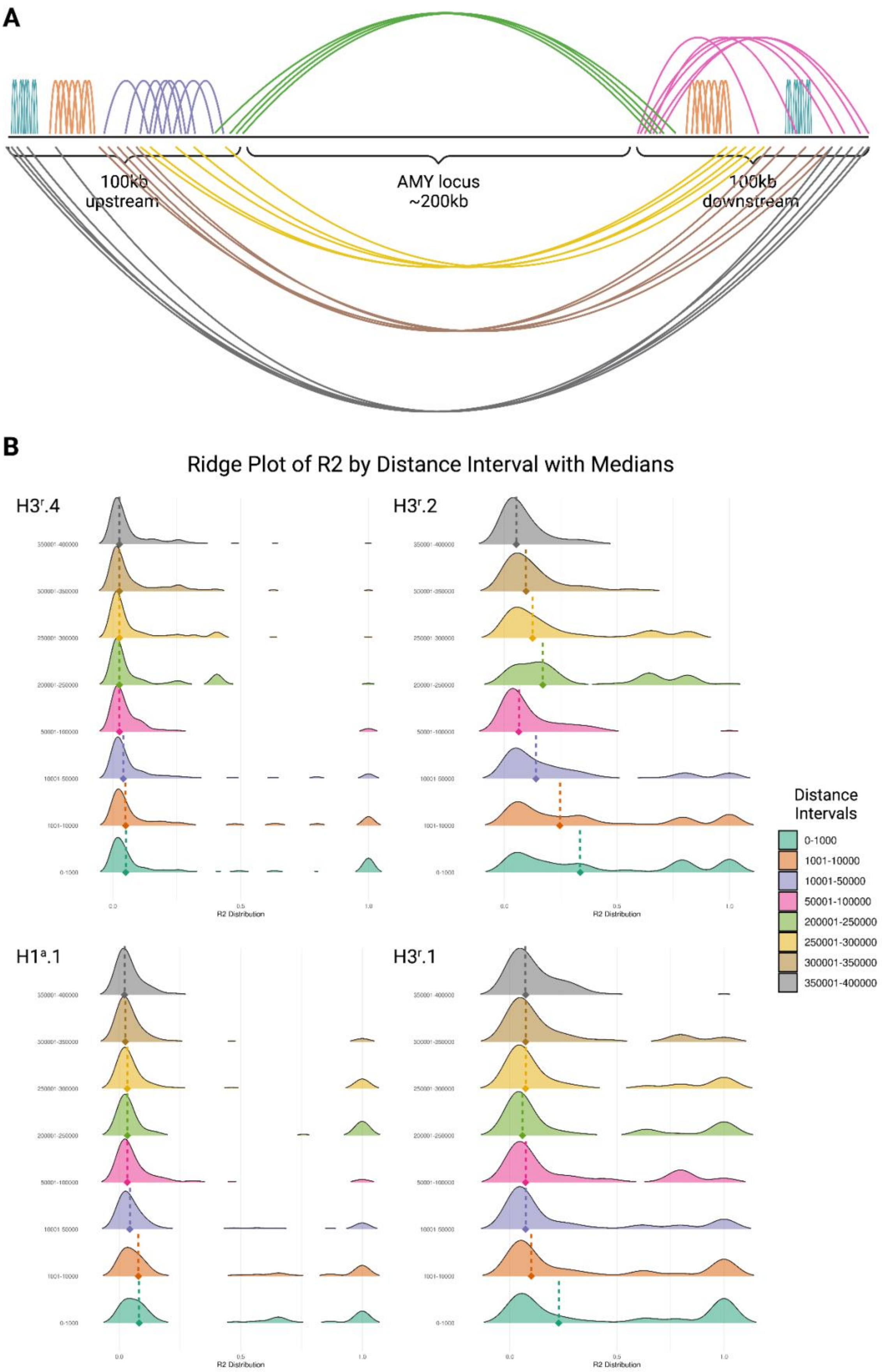
A
100kb
upstream
AMY locus
~200kb
100kb
downstream
B
Ridge Plot of R2 by Distance Interval with Medians
H3r.4
H3r.2
H1a.1
H3r.1
R2 Distribution
Distance
Intervals
0-1000
1001-10000
10001-50000
50001-100000
200001-250000
250001-300000
300001-350000
350001-400000

**Fig. S40. Ridge plot of $R^2$ by distance interval with medians for the four common haplotypes.** (A) Each color arc represents instances of LD estimates for a different distance interval. Each distance interval (*e.g.*, 0-1,000 bp in teal or 50,001-100,000 bp in pink) aggregates data from multiple combinations of pairs of alleles, also spanning across the upstream and downstream amylase region. (B) The distribution of linkage disequilibrium (LD) as measured by $R^2$ across various distance intervals for the four common haplotypes. Each plot corresponds to one haplotype and is divided into ten distance intervals (0-1,000 bp, 1,001-10,000 bp, 10,001-50,000 bp, and intervals of 50,000 bp beyond the 10,001 bp threshold). The intervals between 100,001-150,000 bp and 150,001-200,000 bp remain empty because we were investigating the upstream and downstream flanking regions, and excluding the amylase locus (see **Supplementary Methods**). The $R^2$ distribution within each distance interval is visualized as a ridge, with the height of the ridge representing the density of data points at different $R^2$ values. The colors from light green to dark gray represent progressively larger distance intervals. The medians of these distributions are marked by a diamond and a vertical dotted line within each ridge.

### Data Files (separate files)

DataFile-S1-OrthogonalValidation_MoleculeSupport.pdf: doi: 10.6084/m9.figshare.24645912
DataFile-S2-AmylaseAlignments.zip
